# RADICL-seq identifies general and cell type-specific principles of genome-wide RNA-chromatin interactions

**DOI:** 10.1101/681924

**Authors:** Alessandro Bonetti, Federico Agostini, Ana Maria Suzuki, Kosuke Hashimoto, Giovanni Pascarella, Juliette Gimenez, Leonie Roos, Alex J. Nash, Marco Ghilotti, Christopher JF Cameron, Matthew Valentine, Yulia A Medvedeva, Shuhei Noguchi, Eneritz Agirre, Kaori Kashi, Samudyata, Joachim Luginbuehl, Riccardo Cazzoli, Saumya Agrawal, Nicholas M Luscombe, Mathieu Blanchette, Takeya Kasukawa, Michiel De Hoon, Erik Arner, Boris Lenhard, Charles Plessy, Gonçalo Castelo-Branco, Valerio Orlando, Piero Carninci

## Abstract

Mammalian genomes encode tens of thousands of noncoding RNAs. Most noncoding transcripts exhibit nuclear localization and several have been shown to play a role in the regulation of gene expression and chromatin remodelling. To investigate the function of such RNAs, methods to massively map the genomic interacting sites of multiple transcripts have been developed. However, they still present some limitations. Here, we introduce <u>R</u>NA <u>A</u>nd <u>D</u>NA Interacting <u>C</u>omplexes <u>L</u>igated and <u>seq</u>uenced (RADICL-seq), a technology that maps genome-wide RNA-chromatin interactions in intact nuclei. RADICL-seq is a proximity ligation-based methodology that reduces the bias for nascent transcription, while increasing genomic coverage and unique mapping rate efficiency compared to existing methods. RADICL-seq identifies distinct patterns of genome occupancy for different classes of transcripts as well as cell type-specific RNA-chromatin interactions, and emphasizes the role of transcription in the establishment of chromatin structure.

## Introduction

The vast majority of mammalian genomes is pervasively transcribed, accounting for a previously unappreciated complexity of the non-coding RNA (ncRNA) fraction^1^. In particular, long noncoding RNAs (lncRNAs) have emerged as important regulators of various biological processes^2^. Although most lncRNAs exhibit nuclear localization with an enrichment for the chromatin fraction, the genomic interacting regions for the majority of these transcripts are still unknown^2, 3^.

Several technologies have been developed to map the genomic interacting sites of lncRNAs^4–6^. However, these methodologies rely on the use of antisense probes to target individual transcripts, and are not suitable for *de novo* discovery and high throughput application in multiple cell types.

A few technologies have emerged to assess genome-wide RNA-chromatin interactions^7–9^ which nevertheless present limitations. Mapping RNA-Genome Interactions (MARGI) is a proximity ligation-based technology that requires a high number of input cells (i.e. hundreds of millions) and the disruption of the nuclear structure^7^, which can result in detection of a large number of spurious interactions^10^ and therefore offers a limited pan-cellular application to investigate cell type-specific RNA-chromatin interactions. Chromatin associated RNA sequencing (CHAR-seq) and global RNA Interaction with DNA by deep-sequencing (GRID-seq) utilize *in situ* approaches to detect genome-wide RNA-chromatin contacts^8, 9^. CHAR-seq requires a large number of cells as starting material and the use of Dpn II to digest the chromatin, which has a restricted number of sites across the genome and, hence, a lower coverage for captured RNA-DNA interactions^9^. In addition, the technology does not size-select the molecules containing both interacting RNA and DNA, resulting in a large fraction of uninformative sequences in the final library. GRID-seq preferentially captures nascent RNA-chromatin interactions and, consequently, may overlook the presence of other patterns of genome occupancy by specific classes of transcripts. Furthermore, reliance on a restriction enzyme for chromatin fragmentation coupled with read length limited to 20 nucleotides (nts) reduces both genome coverage and mappability.

To address these limitations, we have developed **<u>R</u>**NA **<u>A</u>**nd **<u>D</u>**NA **<u>I</u>**nteracting **<u>C</u>**omplexes **<u>L</u>**igated and **<u>seq</u>**uenced (RADICL-seq), a methodology to identify genome-wide RNA-chromatin interactions in crosslinked nuclei that significantly improves on previously published methods. Specifically, RADICL-seq reduces the bias for nascent transcription, while increasing genomic coverage and unique mapping rate efficiency. Application of RADICL-seq to mouse embryonic stem cells (mESCs) and mouse oligodendrocyte progenitor cells (mOPCs) reveals distinct genome occupancy patterns for specific classes of transcripts and uncovers cell-type specific RNA-chromatin interactions. Furthermore, our results highlight the role of transcription in the establishment of the three-dimensional (3D) structure of chromatin.

## Results

### RADICL-seq technology

We developed RADICL-seq by using R08, a *Mus musculus* male embryonic stem cell line with a deeply characterized transcriptome^11^, to identify genome-wide RNA-chromatin (or RNA-DNA) interactions in preserved nuclei (**Fig. 1a**). We crosslinked cells with two different formaldehyde (FA) concentrations (1% and 2%) to test whether captured interactions were dependent on the amount of crosslinking agent. After crosslinking we isolated the nuclei, partially digested the genomic DNA with DNase I and ends-prepared the chromatin. During technical development of RADICL-seq we evaluated different enzymes that specifically act on RNA to generate a 3’-hydroxyl end, compatible with RNA ligation (**Supplementary Fig. 1a**). Sequencing data of test RADICL-seq libraries showed that RNase H treatment increased the percentage of uniquely mapped RNA-chromatin interactions by decreasing the ribosomal RNA (rRNA) content, when compared to nuclease S1, RNase V1 or absence of treatment. RNase H enzymatic treatment is known to target RNA-DNA hybrids and, therefore, it could potentially digest nascent RNA bound to its transcription locus, including the highly transcribed rRNA. Indeed, we observed a 2.5-fold reduction in the number of RNA-DNA interactions occurring at a distance below 1 kb between RNAse H-treated and untreated samples (**Supplementary Fig. 1b**). After enzymatic treatment of the RNA, we introduced a bridge adapter to specifically ligate proximal RNA and DNA (**Supplementary Fig. 1c**). The adapter is a 5’ pre-adenylated, partially double stranded DNA linker with an internal biotin moiety and a thymidine (T) overhang located at the 3’end. The adapter was selectively ligated to available 3’-OH RNA ends and the excess of non-ligated adapter was washed away before performing DNA ligation to capture the digested genomic DNA ends located in proximity (**Fig. 1a**). The experimental design of RADICL-seq not only allows for unambiguous discrimination of RNA and DNA tags within the chimeric construct but also correctly assigns sense and antisense transcripts by retaining the information on the RNA fragment strand.

**Figure 1.**
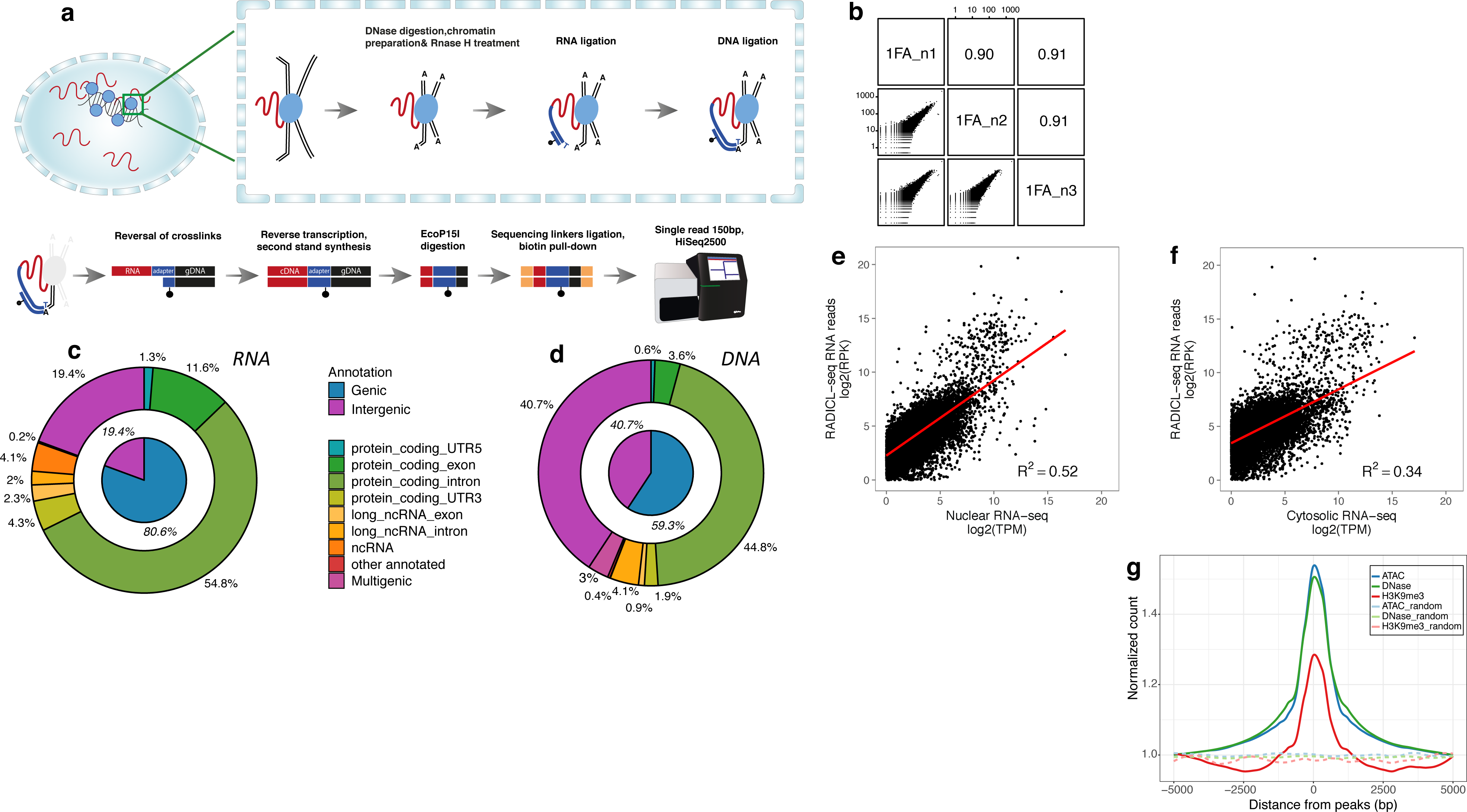
RADICL-seq method for the identification of RNA-chromatin interactions. a) Schematic representation of the RADICL-seq protocol. Top: sequence of enzymatic reactions taking place in fixed nuclei after partial lysis of the nuclear membrane. The adduct formed by genomic DNA (black), RNA (red) and proteins (blue circles) goes through controlled DNase digestion and chromatin preparation. After RNase H digestion, an adapter (dark blue) containing an internally biotinylated residue (black dot) bridges RNA and DNA in close proximity. Bottom: sequence of enzymatic reactions performed in solution. After reversal of crosslinks, the RNA-DNA chimera is converted into a fully dsDNA molecule and digested by EcoP15I enzyme to a designated length. After ligation of the sequencing linker and biotin pull-down, the library is PCR-amplified and high-throughput sequenced. b) Reproducibility of the RNA-DNA interaction frequencies across replicates, assessed by counting the occurrences of transcribed genes and 25 kb genomic bins pairs. c) RNA and d) DNA tags origin. The inner pie charts represent a broader classification into intergenic and genic (annotated genes), while the outer circles show a finer classification of the genic portion. e) Nuclear and f) cytosolic RNA-seq tags comparison with RADICL-seq RNA reads gene counts. The former are normalized to tags per million (TPM), while the latter are normalised to reads per kilobase (RPK). The linear regression lines are shown in red. g) Density of the normalized counts of DNA reads detected by RADICL-seq around ATAC-seq (red), DHS-seq (green) and H3K9me3 ChIP-seq (blue) peaks; dashed lines represent the density profiles of random genomic reads.

After reversal of crosslinks, the resulting RNA-adapter-DNA chimera was converted to dsDNA by reverse transcription and second strand DNA synthesis, followed by digestion with type III restriction enzyme EcoP15I which cleaves 25 to 27 nts away from each of its two recognition sites strategically placed within the adapter (**Supplementary Fig. 1c**). Next, the digested DNA fragments were ends-prepared and ligated to sequencing linkers. Finally, the biotinylated adapter-ligated molecules were captured, PCR-amplified and the library corresponding to the correct RNA-adapter-DNA ligation product size was gel-purified (**Supplementary Fig. 1d**).

After deep sequencing, RADICL-seq produced, on average, 120 and 115 million 150 nt single-end raw reads from libraries crosslinked with 1% and 2% FA, respectively (**Supplementary Fig. 2a**). Each library with the two different crosslinking conditions yielded over 15 million RNA-DNA pairs uniquely mapping to the reference genome (**Supplementary Fig. 2a**). We prepared libraries from three biological replicates for each experimental condition. RADICL-seq exhibited high reproducibility among biological replicates and conditions, even when crosslinked with different formaldehyde concentrations. (**Fig. 1b** and **Supplementary Fig. 2b**). Since results obtained with 1% and 2% formaldehyde crosslinking were highly comparable, we decided to focus on 1% FA data for downstream analyses.

To characterize the interactions detected by RADICL-seq, we annotated RNA-DNA pairs that could be uniquely mapped to the genome. The RNA tags were found to be primarily from genic regions with a dominant contribution from intronic reads (**Fig. 1c**). In contrast, DNA tags had an equivalent contribution from genic (mainly intronic) and intergenic regions (**Fig. 1d**).

We assigned the RNA and DNA fragments captured by RADICL-seq to the genomic features annotated by the GENCODE consortium^12^ and analyzed their distribution among different classes of gene biotypes (**Supplementary Fig. 2c**). Protein-coding genes were the most abundant class of loci detected by RADICL-seq at both RNA and DNA level. Indeed, we observed multiple classes of transcripts interacting with chromatin regions encompassing protein-coding genes, suggesting a multi-layered regulation for the expression of these mRNAs.

When the expression of chromatin-interacting RNAs was compared with fractionated RNA-seq data^11^, a significantly higher correlation was found with the nuclear fraction than the cytosolic counterpart (two-sided Wilcoxon-Mann-Whitney test; p-value <10^−115^, **Fig. 1e,f**). This finding is in line with RADICL-seq capturing ligation events occurring between RNAs and DNAs located within intact nuclei.

Although the majority of DNA reads captured by RADICL-seq originates from euchromatin (based on DHS- and ATAC-seq data), we observed a minor enrichment from genomic regions located in heterochromatic regions, consistent with the role of some lncRNAs as repressors of gene expression^13^ (**Fig. 1g**).

To better evaluate the quality of our results, we developed two controls (**Supplementary Fig. 3a**). In order to test the stability of RNA-chromatin interactions upon transcriptional blockade we treated mESCs with Actinomycin D (ActD), an inhibitor of RNA polymerase II (RNA pol II) elongation^14^, for 4 hours before crosslinking (**Supplementary Fig. 3b,c**). The second control was developed to estimate the specificity of RNA-chromatin interactions mediated by the presence of proteins. To this end, crosslinks were reversed immediately before the RNA ligation reaction by digesting the sample with proteinase K in denaturing conditions. As a result, RNA and DNA would be able to reproducibly interact only if the binding was direct and not mediated by the presence of proteins. We defined this dataset as “non-protein-mediated” (NPM) (**Supplementary Fig. 3d,e),** as opposed to the standard experimental condition where crosslinks were maintained throughout the ligation steps (here defined as “total” and including protein-mediated as well as non-protein-mediated interactions). As expected, comparison of total, ActD-treated and NPM datasets displayed qualitative and quantitative differences in RNA-chromatin interactions across experimental conditions (**Supplementary Fig. 4**).

### Comparison of RADICL-seq with existing technologies that target RNA-chromatin interactions

RADICL-seq introduces substantial improvements over similar RNA-chromatin proximity ligation approaches^7,8,9^. Compared to MARGI, RADICL-seq minimizes the frequency of spurious interactions in the dataset by performing the *in situ* ligation in intact nuclei. Moreover, RADICL-seq requires a significantly lower amount of cells (2 million) compared to MARGI and CHAR-seq that require, respectively, 400 and 100 million cells.

Compared to GRID-seq, RADICL-seq differs by four main technical aspects (**Fig. 2a**): i) GRID-seq employs in the fixation step a higher concentration of formaldehyde and disuccinimidyl glutarate (DSG), a strong protein-protein crosslinker, thus capturing RNA and DNA linked together indirectly via multiple protein intermediates^15^; ii) the GRID-seq protocol employs the type II restriction enzyme MmeI to trim RNA and DNA interacting sequences, generating 20 nt tags as opposed to the 27 nt tags produced by the EcoP15I restriction enzyme, thus resulting in tags that are more difficult to map uniquely to the genome. Indeed, when we compared the percentage of sequencing reads that can be mapped to the mouse genome, RADICL-seq outperformed GRID-seq with more than a three-fold increase in uniquely mappable reads (45% vs 14%) (**Fig. 2b**). To confirm this result, we artificially trimmed down RADICL-seq tags to generate RNA and DNA reads with lengths similar to those obtained by GRID-seq tags (i.e., 20 nt) and observed a dramatic reduction in the fraction of uniquely mapped RNA-DNA interactions (from 45% to 10%), comparably to the rate observed for the GRID-seq dataset and in concordance with previous findings^16^. Furthermore, reducing RNA and DNA read lengths from 27 to 20 nt resulted in diminished frequency of uniquely mapped DNA reads irrespective of the RNA reads mapping quality. This difference affects both the number and the type of detected interactions since reads encompassing repetitive regions are intrinsically more difficult to map compared to those from other genomic regions.

**Figure 2.**
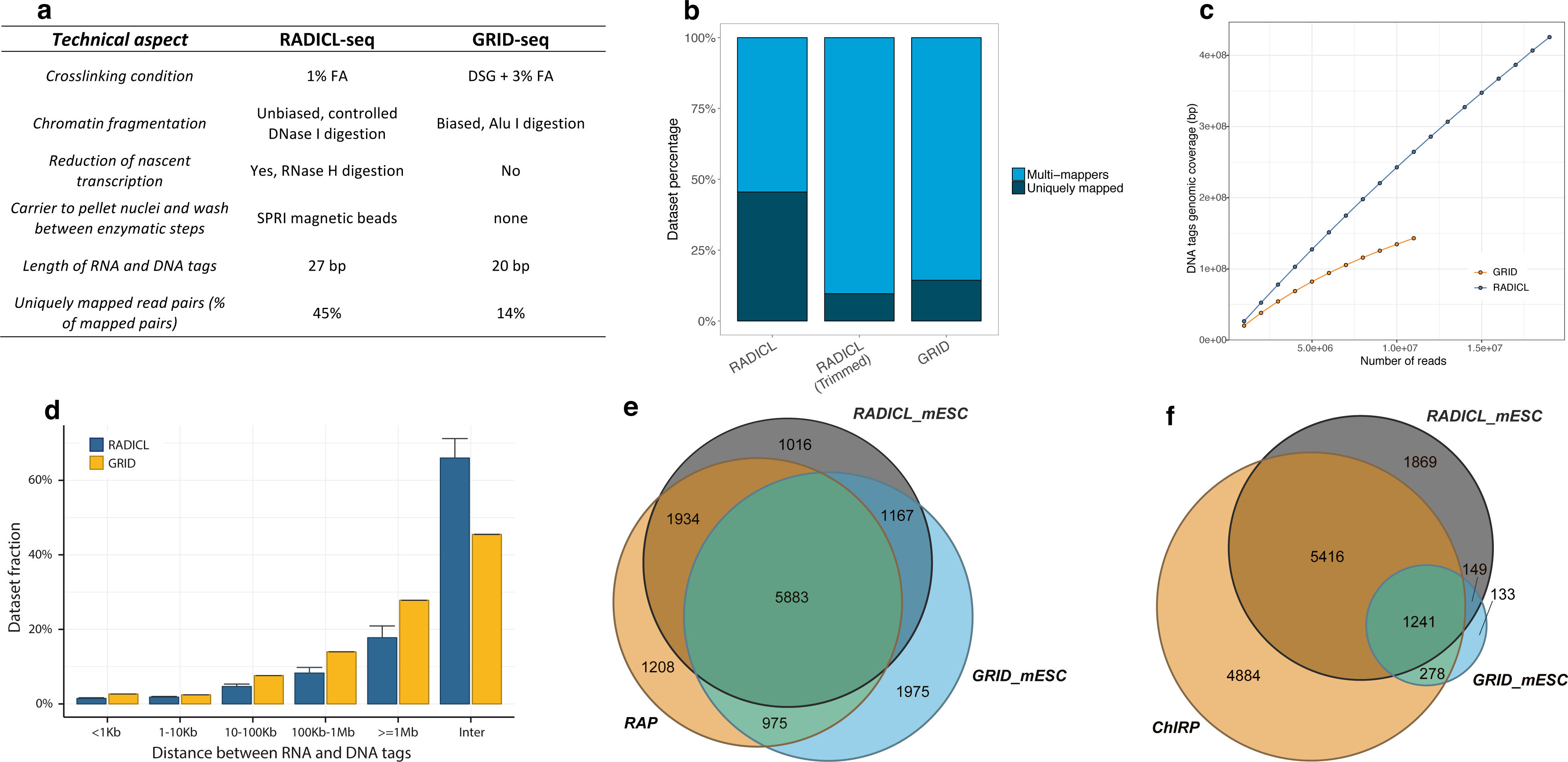
RADICL-seq comparison with similar methods. a) Summary of the features that distinguish RADICL-seq from GRID-seq. b) Analysis of the read length and mapping outcome. Unique (dark grey) and multi-mapping (blue) reads are reported as percentage of the reads pool. RADICL-seq reads were artificially trimmed down to 20nt for direct comparison with the GRID-seq dataset. c) Assessment of the genomic coverage as a function of the sequencing depth for RADICL-seq (blue) and GRID-seq (orange). The coverage was calculated for both datasets by sub-sampling with a step of 1’000’000 reads up to the maximum available number of reads. d) Distribution of the linear genomic distance between RNA and DNA tags derived from the same read for the GRID-seq (yellow) and RADICL-seq (blue) datasets. e) Comparison of Malat1 transcript target DNA loci identified by RAP (yellow), RADICL-seq (grey) and GRID-seq (blue) methods. f) Comparison of Rn7sk transcript target DNA loci identified by ChIRP (yellow), RADICL-seq (grey) and GRID-seq (blue) methods.

iii) GRID-seq technology digests genomic DNA with AluI, whereas the RADICL-seq protocol employs a controlled DNase I digestion that avoids sequence biases as encountered with restriction enzymes and, therefore, generates a more homogenous shearing of the chromatin. When we looked at the genomic coverage of DNA regions identified by both technologies, RADICL-seq exhibited a higher coverage (**Fig. 2c**). We additionally observed that RADICL-seq genome coverage increased proportionally with the sequencing depth whereas the coverage of GRID-seq converged to a plateau (**Fig. 2c**)

Finally, iv) RADICL-seq employs RNase H treatment prior to the RNA ligation step to reduce the number of captured interactions generated by nascent transcription and consequently increases the variety of captured RNA-DNA interactions. When compared to uniquely mapped RNA and DNA reads detected by GRID-seq, the RADICL-seq dataset showed increased detection of intergenic transcripts and ncRNAs (**Supplementary Fig. 5a**). Moreover, in GRID-seq data we observed both a higher contribution of intronic coding RNA reads (66.9% vs 54.7%) and a 2.5 fold-change in RNA-DNA interactions occurring at a distance below 1 kb (**Supplementary Figure 5a**, **Fig. 2d**) that suggests higher content of nascent transcripts in this set of interactions. Furthermore, GRID-seq DNA reads exhibit stronger enrichment for H3K36me3 signal, a marker of elongating RNA pol II, suggesting a stronger bias for nascent transcription when compared to RADICL-seq (**Supplementary Fig. 5b**). In contrast, RADICL-seq captures RNA sequences derived from the body of annotated genes and enriched for histone modifications associated with exonic regions (H3K4me3 and H3K9Ac), indicating enrichment for mature transcripts^17^ (**Supplementary Fig. 5c,d**). Interestingly, while GRID-seq displayed a larger fraction of captured *cis* (*i.e.*, intra-chromosomal) interactions, RADICL-seq recovered ∼30% more *trans* (*i.e.*, inter-chromosomal) interactions, thus providing a vastly expanded interactions dataset to investigate long-range RNA-chromatin associations (**Fig. 2d**). We overall observed lower correlation between technologies than within replicates, suggesting RADICL-seq and GRID-seq capture different sets of interactions (**Supplementary Fig. 5e**).

We evaluated the extent of known interactions captured by RADICL-seq and GRID-seq by comparing the genomic targets of Malat1 and Rn7sk detected by RADICL-seq with those observed using respectively RNA antisense purification followed by DNA-sequencing (RAP-DNA)^15^ and chromatin isolation by RNA purification (ChIRP-seq)^18^. RADICL-seq data had a genomic distribution comparable to that of Malat1-targeted RAP-DNA libraries prepared from pSM33 ES cells^15^ (**Supplementary Fig. 5f**). The RADICL-seq list of targets confirmed 78% of Malat1-decorated protein-coding genes in the RAP-DNA library compared to 71% of targets found by GRID-seq (**Fig. 2e**). When compared to GRID-seq, RADICL-seq exhibited a lower percentage of genomic targets not detected by RAP-DNA. We then turned our attention to the genomic targets of Rn7sk detected by ChIRP-seq^18^. In this case, we observed a clear difference in the capture rate of noncoding transcripts between GRID-seq and RADICL-seq, with the latter comprising over twenty-fold more interactions mediated by Rn7sk (0.357 and 9.491 average normalized counts in GRID-seq and RADICL-seq, respectively). Furthermore, RADICL-seq detected 56% of the protein-coding genes interacting with Rn7sk compared to only 13% of targets for GRID-seq (**Fig. 2f**).

Collectively, these analyses suggest that RADICL-seq is able to capture RNA-chromatin interactions more comprehensively and with less genomic bias when compared to GRID-seq.

### Identification of robust RNA-chromatin interactions in distinct cell types

The RADICL-seq technology yields a large amount of interactions data with a complexity comparable to that obtained with the Hi-C technology. Consequently, to account for the occurrence of spurious events, we decided to adopt an approach similar to those employed in Hi-C analyses^19–21^, which assumes that all biases (e.g., amplification biases due to differences in sequence composition across the genome) are reflected in the observed interaction counts. To this end, we partitioned the linear genome into intervals (*i.e.*, bins of 25 kb, see Methods) to represent the RADICL-seq data as a “contact matrix” between RNA and DNA loci. We then used a one-sided cumulative binomial test to detect significant RNA-chromatin interactions, assuming that the transcript-specific background interaction frequency of a given RNA and a genomic interval depends also on their relative genome-wide coverage^22^. We employed the Benjamini–Hochberg multiple-testing correction to control for the false discovery rate (FDR) and used a cut-off of 0.05 to define the “significant” set (**Supplementary Fig. 6a**). By applying this method to RADICL-seq data, 288,068 unique robust RNA-DNA interacting loci, supported by 7,909,964 interactions, were identified as statistically significant (**Supplementary Fig. 6b**). These RNA-chromatin interactions were mediated by 14,122 transcripts, with a prevalent contribution from mRNAs (88%) followed by lncRNAs (10%, around 1,400 transcripts) (**Supplementary Fig. 6c**). As expected, many of the *trans*-interactions were removed because of their lower occurrence and inherent difficulty in being consistently detected at the chosen sequencing depth (**Supplementary Figure 6b**).

Furthermore, to compare RNA-chromatin interactions patterns across different cell types, we performed RADICL-seq on oli-neu, a neural cell line derived from mouse oligodendrocyte progenitor cells (mOPCs)^23^ (**Supplementary Fig. 7a**). Again, the three biological replicates exhibited high reproducibility, but markedly lower correlation with the mESCs biological replicates, suggesting cell specificity in a substantial fraction of the captured interactions (**Supplementary Fig. 7b**).

Interestingly, the comparison of the non-coding transcriptome captured by RADICL-seq in mESCs and mOPCs highlighted a marked increase in the detection of lncRNAs in the mOPCs dataset (**Supplementary Fig. 7c,d**). Although mapping of RNA and DNA tags from the NPM revealed a similar pattern as the total dataset (**Supplementary Fig. 8a-c**), comparison of the interaction counts in the two dataset displayed higher variability (**Supplementary Fig 8d**).

To globally visualize the RNA-DNA interactions, we arranged the transcripts and their interacting genomic regions in a two-dimensional contact matrix, where the highest contribution of each RNA class and feature in 25 kb bins was depicted in a two-dimensional matrix and quantified per distance categories **(Fig. 3a-c)**. A clear trend for local interactions emerged, highlighted by a diagonal signal dominated by intronic signal from protein-coding genes (**Fig. 3a,b**). On the one hand, we observed that the number of interactions in *cis* from intronic regions increased with the distance of the genomic region bound by the transcript. On the other hand, we observed a dominant contribution of exonic regions from noncoding transcripts in the *trans* interactions (**Fig. 3c**). Remarkably, a few noncoding transcripts, such as Malat1, the small nuclear RNA Gm22973 and the small nucleolar RNAs involved in splicing, exhibited extensive *trans-*interaction patterns (**Fig. 3a**). In mOPCs we observed the same trend with dominant contribution of intronic protein-coding and exonic noncoding transcripts for *cis*- and *trans*-interactions, respectively (**Fig. 3 d-f**). In addition, lncRNAs displayed preferential binding to chromatin locally (<10kb) or in *cis* (>10kb but on same chromosome) (**Supplementary Table 1**). Global patterns of interactions indicated clear differences between the 2 cell types for both *cis-* and *trans-*interactions. Specifically, we observed large domains of *cis*-interactions distributed along each chromosome with the majority of transcripts interacting with broad regions of the chromosome from which they originate in a cell type-specific manner (**Supplementary Figure 9a,b**). For example, Pvt1 contacted in *cis* large portions of its chromosome of origin in both cell types, whereas Malat1 interacted in *trans* with a significant portion of the genome, albeit with cell type-specific patterns, and Gm22973 contacted multiple chromosomes with trans-interactions but only in mESCs (**Supplementary Figure 9a,b**).

**Figure 3.**
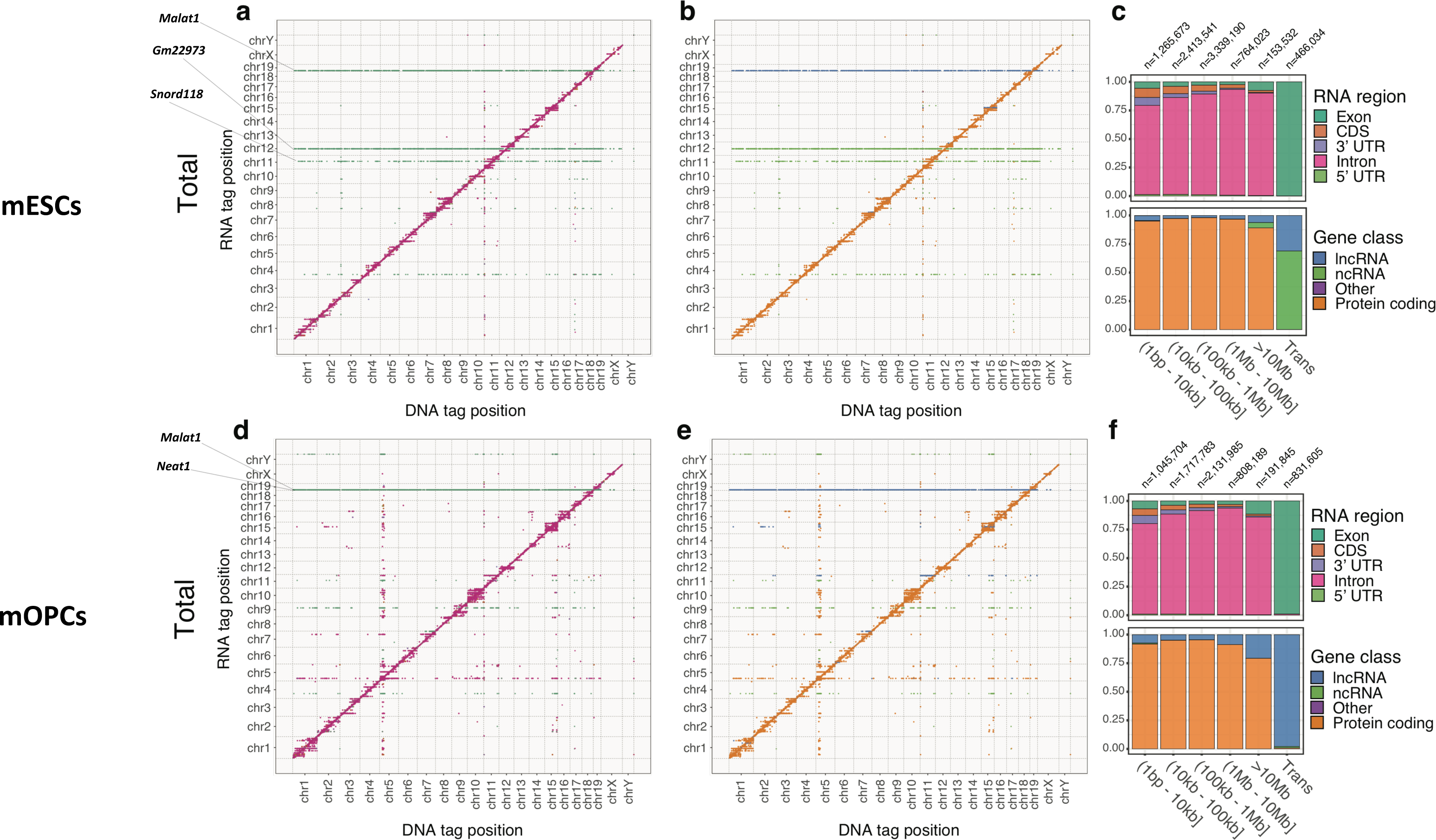
RADICL-seq identifies genome-wide RNA-chromatin interactions. a-b) RNA-DNA interactions shown as a single point per 25 kb bins and coloured by the most represented RNA class or region in that bin for mESCs. c) RNA-DNA interactions quantified for genomic distance between RNA and DNA tags for mESCs. d-e) RNA-DNA interaction matrix for mOPCs similar to a-b. f) RNA-DNA interactions quantified for genomic distance between RNA and DNA tags for mOPCs.

In the ActD-treated cells, we observed a general reduction in the number of interactions per distance category (17.7% or lower of untreated cells), however the strongest depletion of signal was in the long-range *cis*-interactions (> 100 kb; 4.9%, 1.3% and 2.2% respectively), which reflects the effect of ActD treatment on RNA pol II elongation and possibly local concentration of transcripts (**Supplementary Fig. 10a-c**). Although the diagonal was still dominated by intronic RNAs from protein-coding genes, the ActD treatment appears to have enhanced the contribution of the noncoding RNAs in the subset of long-range interactions in *cis* (> 10 Mb). Interestingly, a significant number of *trans-*interactions was preserved, and appeared in the same regions of the genome as in the total dataset (**Supplementary Fig. 10a-c**).

In both NPM datasets, the broadening of the signal from the diagonal was completely lost (**Supplementary Fig. 10 d,e,g,h**). The genome-wide binding of specific noncoding RNAs is absent, though we observed genomic regions highly bound by RNAs with cis and trans contacts. Moreover, we observed a dramatic drop in exonic RNA interactions from noncoding genes in the *trans*-interactions, which suggests that the majority of these interactions are protein-mediated (**Supplementary Fig. 10 f,i**).

To further investigate the nature of direct binding between RNA and DNA molecules in the NPM dataset we looked into the distance distribution of interacting RNA and DNA tags (**Supplementary Fig. 11a**). We observed a dominant contribution of interactions where both RNA and DNA tags were complementary to each other (compared to only 4.5% for the total condition) (**Supplementary Fig. 11a,b**). Next, we looked at the overlap of DNA tags captured by RADICL-seq in NPM and total datasets with available DRIP-seq data in mESCs that maps the location of R-loops genome-wide^24^ (**Supplementary Fig 11c**). There was a relative enrichment of RADICL-seq DNA tags across DRIP-seq peaks in the NPM dataset, possibly indicating an over-representation of R-loops compared to the total condition. As the remaining fraction of direct RNA-DNA interactions in the NPM dataset could not be explained by complementarity of RNA and DNA strands, we examined if these interactions could be mediated by the formation of triple-helical nucleic acid structures^25^. To this end, we analyzed the interactions of lncRNAs that are known to form triple helices with DNA^25, 26^, namely Malat1 and Meg3. We found that these transcripts were among the lncRNAs with the highest number of trans-contacts (**Supplementary Table 2**) and that they are likely to form triplexes in close proximity to these contacts (**Supplementary Fig. 11d-i**).

### Influence of three-dimensional chromatin architecture on RNA-DNA interactions

To better understand the relationship between the interactions captured by RADICL-seq and 3D architecture of the genome, we leveraged Hi-C data produced from similar cell types (mESCs and neural progenitor cells)^27^. At 25 kb resolution, RADICL-seq RNA-DNA *cis*-contacts were shown to moderately correlate with normalized Hi-C DNA-DNA contacts (Pearson’s correlation coefficient = 0.56 for both cell types, **Supplementary Fig. 12**). However, about 30% of the variance (based on Pearson’s correlations) in intra-chromosomal RNA-DNA contact frequency could be explained by DNA-DNA contacts and/or genomic distance. The remaining ∼70% of variance in RNA-DNA contacts is most likely a combination of noise and true signal that is not linearly dependent on spatial distance between genomic loci.

We investigated the distribution of DNA tags with respect to topologically associating domains (TADs) and found a clear signal enrichment at TAD boundaries in both cell types (**Fig 4a,b**). This finding was replicated in all other experimental conditions (**Supplementary Fig. 13a,b**). Furthermore, we observed enrichment for RNA tags at TAD boundaries primarily in NPM conditions in both cell types (**Fig. 4c,d** and **Supplementary Fig 13c,d**). Interestingly, we obtained a similar enrichment for DRIP-seq signal at the TAD boundaries, possibly suggesting a relationship between TADs formation and generation of R-loops (**Supplementary Figure 13e**). These results underline the importance of including the NPM condition to capture biological features that might be otherwise overlooked by solely using total datasets.

**Figure 4.**
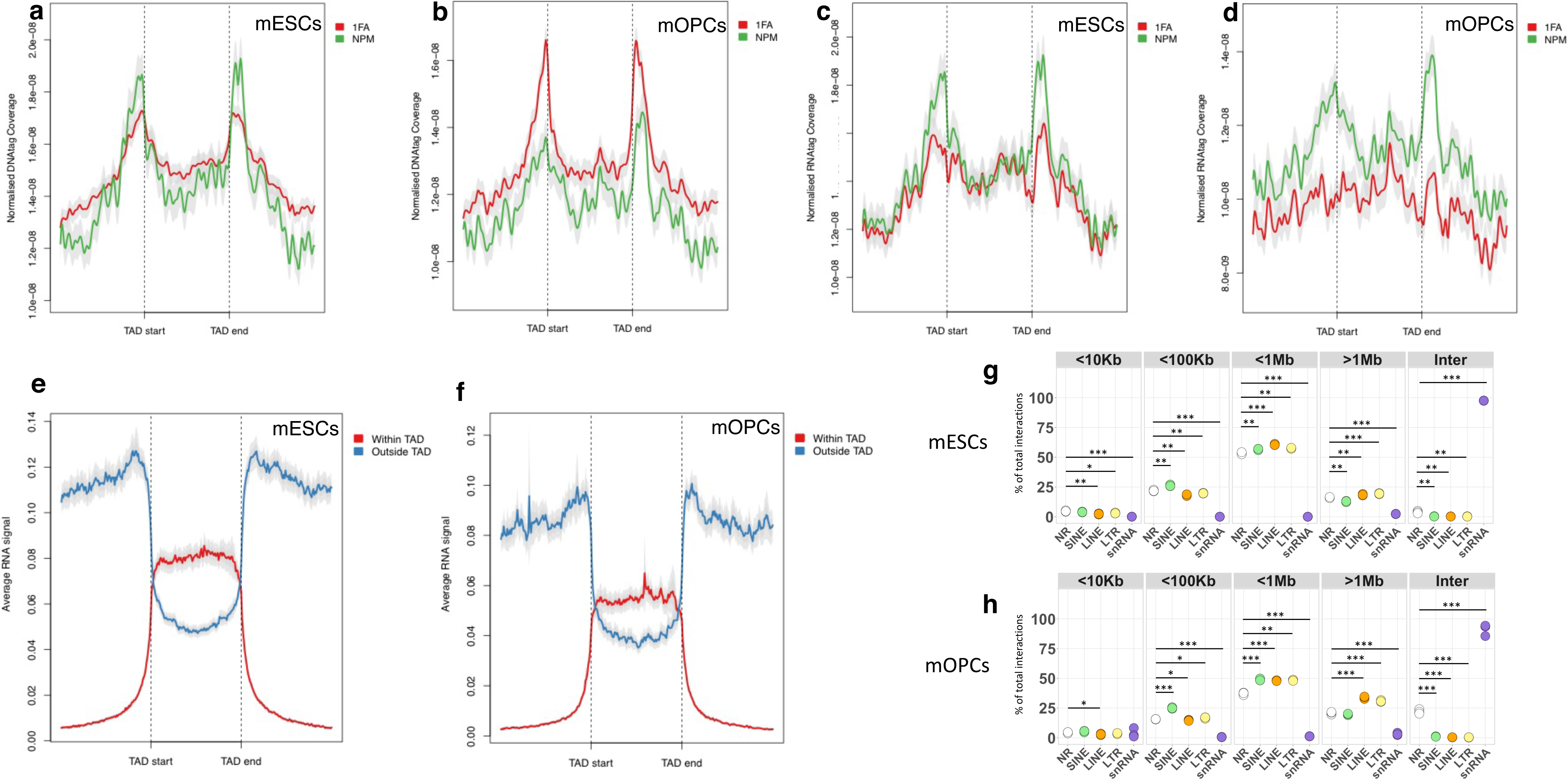
RNA interactions and genomic structural features. a-b) Metadata profiles showing the average coverage of DNA tags at the boundaries of TADs in mESCs and mOPCs for each condition. c-d) Metadata profiles showing the average coverage of RNA tags at the boundaries of TADs in mESCs and mOPCs for each condition. e-f) Average RNA binding signal at TAD boundaries in mESCs and mOPCs. The signal was split based on whether the RNA originated within or outside of the TAD. g-h) Percentage of RNA-DNA interactions in mESCs (g) and mOPCs (h) 1%FA RADICL-seq libraries divided in incremental RNA-DNA distance and with RNA tags grouped by the identity of intersected repeat elements (“NR”, RNA tags not mapped on any repeat). Statistical significance was calculated with two-tailed t-test. * = p≤0.05; ** = p≤0.001; *** = p≤0.001.

Next, we asked whether RNAs originating from loci positioned within or outside TADs showed specific DNA binding patterns, possibly dictated by genomic structural constraints. When we looked at the distribution of the signal for RNAs transcribed within TADs, we found it to dramatically drop outside the domain regions (**Fig. 4e,f**), whereas the signal from transcripts transcribed outside TADs was showing the opposite trend, thus suggesting a “barrier effect” for the RNA migration into or from TADs that prevents free diffusion (**Fig. 4e,f**). Collectively, our results highlight a putative role for TADs in shaping RNA-chromatin interactions in mESCs, mOPCs and possibly other cell types.

### Transcripts containing repeat elements are differentially engaged in specific chromatin interactions

Repeat elements (REs) have emerged in recent years as key contributors to genomic regulation and organization ^28^. In the mESCs and mOPCs RADICL-seq significant datasets we observed that ∼12% and ∼8% of the uniquely mapped genic RNA tags intersected respectively with REs as defined by RepeatMasker^29^ (**Supplementary Fig. 14**). In mESCs the most abundant classes of intragenic REs were snRNA (∼39%) and SINE (∼35%), followed by LINE and LTR (both ∼9%) (**Supplementary Fig. 15**). In mOPCs the most abundant classes were SINE (∼48%) followed by LINE (∼20%) and LTR (∼15%); intriguingly, the frequency of snRNAs involved in RNA-DNA interactions in mOPCs was dramatically reduced compared to mESCs (<1%) (**Supplementary Fig. 15**). We annotated non-self RNA-chromatin interactions for the most abundant classes of intragenic REs across increasing distances from the site of transcription. When compared them to interactions not involving REs, we found a remarkably well-defined REs-specific intra-chromosomal pattern that was reproducible in both cell types (mESCs and mOPCs) (**Fig. 4g,h**). In terms of differences among different RE families, RNA-DNA pairs where the RNA mapped to SINE were found to be enriched at distance intervals >10kb and <1Mb, whereas RNA that mapped to LINE and LTR were proportionally depleted at linear distances below 100kb but significantly enriched at longer range intervals (<1Mb, >1Mb, **Fig. 4g,h**) even in the absence of nascent transcription (**Supplementary Fig. 16**). Although these pairs displayed no *trans-*interactions, those mapping to snRNAs exhibited extensive *trans*-interactions (>95%, **Fig. 4g,h** and **Supplementary Fig. 16**). Collectively, these analyses show that transcripts containing REs are engaged in *cis*-interactions with the chromatin, which is in agreement with previous studies that reported their association with euchromatin^28^.

**Figure 5.**
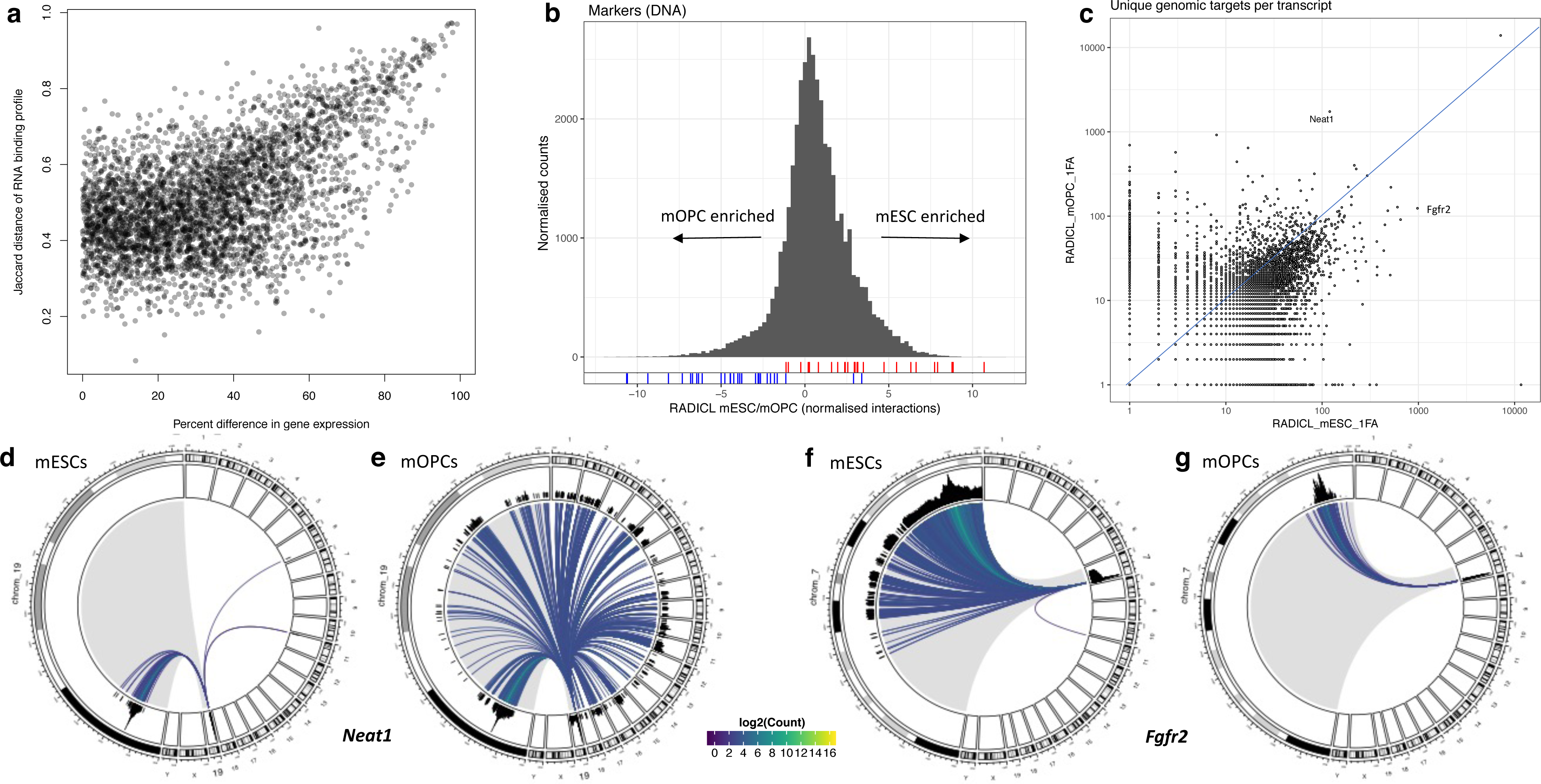
Cell type-specific RNA-chromatin interaction patterns. a) The Jaccard distance of genome-wide RNA-DNA binding profiles between cell types (mESC vs mOPC) for each gene compared to their difference in gene expression between the two cell types. b) Distribution of RADICL-seq mESC to mOPC DNA tags ratio for targeted gene promoter regions, which are defined as ± 2kb around the transcription start site (TSS). Cell type-specific marker genes positions are highlighted for both mESC (red) and mOPC (blue). c) RADICL-seq mESC versus mOPC count of unique genomic targets per interacting RNA. d-e) Circos plots depicting Neat1 genomic interactions in mESC and mOPC. f-g) Circos plots depicting Fgfr2 genomic interactions in mESC and mOPC. Each line represents the interaction between the RNA gene and the contacted DNA locus, while its color highlight the occurrence of the association. The chromosome of origin of the RNA under investigation is shown enlarged on the left portion of each circos plot.

### RADICL-seq identifies cell-type specific RNA-DNA interactions

To better understand the involvement of RNA in genome organisation and fine-tuning cell-specific gene expression, we compared gene binding patterns and capture rates between mESCs and mOPCs. We calculated the Jaccard distance between genome-wide DNA binding profiles as a function of the difference in capture rate, here considered a proxy for gene expression (**Fig. 5a**). As expected, there was a clear relationship between differential transcript abundance and variety in DNA binding profiles. However, even at comparable expression levels we observed a diversity of interaction patterns among different transcripts between the two cell types (**Supplementary Fig**. **17a-d**). When transcripts were divided into differential expression deciles, for genes with < 40% difference in gene expression between the two cell types we found no link between differential gene expression and changes in genome-wide binding profile (**Supplementary Fig. 17e**). These results suggest that although expression plays a role in determining the binding pattern of a transcript, the DNA binding profile for genes expressed at similar levels may be governed by other factors, such as three-dimensional chromatin architecture or epigenomic state.

Next, we assessed whether RADICL-seq can be used to discriminate biologically relevant differences between cell types. To this end, we collected from literature a list of marker genes that are specifically expressed in these two cell types and use RADICL-seq to analyze patterns of RNA-chromatin regulation at these genomic regions (**Supplementary Table 3**). To compare cell-specific chromatin interactions at gene level, we calculated the mESC to mOPC ratio of the normalized counts of bound RNAs for the promoter region of all genes. We used the mESC to mOPC distribution to highlight the positions of the cell-specific markers and found that these segregated towards the tails of the distribution according to the cell line in which they were expected to be more highly expressed (**Fig. 5b**). Our results suggest that the technology is able to discriminate cell type-specific features. To investigate in greater detail whether RNA-chromatin interactions could possibly play a role in gene expression, we generated CAGE data from the nuclear fraction of both cell types to annotate *de novo* promoters, and calculated the distribution of unique RNAs interacting within these regions. Although we observed comparable distributions, promoters of genes interacting with the highest number of unique RNAs clustered in different chromosomes in the two cell types (**Supplementary Fig 18a,b**). Furthermore, we looked at the distribution of unique RNAs interacting with promoter regions of genes transcriptionally active and compared to those that are inactive. In both cell types we did not observe any significant difference between the two groups, indicating that gene expression does not seem to be dependent on the number of unique interacting RNAs (**Supplementary Fig. 18c,d**).

When looking at the number of unique genomic targets for transcripts detected by RADICL-seq in both cell types we observed a rather linear correlation with the presence of some outliers (**Fig. 5c**). Among the RNAs that showed the larger deviations from the diagonal (*i.e.*, transcripts that have more dissimilar patterns of interactions) we selected Neat1 and Fgfr2 as representative example for mOPCs and mESCs, respectively. The nuclear lncRNA Neat1 is one of the main components of paraspeckles, membrane-less compartments present in the nucleus of differentiated cells^30^. In mESCs Neat1 exists as a shorter isoform which is unable to promote the formation of paraspeckles, whereas its longer isoform is expressed in differentiated cells^31, 32^. Consistent with these observations RADICL-seq RNA reads mapped only to the shorter isoform in mESCs, as opposed to mOPCs where the RADICL-seq signal covered the whole span of the longer isoform (**Supplementary Fig. 19a**). Furthermore, the genomic binding pattern of Neat1 exhibited dramatic differences between the two cell types, with the lncRNA interacting mostly in *cis* in mESCs as opposed to the extensive *trans*-interactions mediated in mOPCs (**Fig. 5d,e**). Analysis of the DNA binding regions in mOPCs confirmed a preference for the 5’ end of the target genes^33^ (**Supplementary Fig. 19b,c**). Fgfr2 is a protein-coding gene with an important role in pluripotency, as mutations affecting its expression result in early embryonic lethality due to inner cell mass defects^34^. Unlike the localized pattern exhibited in mOPCs, Fgfr2 displays extensive *cis*-interactions in mESCs, covering above 30% of the chromosome from which it is transcribed (**Fig. 5f,g** and **Supplementary Fig. 9a**). RADICL-seq results thus suggest a potentially novel structural role for Fgfr2 in mESCs.

Finally, we turned our attention to *trans* RNA-DNA interactions. To compare the extent of overlap between the two cell types, we calculated the intersection of RNA-chromatin interactions divided by linear distance (**Supplementary Fig. 20**). To our surprise, we found 3,414 unique RNA-DNA pairs shared by both cell types, accounting for an extent of overlap comparable to RNA-chromatin interactions separated by a linear distance of over 1 Mb (**Supplementary Fig. 20**). Furthermore, these *trans*-interactions captured in both cell types were mediated by 14 transcripts with a major contribution from Malat1 (**Supplementary Table 4**). These results highlight the contribution of Malat1 in the organization of general principles of RNA-chromatin interactions.

## Discussion

RADICL-seq provides four main advantages over existing methods: i) chromatin shearing is achieved by a controlled DNase I digestion, which results in a greater resolution compared to digestion with restriction enzymes. Indeed, the distribution of cut sites for restriction enzymes is often uneven and may result in the inability to detect important interaction chromatin regions. The sequence-independent digestion of chromatin by DNase I enables RADICL-seq to overcome such resolution limitations. ii) The use of paramagnetic carboxylated beads as carriers for the nuclei allows additional washes to remove small genomic fragments (upon DNase digestion) and the excess of biotinylated bridge adapter, thus reducing noise. Furthermore, beads improve visualization of the nuclear pellet when using fewer cells and can potentially decrease the number of input cells for future applications. iii) Digestion of RNA-DNA hybrids with RNase H reduces the fraction of nascent RNA-chromatin interactions captured by RADICL-seq, thereby increasing the capture rate of other types of interaction. iv) Use of EcoP15I to generate RNA and DNA reads of uniform size greatly improves unique alignment to the genome.

Furthermore, RADICL-seq uses the same amount of input cells as GRID-seq but it achieves a higher detection power for uniquely mapped RNA-DNA interactions and consequently a better performance/cost ratio.

Our results confirm previous observations regarding modality of interaction for lncRNAs, with the majority of non-coding transcripts binding genomic targets locally or in *cis*^35^. However, one surprising finding is the dominant contribution of intronic RNA sequences from protein-coding genes in *cis*-interactions. These binding events seem to be of stable nature, as we observe interactions mediated by intronic RNAs following inhibition of transcription elongation for several hours. Excised introns have been reported to exert a biological function on growth phenotype in yeast^36^. We speculate the existence of a similar mechanism in higher eukaryotes where specific intronic RNA sequences might escape degradation and interact with the chromatin in *cis*. Additionally, a subset of interactions mediated by intronic RNAs from protein-coding genes might be involved in the local transcriptional regulation mediated by protein-coding transcripts as previously reported^37^.

We hypothesized an involvement of the genome structure in transcriptional regulation mediated by RNA interactions and used RADICL-seq to assess the frequency of RNA-DNA interactions at TAD boundaries. The widespread enrichment of RNA-chromatin interactions indicated a possible role for transcription or for its products in influencing genome 3D structure. CHAR-seq technology identified enrichment of transcription-associated RNAs at TAD boundaries in *Drosophila melanogaster*, suggesting an evolutionary conservation for this phenomenon. Intriguingly, Heinz and colleagues have reported that transcription elongation remodels chromatin 3D architecture by displacing TAD boundaries^38^. Moreover, SINE sequences have been previously found to be enriched at TAD boundaries^39^. With RADICL-seq we have uncovered specific patterns of interactions for transcripts overlapping REs, thus suggesting that transcripts from REs might facilitate the formation of 3D structures in cells, especially after cell division when TADs are temporarily dismantled.

The enrichment of chromatin-associated RNAs at TAD boundaries also in the ActD dataset – where RNA pol II elongation is inhibited - potentially suggests that TADs might generate a “barrier effect”, which is consistent with the transcripts preferentially binding to DNA within or outside, but not across, TADs. This observation confirms recent evidence of a TAD-restricted genome occupancy mediated by immune gene-priming lncRNAs^40^.

The NPM datasets identified reproducible RNA-chromatin interactions that are not mediated by the presence of proteins. Such interactions can be explained either by direct RNA-DNA pairing or binding of RNA to duplex DNA via the formation of triplexes. Increased frequency of RNA-chromatin interactions and enrichment for R-loops at TAD boundaries in NPM datasets indicate the possible role of active transcription in partitioning the genome as disruption of the cohesin/CTCF complex in mammals does not lead to disappearance of TAD boundaries^41^. Although the RADICL-seq protocol includes an RNase H step to remove RNAs paired with DNA in Watson-Crick fashion while the sample is still crosslinked, proteins located nearby the complementary RNA-DNA binding could hinder enzyme accessibility, thus preventing the complete digestion of these hybrids.

Nevertheless, we were still able to observe a consistent number of interactions that can be explained by triple helix formation. This is indeed an enticing perspective for future extensions of the protocol that could include an RNase H step after reversal of crosslinks to enrich for triplexes structures. Although a protein-free method that separately identifies RNA and DNA involved in triple helix formation has been published^42^, to our knowledge RADICL-seq performed in the NPM condition is the first approach that can simultaneously link interacting RNA and DNA to triplex structures.

Our results further show that RADICL-seq can pinpoint cell type-specific RNA-chromatin interactions. We uncovered a mOPC-specific genome occupancy pattern for Neat1 RNA where the long isoform interacts in *trans* with the chromatin. Recently, Katsel and colleagues have reported a down-regulation of Neat1 expression in schizophrenia patients that associated with reduction in the number of cells of the oligodendrocyte lineage^43^.

In summary, we have developed a technology to map genome-wide RNA-chromatin interactions that significantly improves upon other existing technologies by reducing nascent transcription bias, and increasing genomic coverage and unique mapping rate efficiency. Application of RADICL-seq in mESCs and mOPC-derived cells allowed to unveil novel principles of RNA-chromatin *cis-* and *trans*-interactions, and to identify the cell type specificity of such associations. We anticipate that the RADICL-seq technology will pave the way for a deeper understanding of the fine regulatory network governing gene expression and ultimately cell identity.

## Online Methods

### Cell Culture

mES R08 cells were grown under feeder-free conditions in mouse ESC medium containing DMEM (Wako), 1,000 U/ml leukemia inhibitory factor (LIF; Millipore), 15% FBS (Gibco), 2.4 mM l-glutamine (Invitrogen), 0.1 mM non-essential amino acids (NEAA; Invitrogen), 0.1 mM 2-mercaptoethanol (Gibco), 50 U/ml penicillin and 50 µg/ml streptomycin (Gibco). Culture media were changed daily, and cells were passaged every 2–3 days. For the ActD-treated RADICL-seq libraries, mES R08 were treated with Actinomycin D (Sigma) at a final concentration of 5 µg/ml for 4 h before crosslinking, as described below.

Oli-neu cells were grown on poly-L-lysine coated dishes and expanded in proliferation media consisting of DMEM (Lifetech 41965062), N2 supplement, Pen/Strep Glu (Lifetech 10378016), T3 (Sigma-Aldrich T6397) 340ng/ml, L-thyroxine (Sigma-Aldrich 89430) 400 ng/ml, bFGF 10ng/ml and PDGF-BB 1ng/ml. Culture media was changed on alternate days and the cells were passaged every 5-6 days.

### Crosslinking of cells

Confluent cells were rinsed with pre-warmed phosphate-buffered saline (PBS) and trypsinized. Detached cells were pelleted, resuspended in PBS, counted and pelleted again. Cell pellets were then crosslinked by resuspension with freshly prepared 1% or 2% formaldehyde (Thermo Fisher) solution using 1 ml for every one million cells. Cells were incubated at room temperature for 10 minutes with rotation followed by quenching with 125 mM glycine (Sigma). Cells were pelleted at 4 °C, washed with ice-cold PBS, pelleted again and snap frozen in liquid nitrogen.

With the term biological replicates, we refer to batches of cells having different passage number.

### Generation of RADICL-seq libraries

#### Adenylation of the adapter

The adapter is a partially double-stranded DNA molecule containing chemical modifications (IDT). The upper strand sequence is 5’-/5Phos/CTGCTGCTCCTTCCCTTTCCCCTTTTGGTCCGACGGTCCAAGTCAGCAGT-3’. The lower strand sequence is 5’-/5Phos/CTGCTGACT/ibiodT/GGACCGTCGGACC-3’. The upper strand was pre-adenylated by using DNA 5′ Adenylation Kit (NEB). The pre-adenylated upper strand was mixed with equimolar quantity of the lower strand and subsequently incubated at 95°C for 2 min followed by 71 cycles of 20 s, with a reduction of 1 °C every cycle. Annealed pre-adenylated adapter was then purified using Nucleotide Removal kit (Qiagen) following manufacturer recommendations.

#### Chromatin digestion

Chromatin preparation was performed as described previously^44^ with some modifications. Briefly, cell pellets containing approximately two million crosslinked cells were resuspended in cold lysis buffer (10 mM Tris–HCl pH 8.0, 10 mM NaCl, 0.2 % NP-40) and incubated on ice for 10 min. Nuclei were pelleted at 2,500 g for 60 s, resuspended in 100 µl of 0.5× DNase I digestion buffer (0.5× DNase I digestion buffer (Thermo Fisher), 0.5 mM MnCl₂) containing 0.2 % SDS, and incubated at 37 °C for 30 min. An equal volume of 0.5× DNase I digestion buffer containing 2 % Triton X-100 was added and incubation at 37 °C was continued for 10 min. Then, 1.5 U DNase I (Thermo Fisher) was added and digestion was carried out at room temperature for 4 or 6 min (1% or 2% FA, respectively). DNase I digestion was stopped by adding 40 µl of 6× Stop Solution (125 mM EDTA, 2.5 % SDS), followed by centrifugation at 2,500 g for 60 s. Nuclei were resuspended in 150 µl nuclease-free H_2_O and purified with two volumes (300 µl) of AMPure XP magnetic beads (Beckman Coulter). After 5 min incubation at room temperature, beads were separated using a magnetic rack, washed twice with 80 % ethanol and air dried for 2 min.

#### Chromatin end-repair, dA-tailing and RNase H treatment

The purified bead-nuclei pellet was resuspended in 200 µl 1× T4 DNA Ligase Buffer (New England Biolabs) containing 0.25 mM dNTPs, 0.075 U/µl T4 DNA polymerase (Thermo Fisher) and 0.15 U/µl Klenow Fragment (Thermo Fisher), and incubated at room temperature for 1 h. The end-repair reaction was stopped by adding 5 µl of 10 % SDS. The bead-nuclei mixture was pelleted at 2500 g for 60 s, resuspended in 200 µl 1× NEBuffer 2 (New England Biolabs) containing 0.5mM dATP, 1 % Triton X-100 and 0.375 U/µl Klenow (exo-) (Thermo Fisher), and incubated at 37 °C for 1 h. After that, 0.122 U/µl RNase H (New England Biolabs) was added and the reaction was incubated at 37 °C for further 40 min. The dA-tailing and RNase H reactions were stopped by adding 5 µl of 10 % SDS.

#### Bridge adapter RNA ligation

The bead-nuclei mixture was pelleted at 2,500 g for 60 s and resuspended in 200 µl H_2_O. To remove soluble RNA, 165 µl of 20 % PEG in 2.5 M NaCl was added to the mixture, followed by 5 min incubation at room temperature. Beads were collected with magnetic rack, washed once with 80 % ethanol and resuspended in 200 µl H_2_O. This purification step was repeated once. After second ethanol wash, the air-dried bead-nuclei mixture was resuspended in 23 µl H_2_O, 3 µl 10× T4 RNA Ligase Buffer, 1 µl pre-adenylated and biotinylated bridge-adaptor (20µM), 1 µl RNaseOut (Thermo Fisher), 13.3 U/µl T4 RNA Ligase 2, truncated KQ (New England Biolabs). The mixture was incubated at 20 °C overnight to ligate the pre-adenylated bridge adapter to the 3’-OH of the RNA molecules. The reaction was stopped by adding 5 µl 10 % SDS and the bead-nuclei mixture was then pelleted at 2,500 g for 60 s and resuspended in 200 µl H_2_O. To remove excess unligated adapter, 165 µl of 20 % PEG in 2.5 M NaCl was added to the mixture and the reaction was incubated at room temperature for 5 min. Beads were then collected with a magnetic rack, washed once with 80 % ethanol and resuspended in 200 µl H_2_O. This purification was repeated once.

#### Proximity ligation

The *in situ* proximity ligation was carried out by resuspending the air-dried bead-nuclei mixture in 500 µl 1× T4 DNA Ligase Buffer with ATP containing 4 U/µl T4 DNA Ligase (New England Biolabs) and incubating it at room temperature for 4 h. After the incubation, bead-nuclei complexes were pelleted at 2,500g for 60 s and resuspended in 200 µl H_2_O. To remove remaining unligated and DNA-only ligated adapter, 165 µl of 20 % PEG in 2.5 M NaCl was added to the mixture, and purified as previously described. Beads were then resuspended in 200 µl H_2_O.

#### Reversal of crosslinking and purification of RNA-DNA complexes

Reversal of crosslinks was performed by adding 50 µl of proteinase K solution (10 mM Tris-HCl pH 7.5, 1% SDS, 15 mM EDTA) and 1.6 U/µl proteinase K (Ambion) to the resuspended beads. The mixture was incubated overnight at 65 °C and the RNA-DNA complexes were then precipitated with 3 µl GlycoBlue (Ambion), 28 µl 3 M sodium acetate pH 5.2, and 303 µl isopropanol and for 1h on ice followed by 15,000 rpm centrifugation at 4 °C for 30 min. The resulting bead-nucleic acids pellet was resuspended in 100 µl H_2_O and further purified using 100 µl AMPure XP beads. After 5 min incubation at room temperature, beads were separated using magnetic rack, washed twice with 80 % ethanol and air dried for 2 min. DNA was eluted using 130 µl H_2_O and quantified with Qubit High Sensitivity kit (Thermo Fisher).

#### Reverse transcription and second strand synthesis of the RNA-DNA complexes

Since reverse transcriptase can use DNA sequences as primers for the polymerization, the double stranded region of the bridge adapter acts as the primer for the reaction. The RNA ligated to the bridge adapter was reverse transcribed after the sample was concentrated to a final volume of 12 µl. First, 1 µl 10 mM dNTPs was added to the sample and the mixture was incubated at 65 °C for 5 min. Subsequently, 4 µl of 5× first strand buffer, 1 µl of DTT 0.1 M, 1 µl of RNaseOut and 1 µl of SuperScript IV (Thermo Fisher) were added and the reaction was incubated at 56 °C for 10 min and 80 °C for 10 min.

Next, the generated cDNA-RNA hybrid was converted to double stranded DNA through a second strand synthesis reaction by addition of 30 µl 5× Second Strand buffer (Thermo Fisher), 3 µl 10 mM dNTPs (Thermo Fisher), 3 µl RNase H (2U/µl, Thermo Fisher), 4 µl *E. coli* DNA polymerase I (New England Biolabs), 1 µl *E. coli* ligase (New England Biolabs) and H_2_O to the RT sample, in a final volume of 150 µl. The mixture was incubated at 16 °C for 2 h and the reaction was stopped by adding 10 µl of 0.5 M EDTA. The sample was purified using Nucleotide Removal Kit (Qiagen) by adding 1.6 ml of buffer PNI to the sample and following the manufacturer recommendations, with a final elution in 50 µl H_2_O. Sample volume was reduced to 8 µl using Speedvac concentrator (Tomy).

#### Hairpin ligation and EcoP15I digestion of the cDNA-DNA complexes

The sample was then subjected to hairpin linker (5’-/5Phos/GGCCCTCCAAAAGGAGGGCA-3’, IDT) ligation to selectively ligate the bridge adapter that was covalently bound to RNA only and therefore prevent subsequent ligation of sequencing adapters. A total of 100 pmol of hairpin linker was mixed with 10 µl of 2× Quick ligase buffer (New England Biolabs), 8 µl of sample and 1 µl of Quick ligase (New England Biolabs). The reaction was carried out for 15 min at room temperature and was then purified with DNA Clean & Concentrator-5 kit (Zymo) according to manufacturer instructions. Elution was done in 50 µl of H_2_O and final volume was reduced to 30 µl.

The sample concentration was measured by Qubit dsDNA High Sensitivity kit (Invitrogen). EcoP15I digestion of the double-stranded cDNA-DNA complexes was performed using 10U of enzyme for 1.5 µg of DNA in presence of 5 µl NEBuffer 3.1 (New England Biolabs), 5 µl 10× ATP, 0.5 µl 10 mM sinefungin (Calbiochem), and H_2_O in a final reaction volume of 50 µl. The sample was incubated at 37°C overnight.

#### Ends preparation and sequencing linkers ligation

EcoP15I-digested sample was purified using Nucleotide Removal kit (Qiagen) by adding 1.3 ml of PNI buffer and following manufacturer recommendations. Sample was eluted in 50 µl and volume was further reduced to 20 µl. To prepare the sample for the sequencing linkers ligation, 6.5 µl of 10× reaction buffer, 3 µl of End Prep Enzyme Mix from NEB Next Ultra End Repair/dA-Tailing Module (NEB) and H_2_O up to 65 µl were added to the concentrated sample and the reaction was incubated at 20 °C for 30 min and 65 °C for 30 min.

Next, Y-shaped sequencing linkers were prepared. The upper strand (5’- /5Phos/GATCGGAAGAGCGTCGTGTAGGGAAAGAGTGT-3’) and lower strand (5’- CTCGGCATTCCTGCTGAACCGCTCTTCCGATCT-3’) were annealed in 1× NEBuffer 2 (New England Biolabs) at 95 °C for 2 min followed by 71 cycles of 20 s, with a reduction of 1 °C every cycle.

Sequencing linkers ligation was performed with NEB Next Ultra Ligation Module and 20 pmol of annealed Y-shaped sequencing linkers, at 20 °C for 15 min. After ligation, the sample volume was reduced to 40 µl.

#### Pull-down of the RNA-DNA ligated complexes and PCR titration

The RNA-DNA ligated complexes were pulled-down with MyOne C1 Streptavidin magnetic beads (Invitrogen). A total of 20 µl of beads were washed twice with 1× WB buffer (5 mM Tris-HCl pH 7.5, 0.5 mM EDTA, 1 M NaCl, 0.02% Tween-20), once with 2× WB buffer and finally resuspended in 40 µl 2× WB buffer. Equal volume of sample was added to the beads and the mixture was incubated at room temperature for 20 min with rotation. Isolated RNA-DNA ligated complexes were extensively washed three time with 1× WB buffer, once with EB buffer (Qiagen) and finally resuspended in 30 µl EB buffer.

PCR cycle check was performed by using Phusion High Fidelity PCR kit (Thermo Fisher), Universal FW primer (5’- AATGATACGGCGACCACCGAGATCTACACTCTTTCCCTACACGACGCTCTTCCGATCT-3’, Invitrogen), Index RV primer (5’– CAAGCAGAAGACGGCATACGAGATBBBBBBCTCGGCATTCCTGCTGAACCGCTCTTCCGA TCT-3’, Invitrogen. BBBBBB consists of a 6 nt barcode for multiplexing libraries) and 4 µl of isolated libraries. PCR was carried out at 98 °C for 30 s followed by 14 cycles of 98 °C for 10 s, 65 °C for 15 s and 72 °C for 15 s. After respectively 8, 11 and 14 cycles, 10 µl aliquots were collected and run on pre-cast 6% polyacrylamide gel (Invitrogen) at 145V for 60 min. The lowest PCR cycle where the 220 bp band representing the RNA-DNA ligated complexes could be visualized was chosen for the final library amplification.

#### Library amplification and sequencing

A total of four PCR reactions, each using a different barcoded primer, were prepared for each library. After amplification, the four reactions were pooled and run on pre-cast 6% polyacrylamide gel (Invitrogen) at 145V for 60 min. The 220 bp band was excised and purified.

Library size was assessed using High Sensitivity DNA Bioanalyzer kit (Agilent) and quantified by qPCR using the Library Quantification Kit for Illumina sequencing platforms (KAPA Biosystems) and StepOne Real Time PCR System (Applied Biosystems). Sequencing was performed with single-end 150-bp kit on Illumina HiSeq-2500 platform using the sequencing primer 5’- ACACTCTTTCCCTACACGACGCTCTTCCGATCT-3’.

#### Generation of libraries capturing non-protein-mediated interactions

Libraries were produced following the protocol for the generation of RADICL-seq libraries with modifications below. After the purification steps that followed dA-tailing and RNase H reactions, reversal of crosslinks was performed as described above. Samples were concentrated to a final volume of 13.9 µl, followed by the addition of 3 µl 10× T4 RNA Ligase Buffer, 1 µl pre-adenylated and biotinylated bridge-adaptor (20µM), 1 µl RNaseOut (Thermo Fisher), 13.3 U/µl T4 RNA Ligase 2, truncated KQ (New England Biolabs) and 9 µl 50% PEG. The mixture was incubated at 20 °C overnight to ligate the pre-adenylated bridge adapter to the 3’-OH of the RNA molecules. Next, bridge adapter-ligated molecules were purified with DNA Clean & Concentrator-5 kit (Zymo) according to manufacturer instructions and eluted in 50 µl of H_2_O. DNA ligation was carried out by adding 450 µl of 1× T4 DNA Ligase Buffer with ATP containing 4 U/µl T4 DNA Ligase (New England Biolabs) and incubating it at room temperature for 4 h. The RNA-DNA complexes were purified with DNA Clean & Concentrator-5 kit (Zymo) as described above, and then subjected to reverse transcription and subsequent steps of RADICL-seq library preparation and sequencing.

#### Generation of libraries treated with different enzymes targeting the RNA

Libraries were produced following the protocol for the generation of RADICL-seq libraries with the following exceptions. i) For the library with no enzymatic treatment, the RNase H digestion after chromatin dA-tailing was omitted. ii) For the library generated with nuclease S1 treatment, prior to chromatin end-repair step the purified bead-nuclei pellet was resuspended in 200 µl containing 40mM sodium acetate (pH 4.5 at 25 °C), 300mM NaCl, 2mM ZnSO4 and 10U nuclease S1 (Thermo Fisher), and incubated at room temperature for 30 min. Reaction was stopped by adding 5 µl of 10 % SDS and sample followed the RADICL-seq protocol. iii) For the library generated with RNase V1 treatment, after chromatin dA-tailing step sample was digested with 0.01 units of RNase V1 (Ambion) at room temperature for 30 min after chromatin dA-tailing step. Reaction was stopped by adding 5 µl of 10 % SDS and sample followed the RADICL-seq protocol.

#### CAGE library preparation

RNA from nuclear fractions of mESCs and mOPCs were extracted as previously publishedl^11^. CAGE libraries were prepared according to Takahashi *et al.*^45^ Briefly, 3ug of nuclear RNA was used for reverse transcription with random primer. 5’ end of cDNA-RNA hybrids was biotinylated and captured using magnetic streptavidin-coated beads. After capture, cDNAs from cap-trapped RNA were released, ligated to 5’ barcoded linkers and digested with EcoP15I. The cDNA tags were then ligated with a 3’linker and amplified with 9 cycles. Libraries were sequenced on Illumina HiSeq 2500, 50bp single-end reads.

### Data analysis

#### RADICL-seq mapping and processing

RNA and DNA tags at both ends of the adapter were extracted from raw sequencing reads using TagDust2^46^ (ver. 2.31). Multiplexed reads were split by six nucleotide barcodes embedded in the 3’ linker sequence. Artificial sequences were removed using TagDust^47^ (ver. 1.1.3). Ribosomal RNAs were identified and removed from the extracted RNA tags using RNAdust^48^ (ver. 1.0.6) with a ribosomal DNA repeating unit (GenBank: BK000964.2). PCR duplicates were removed from paired RNA-DNA tags using FastUniq^49^. We aligned RNA and DNA tags separately to the mouse genome (mm10 assembly) using BWA^50^ (ver. 0.7.15-r1140) with *aln* and *samse* functions. Mapped RNA and DNA tags were paired with unique sequencing read IDs using samtools^51^ (ver. 1.3.1) and bedtools^52^ (ver. 2.17.0). These processes were run on the MOIRAI pipeline platform^53^.

#### Reference annotation and genome binning

The comprehensive gene annotation for mouse^54^ (GENCODE release M14) was downloaded and genes were divided into 4 major groups of biotypes: protein_coding, long_ncRNA (defined according to GENCODE “Long non-coding RNA gene annotation”), ncRNA (defined according to GENCODE “Non-coding RNA predicted using sequences from Rfam and miRBase”) and other (all remaining genes). The mouse genome binning at 25 kb resolution was performed using bedtools^52^ (v2.26.0) with parameter *-w 25000* (and *-w 1000000* for generating 1 Mb bins for the genomic heatmap).

#### Annotated features overlaps

RNA and DNA tags were associated unambiguously to the corresponding annotated features. To achieve that, we resized each fragment to a single nucleotide position at its centre to reduce the possibility of fragments overlapping with multiple genes/bins. For RNA tags the strand information was taken into account, while for DNA tags this was discarded. Both RNA and DNA tags were required to map uniquely (BWA MAPQ = 37) to the genome.

#### Reproducibility

The summary of interactions by condition, replicate and RNA and DNA pairs, the calculation of pairwise Pearson correlation coefficients on the counts of complete sets of observations, and the visualization of the results were performed using the *CAGEr* package^55^.

#### Correlation with RNA-seq

RADICL-seq interactions were summarised per RNA gene and normalized to reads per kilobase (RPK) to account for overrepresentation of longer transcripts. Nuclear and cytosolic RNA-seq (and CAGE-seq) reads were mapped to the mm10 genome using STAR^56^ (v2.5.0a) with default parameters and low-quality reads (MAPQ <10) were removed. The files were similarly intersected with GENCODE vM14 to generate counts and then normalized to transcripts per million (TPM). A threshold of 1 was applied to both RPK and TPM to select interacting and expressed transcripts, respectively. The square of the Pearson correlation coefficient was used to determine the dependency between variables.

#### Robust interactions calculation

The P-value calculation was performed in R using the *binom.test* function with parameters *N, P* and *alternative=’greater’*. For any given interaction *N* is the total number of interactions of the RNA involved and *P* is 1 over the total number of unique genomic bins the RNA has been observed to interact with. The P-value correction was performed using the *p.adjust* function with *method=’BH’*.

#### Genome-wide RNA-DNA interaction plots

The significant set of interactions per cell type and condition were first removed of any RNA-DNA contacts intersecting blacklisted regions reported for mm10. For the genome-wide interaction plots, the genome was divided into 25 kb bins and each interaction was assigned to these bins for both the RNA and DNA end of the tag. For each unique combination of the two bins (RNA and DNA coordinates), all interactions were collapsed into one value to avoid overplotting. The plot was produced in R with ggplot2 where each value is coloured by most represented RNA class or RNA region if more than one individual interaction was assigned to these coordinates. All interactions were then divided into 6 classes of genomic distance between the RNA and DNA end of the tag and the proportion of interactions derived from specific RNA regions and gene classes were visualised in barplots.

#### RNA-DNA interactions with respect to TAD boundaries

To investigate any relationship between RNA-DNA binding and TAD boundaries, published mESC and mouse neural progenitor cell TADs were retrieved^27^. For all cell types and conditions, the significant set of interactions were filtered to remove and contacts intersecting the mm10 blacklisted regions, and the coverage of RNA and DNA tags was visualised at the boundaries of TADs in the form of a metaplot.

To further investigate the effects of TAD boundaries on the spread of RNA-DNA interactions, interactions were split into those that originated within a TAD and those that did not. This was performed on a TAD by TAD basis, as interactions that originate outside of one TAD may originate within a neighbouring TAD. The coverage of DNA binding locations for each set of interactions was visualised in windows around TADs as metaplots. All plots were produced using the *genomation* package for R (<u>10.1093/bioinformatics/btu775</u>).

#### Comparison of RADICL-seq data with DRIP-seq

To compare DRIP-seq and RADICL-seq DNA binding patterns, mESC DRIP-seq data was retrieved from GEO:GSM1720620^24^. As with RADICL-seq RNA and DNA tags, the average coverage of DRIP-seq reads was visualised in bins around TAD boundaries as in the form of a metaplot. To investigate whether there was an enrichment for RADICL-seq interactions binding within R-loops, the average coverage of RADICL-seq DNA tags within DRIP-seq peaks was visualised in the form of a metaplot for the 1FA and the NPM conditions.

#### Correlation of differences in gene expression and genome-wide DNA binding profiles between mESCs and mOPCs

To explore the relationship between gene expression and RNA-DNA binding, the binding pattern and expression level of all genes were compared between the mESC and mOPC 1FA datasets. To compare gene expression between the cell types, the difference in normalised counts between cell types was calculated as a percentage of the maximum expression in either condition. To compare the DNA binding profiles between the cell types, the genome was divided into 5kb bins, and each bin was assigned either a 1 or 0 depending on whether there was a DNA tag present in that bin. This was performed for all genes. The binary vectors produced in this way were then compared by calculating the Jaccard distance between the vectors derived from each cell type. Subsequently, the resulting data from each analysis was visualized as a scatterplot.

#### Comparison of RADICL-seq with RAP-DNA for Malat1

Malat1 RAP-DNA data was obtained from GEO:GSE55914. The raw reads were processed with TrimGalore!^57^ (v0.4.5) with parameters *--paired --trim1* and aligned using Bowtie2^58^ (v2.3.3.1) to the mouse genome (mm10) with parameters *--no-mixed --no-discordant*. Samtools^51^ (v1.5) was employed to select uniquely mapping fragments and Picard *MarkDuplicates* (v2.9.0.21, http://broadinstitute.github.io/picard/) to remove PCR duplicates. *featureCounts*^59^ was employed to summarize by gene reads from RAP-DNA and Malat1-DNA tags from GRID-seq and RADICL-seq. Gene counts were normalized to reads per kilobase (RPK) and sorted according to the normalized values. The top 10’000 genes from each dataset were used to generate the Venn diagram (*VennDiagram* R package).

#### Comparison of RADICL-seq with ChIRP-seq for Rn7sk

The list of Rn7sk genomic target locations was retrieved from the publication^18^. Peak coordinates were lifted from mm9 to mm10 using *LiftOver* and genes overlapping peaks were considered as targeted by Rn7sk. *featureCounts*^59^ was employed to summarize by gene Rn7sk-DNA tags from GRID-seq and RADICL-seq. The Venn diagram was generated using the *VennDiagram* R package.

#### Analysis of repetitive elements

The significant RNA-DNA pairs in this dataset were generated using the same approach described elsewhere for the main RADICL-seq significant dataset. RNA and DNA tags with mapQ≥37 were processed using bedtools^52^ (v2.26.0) and RepeatMasker^29^ (v4.0.6) to generate the dataset of RADICL-seq RNAs intersecting (with options -s -wb) and not intersecting (with option -v) with Repeat Elements annotated in mm10. Self-interactions were removed from the datasets by comparing the GENCODE Gene ID associated to both the RNA and the DNA tags of the same pair. The relative percentage of interactions across different intervals of RNA-DNA distance was calculated by dividing the number of significant interactions in a given RNA-DNA distance interval by the total number of RNA tags intersecting a given repeat family for each experimental condition. P-values were calculated by unpaired T-test.

#### Transcript-specific genomic interactions

The significant interactions for the transcripts under analysis were selected, summarized by DNA bins, and reported as counts associated to genomic coordinates. The *circlize* package^60^ was used to visualize the results and the counts of the RNA-DNA pairs were used to color the connecting lines.

#### High-throughput Chromosome Conformation Capture (Hi-C) paired-end reads processing

Publicly available mESCs and neural progenitor cells Hi-C sequencing paired-end reads were obtained from Bonev *et al.*^27^ and processed using the Hi-C User Pipeline (HiCUP)^61^ (v0.5.3). HiCUP mapped Hi-C paired-end reads to the mouse (mm10) reference genome and filtered reads for expected artifacts resulting from the sonication and ligation steps (e.g., circularized reads, reads with dangling ends) of the Hi-C protocol. Data from different biological replicates were then pooled together. HOMER^62^ (v4.7.2) software was used to filter HiCUP processed Hi-C paired-end reads for a MAPQ30 (following recommendations described in Yaffe and Tanay^19^) and other HOMER recommended settings (i.e., PCR duplicates, expected HindIII restriction sites, self-ligation events, etc.). HOMER filtered Hi-C mapped paired-end reads were then binned at a fixed resolution of 25 kb and normalized for restriction fragment bias using HOMER’s *simpleNorm* algorithm.

#### Density plots

Data for density plots was extracted from RADICL-seq processed data, DNA and RNA. Random regions were extracted using bedtools *random*. Peak information of histone modification and DNase-Seq and ATAC-seq data was downloaded from ENCODE. Data of density plots were calculated using *makeTagDirectory* followed by *annotatedPeaks* in HOMER^62^ (v4.9) with parameters -size 10000 -hist 1 -histNorm 100. Finally, data was divided by mean of values of +-5000bp to align baselines. Plots were generated by ggplot2 with *geom smooth* option.

#### Triple helix search

To computationally determine the location of triplex helices in Meg3 and Malat1 trans-contacts we used spliced transcripts ENSMUST00000146701.7 and ENSMUST00000172812.2, respectively. We analysed an enrichment in triple helices in NPM trans-contacts expanded by 1000bp over genomic average. To do so, TDF region test^63^ was used (triple helix parameters: 12 bp, 1 mismatch) with 1000 random genomic background sets.

#### Distribution of unique interacting RNAs at gene promoters

DNA tags from the RADICL-seq significant RNA-DNA interaction dataset were mapped to de novo CAGE-derived gene promoters using intersectBed, allowing for a 2 kb window around the actual promoter coordinates. The number of RNA tags originating from unique genes was then calculated for each promoter. Promoters were ranked from highest to lowest number of unique interactions, and additional information was collated (gene name, nuclear CAGE expression, locus, biotype) for use in further analyses.

#### CAGE data mapping and processing

Multiplexed sequencing reads were split by barcode sequences. Reads with ambiguous bases “N” were removed, and linker sequences were trimmed from the 3′ end to create reads between 21-27 nt long. The reads were mapped to the mouse genome (mm10/GENCODE release M14) using bowtie version 1.2.2^64^ with up to 2 mismatches and only keeping unique alignments with MAPQ of 20. The CAGEr (1.20.0)^55^ R package was used for the extracting TSS, normalizing counts, and defining transcriptional clusters (TCs). Briefly, the most 5′ position of a CAGE tag represents a TSS, the 5′ coordinates were extracted from the tags and generated a genome-wide TSS map at single-nucleotide resolution. The CTSS raw counts were normalized to expression per million tags using a power-law distribution. We then clustered the tags to define distinct CAGE peaks that occurred within 20 bp and expression was summed per TC.

#### Sequencing data

RADICL-seq and CAGE sequencing data were deposited at Gene Expression Omnibus. As the process is not yet completed we have created a temporal link for the reviewers as following: https://genomec.gsc.riken.jp/gerg/owncloud/index.php/s/6l5mu0bX0hORUxj (password: temp).

## Acknowledgements

We thank Mikael Rydén for providing logistical help. This work was founded by a Research Grant from MEXT to the RIKEN Center for Life Science Technologies (http://www.mext.go.jp/en/). This work was also supported by the Francis Crick Institute which receives its core funding from Cancer Research UK (FC010110), the UK Medical Research Council (FC010110), and the Wellcome Trust (FC010110). NML is a Winton Group Leader in recognition of the Winton Charitable Foundation’s support towards the establishment of the Francis Crick Institute. NML is additionally funded by a Wellcome Trust Joint Investigator Award (103760/Z/14/Z) and the MRC eMedLab Medical Bioinformatics Infrastructure Award (MR/L016311/1). Work at the GCB lab was supported by European Union (Horizon 2020 European Research Council Consolidator Grant EPIScOPE) and Ming Wai Lau Centre for Reparative Medicine.

## Authors contribution

V.O. and P.C. conceived the original idea and managed the project; A.B and P.C. developed the RADICL-seq method and designed the experiments; A.B. and A.M.S. performed the experiments; M.G., J.G. and R.C. contributed to experimental optimization; J.G. and G.P. provided valuable feedback on the RADICL-seq method; K.K. and S. contributed to sample preparation; K.H. developed bioinformatic pipeline; A.B. and F.A. planned the computational analyses and provided interpretation of the results; F.A. performed most of the computational analyses; L.R., A.J.N., C.J.F.C., Y.A.M., S.N., M.D.H., M.V., E.A. and S.A. performed the computational analyses; J.L., M.B., M.D.H., E.A., B.L. and N.M.L. provided valuable feedback on computational analyses; T.K. provided information technology on data handling; G.C.B., C.P., V.O. and P.C. provided critical feedback and helped to shape research, analysis and manuscript; A.B., F.A. and G.P. wrote the manuscript with valuable feedback from all authors.

---

**Suppl. Fig. 1. Characterization of RADICL-seq technology.** a) Summary statistics of the sequencing outcome of the samples after testing different enzymatic treatments. b) Distribution of the linear genomic distance between RNA and DNA tags derived from the same read after treatment with different enzymes. c) Sequence and features of the bridge adapter. The adapter contains a 5’ adenylated end (App), an internal T residue with biotin modification (bold) and two restriction sites for EcoP15I. The 3’-overhanging T allows DNA ligation with dA-tailed genomic fragments. d). Generation of RADICL-seq libraries and controls. Gel migration pattern of PCR-amplified RADICL-seq library compared with controls generated by omitting DNA ligase or RNA ligase. The expected band was detected only when both RNA and DNA ligases were used for the construction of the library.

**Suppl. Fig. 2. Features of RADICL-seq libraries.** a) Summary statistics of the 1% and 2% formaldehyde treatments and replicates sequencing results. b) Reproducibility of the RNA-DNA interaction frequencies across formaldehyde treatments and replicates, assessed by counting the occurrences of transcribed genes and 25 kb genomic bins pairs. c) Distribution of RNA and DNA tag biotypes according to their mapped loci and gene annotation.

**Suppl. Fig. 3. Features of RADICL-seq libraries from ActD and NPM datasets.** a) Summary statistics of the ActD treated and NPM samples and replicates sequencing results. b,d) RNA and c,e) DNA tags origin in b-c) ActD treated and d-e) NPM datasets. The inner pie charts represent a broader classification into intergenic and genic (annotated genes), while the outer circles show a finer classification of the genic portion.

**Suppl. Fig. 4. Reproducibility of RADICL-seq libraries among total, ActD and NPM datasets.** Reproducibility of the RNA-DNA interaction frequencies across total (1% FA), ActD and NPM replicates, assessed by counting the occurrences of transcribed genes and 25 kb genomic bins pairs.

**Suppl. Fig. 5. Comparison between RADICL-seq and GRID-seq technologies.** a) RADICL-seq (blue) and GRID-seq (yellow) RNA (left panel) and DNA (right panel) tags distribution across biotypes and genic features. b-d) RADICL-seq (blue) and GRID-seq (yellow) tags density around H3K36me3 (b), H3K4me3 (c) and H3K9ac (d) ChIP-seq peaks. The red line represents the density of tags around randomly selected positions of the genome. e) Reproducibility of the RNA-DNA interaction frequencies across 1% formaldehyde RADICL-seq and GRID-seq replicates, assessed by counting the occurrences of transcribed genes and 25 kb genomic bins pairs. f) RADICL-seq (red) and RAP (blue) of Malat1 RNA-chromatin interactions density.

**Suppl. Fig. 6. Comparison between raw and significant datasets.** a) Interaction counts distribution before (raw; red) and after the significance calculation and filtering (significant; blue). b) Summary of the number and types of interactions for the raw and significant datasets. c) Summary of the number and types of RNA biotypes for the raw and significant datasets.

**Suppl. Fig. 7. Features of RADICL-seq libraries from mOPCs.** a) Summary statistics of the 1% formaldehyde mOPC RADICL-seq replicates sequencing results. b) Reproducibility of the RNA-DNA interaction frequencies across 1% formaldehyde replicates in mESCs and mOPCs, assessed by counting the occurrences of transcribed genes and 25 kb genomic bins pairs. c) RNA and d) DNA tags origin in mOPC. The inner pie charts represent a broader classification into intergenic and genic (annotated genes), while the outer circles show a finer classification of the genic portion.

**Suppl. Fig. 8. Features of RADICL-seq libraries from NPM dataset in mOPCs.** a) Summary statistics of the mOPC RADICL-seq NPM replicates sequencing results. b) Reproducibility of the RNA-DNA interaction frequencies across 1% formaldehyde and NPM replicates in mOPCs, assessed by counting the occurrences of transcribed genes and 25 kb genomic bins pairs. c) RNA and d) DNA tags origin in mOPC NPM samples. The inner pie charts represent a broader classification into intergenic and genic (annotated genes), while the outer circles show a finer classification of the genic portion.

**Suppl. Fig. 9. Chromosome-wide binding of selected transcripts.** a) mESC and b) mOPC RADICL-seq heatmaps showing the percentage of loci covered on each chromosome by the most interacting (>=10% in at least one chromosome) RNAs.

**Suppl. Fig. 10. Genome-wide RNA-chromatin features of RADICL-seq libraries from ActD and NPM datasets in mESCs and mOPCs.** a-b) RNA-DNA interactions shown as a single point per 25 kb bins and coloured by the most represented RNA class or region in that bin for ActD-treated mESCs. c) RNA-DNA interactions quantified for genomic distance between RNA and DNA tags for ActD-treated mESCs. d-e) RNA-DNA interaction matrix for NPM mESCs similar to a-b. f) RNA-DNA interactions quantified for genomic distance between RNA and DNA tags for NPM mESCs. g-h) RNA-DNA interaction matrix for NPM mOPCs similar to a-b. i) RNA-DNA interactions quantified for genomic distance between RNA and DNA tags for NPM mOPCs.

**Suppl. Fig. 11. Characteristics of RADICL-seq libraries from NPM datasets.** a) Distribution of the linear genomic distance between RNA and DNA tags derived from the same read for all datasets. b) Density of the linear genomic distance between RNA and DNA for tags located at less than 1 kb from each other. c) Metadata profiles showing the average coverage of RADICL-seq DNA tags at the peaks defined by DRIP-seq signal in mESCs for total and NPM conditions. d, f, h) Number of the triplexes formed by various regions of Malat1 and Meg3 transcripts in NPM set trans-contacts. Orange dash lines correspond to significant DNA binding domains (DBD). e, g, i) Number of the DNA binding sites involved in triplex formation in DNA contacts (orange dot) and in the random background DNA set (box-plot, 1000 permutations). Only DBD with a significant overrepresentation of triplexes over the background are shown.

**Suppl. Fig. 12. Comparison between RADICL-seq and Hi-C in mESCs and mOPCs.** Spearman correlation (ρ) of HOMER ‘simpleNorm’ normalized Hi-C vs. significant RADICL-seq 1% FA dataset contact frequencies for mESCs (left) and mOPCs (right) at 25 kb resolution.

**Suppl. Fig. 13. Frequency of RNA-DNA interactions at TADs in different dataset in mESCs.** a-b) Metadata profiles showing the average DNA tag coverage at TAD boundaries in mESCs for the 2FA and ActD conditions. c-d) Metadata profiles showing the average RNA tag coverage at TAD boundaries in mESCs for the 2FA and ActD conditions. e) Average DRIP-seq signal at TAD boundaries in mESCs.

**Suppl. Fig. 14. Frequency of REs in RADICL-seq libraries from mESCs and mOPCs.** Proportion of significant RNA-DNA interactions in RADICL-seq mESCs and mOPCs libraries with RNA tags intersecting or non-intersecting RepeatMasker elements. The table displays the number of interactions shown in the plot.

**Suppl. Fig. 15. Distribution of RE families in RADICL-seq libraries from mESCs and mOPCs.** Representation of main repeats families for RNA-DNA pairs where the RNA intersects with RepeatMasker elements contained within GENCODE vM14 annotations for RADICL-seq mESCs and mOPCs libraries. snRNA for GRID-seq replicas were <1% and therefore values were not plotted. Only non-self interactions were used for this plot. The table displays the count of the interactions used for the plot.

**Suppl. Fig. 16. Patterns of RNA-DNA distance interval for RE-mediated interactions in ActD condition.** Distribution of RNA-DNA pairs distance intervals for RNA tags intersecting RepeatMasker annotations paired with DNA tags intersecting GENCODE annotations in RADICL-seq mESCs libraries upon ActD treatment. Only RNA tags intersecting RepeatMasker elements contained within GENCODE vM14 features were used for this analysis. Statistical significance was calculated with two-tailed t-test. * = p≤0.05; ** = p≤0.01; *** = p≤0.001

**Suppl. Fig. 17. Cell type-specific patterns of interactions.** a-b) Circos plots depicting Hunk genomic interactions in mESC and mOPC. c-d) Circos plots depicting Ralgapa2 genomic interactions in mESC and mOPC. Each line represents the interaction between the RNA gene and the contacted DNA locus, while its color highlight the occurrence of the association. The chromosome of origin of the RNA under investigation is shown enlarged on the left portion of each circos plot. e) The distribution of Jaccard distances for genome-wide RNA-DNA binding profiles between cell types for deciles of difference in gene expression between cell types. For genes with a difference in gene expression between cell types < 40%, there is no significant difference in the distribution of Jaccard distances for genome-wide RNA-DNA binding profiles.

**Suppl. Fig. 18. RNA-DNA interactions at promoter regions.** a-b) Distribution of CAGE-derived promoters (top 100 ranked by number of unique interacting RNAs) per chromosome in a) mESCs and b) mOPCs. c) Distribution of unique interacting RNAs on expressed (>=3TPM; yellow) and not expressed (<3TPM; blue) CAGE-derived promoters (+/−2 kb) in mESCs and d) mOPCs.

**Suppl. Fig. 19. Characterization of Neat1-mediated RNA-chromatin interactions.** a) Genomic view of the density of Neat1 RNA tags across its gene body in 1% FA mOPCs (top) and mESCs (bottom). b-c) Density of Neat1 mESCs and mOPCs target genes DNA tags across their gene bodies. Position is reported as percentage of the gene body length.

**Suppl. Fig. 20. Comparison of mESCs and mOPCs *cis*- and *trans*-interactions.** Interactions overlaps between mESC (red) and mOPC (blue) at less than 10kb, 100kb, 1Mb and more than 1Mb, 10Mb in linear distance between RNA and DNA tags or interacting in *trans*.

**Suppl. Table 1. Analysis of lncRNA distance interaction patterns.** For each pair of RNA and DNA distance interval the number of unique lncRNAs and the number and fraction of all lncRNA-mediated interactions are reported for both mESCs and mOPCs datasets.

**Suppl. Table 2. Top ten interacting transcripts for *cis-* and *trans-*contacts.** Only lncRNAs and protein-coding transcripts are listed for each condition.

**Suppl. Table 3. List of cell type-specific markers.**

**Suppl. Table 4. Shared *trans*-interacting long noncoding RNAs captured in both mESCs and mOPCs.** For each transcript, the numbers of shared interactions in each cell type are shown.

