## Supplementary figures and tables for "RADICL-seq identifies general and cell type-specific principles of genome-wide RNA-chromatin interactions"

a

| Enzymatic treatment | Raw reads | rRNA-containing (% of input) | Uniquely mapped (% of input) |
|---|---|---|---|
| No treatment | 2.2 M | 0.8 M (39%) | 0.3 M (14%) |
| Nuclease S1 | 2.2 M | 0.9 M (43%) | 0.4 M (16%) |
| RNase V1 | 2.9 M | 1.4 M (49%) | 0.5 M (17%) |
| RNase H | 3 M | 0.9 M (30%) | 0.8 M (26%) |

b

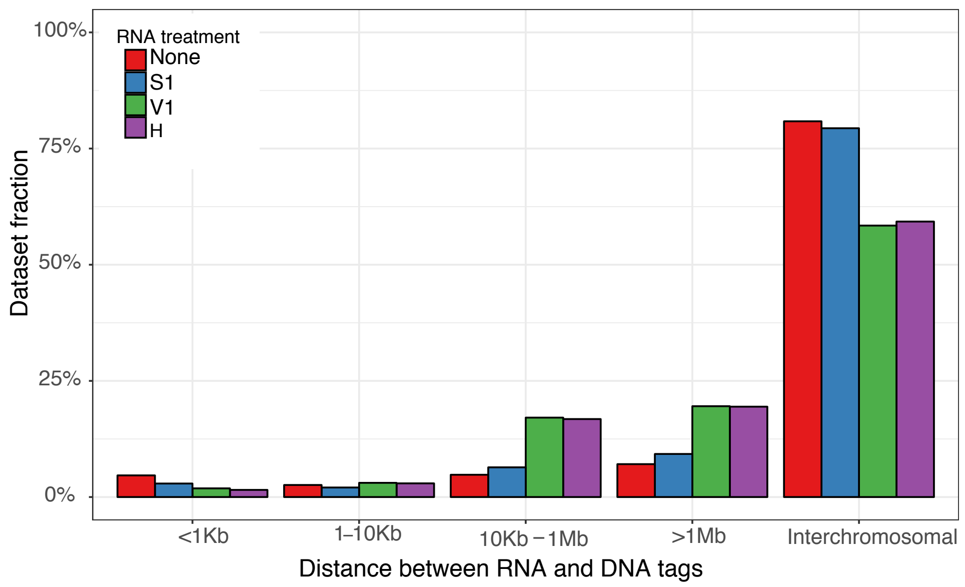


c

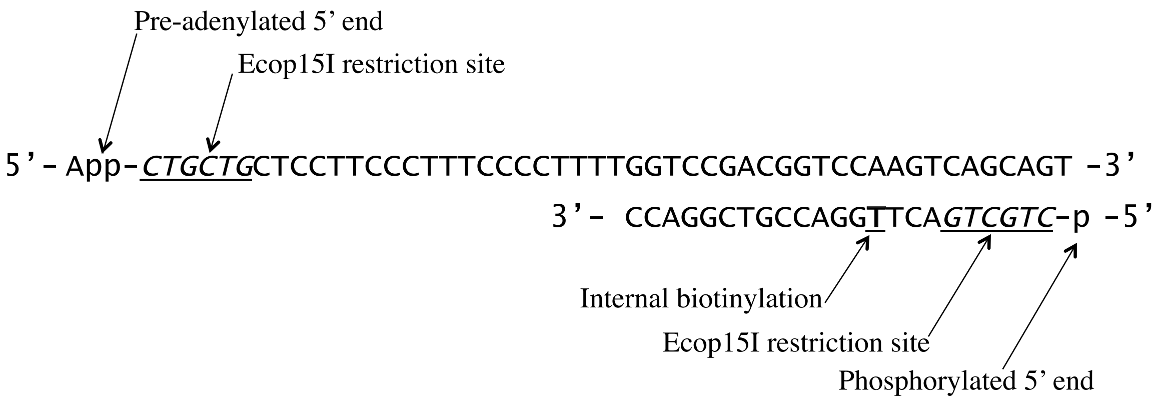


d

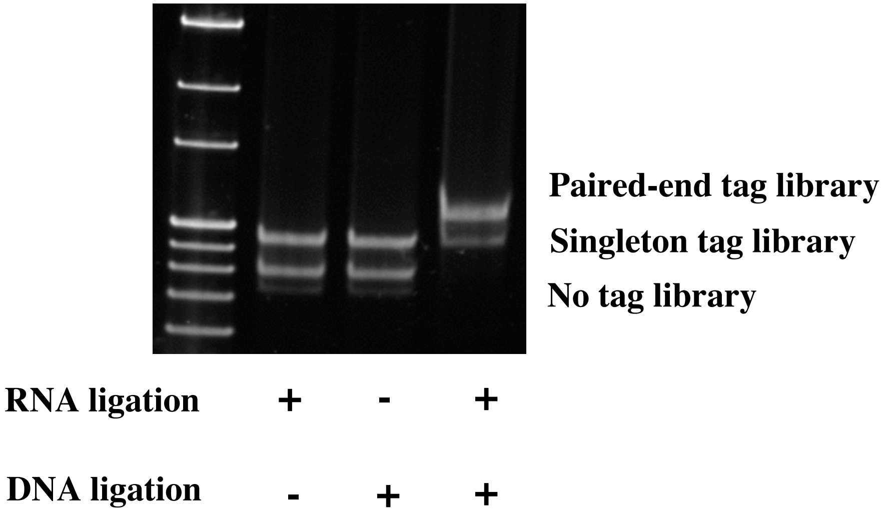


**Suppl. Fig. 1. Characterization of RADICL-seq technology.** a) Summary statistics of the sequencing outcome of the samples after testing different enzymatic treatments. b) Distribution of the linear genomic distance between RNA and DNA tags derived from the same read after treatment with different enzymes. c) Sequence and features of the bridge adapter. The adapter contains a 5' adenylated end (App), an internal T residue with biotin modification (bold) and two restriction sites for EcoP15I. The 3'-overhanging T allows DNA ligation with dA-tailed genomic fragments. d). Generation of RADICL-seq libraries and controls. Gel migration pattern of PCR-amplified RADICL-seq library compared with controls generated by omitting DNA ligase or RNA ligase. The expected band was detected only when both RNA and DNA ligases were used for the construction of the library.

**a**

| Experimental condition | Biological replicate | Raw reads | rRNA | Multi-mapping | Uniquely mapped |
|---|---|---|---|---|---|
| 1FA | n1 | 141.3M | 42.1M | 20.4M | 18.2M |
| 1FA | n2 | 120.5M | 46.9M | 27.9M | 21.9M |
| 1FA | n3 | 103.8M | 37.5M | 23.3M | 19.7M |
| 2FA | n1 | 141.8M | 31.4M | 17.9M | 14.9M |
| 2FA | n2 | 107.4M | 44.5M | 19.8M | 15.8M |
| 2FA | n3 | 97.6 M | 26.9M | 20.9M | 17.3M |

**b**

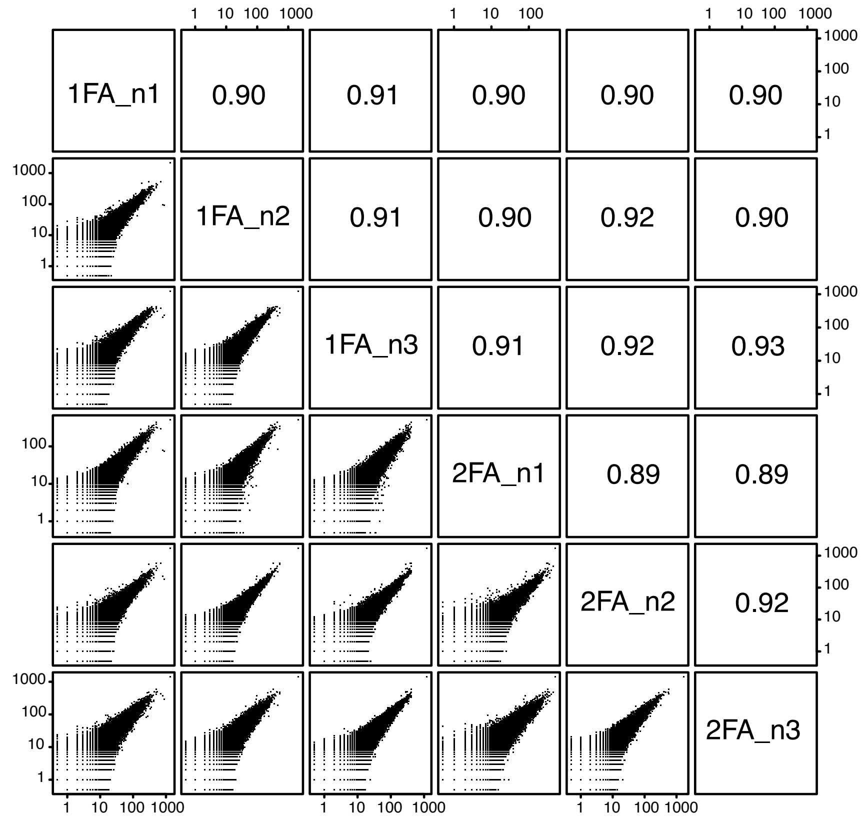


**c**

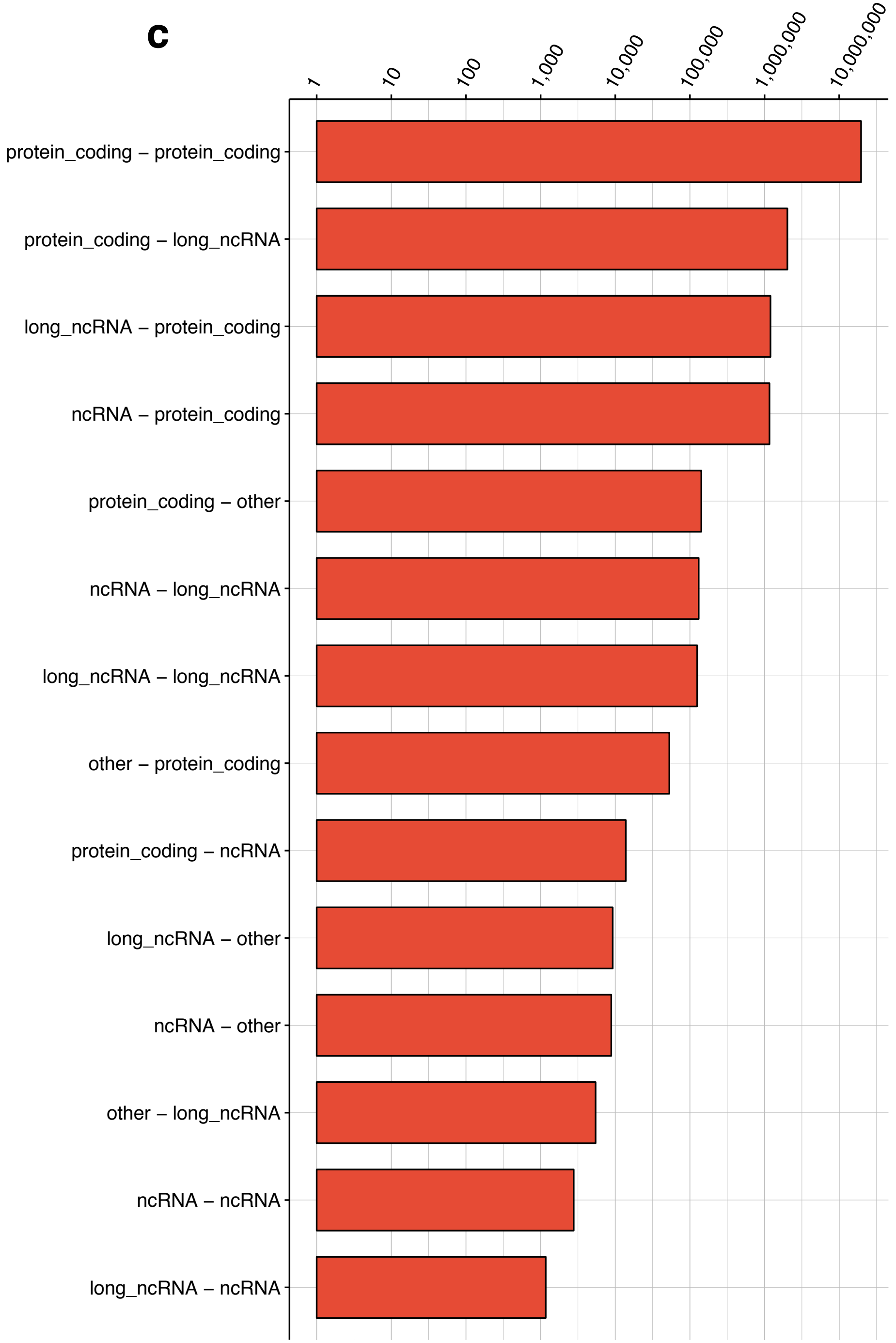


**Suppl. Fig. 2. Features of RADICL-seq libraries.** a) Summary statistics of the 1% and 2% formaldehyde treatments and replicates sequencing results. b) Reproducibility of the RNA-DNA interaction frequencies across formaldehyde treatments and replicates, assessed by counting the occurrences of transcribed genes and 25 kb genomic bins pairs. c) Distribution of RNA and DNA tag biotypes according to their mapped loci and gene annotation.

a

| Experimental condition | Biological replicate | Raw reads | rRNA-containing read pairs | Multi-mapping read pairs | Uniquely mapped read pairs |
|---|---|---|---|---|---|
| ActD-treated | n1 | 80.8 M | 34 M | 12.2 M | 9.3 M |
| ActD-treated | n2 | 46.4 M | 19.7 M | 7 M | 5.4 M |
| Non protein-mediated | n1 | 185.4 M | 86.4 M | 28.9 M | 14.2 M |
| Non protein-mediated | n2 | 101.2 M | 50.3 M | 17.6 M | 13.9 M |

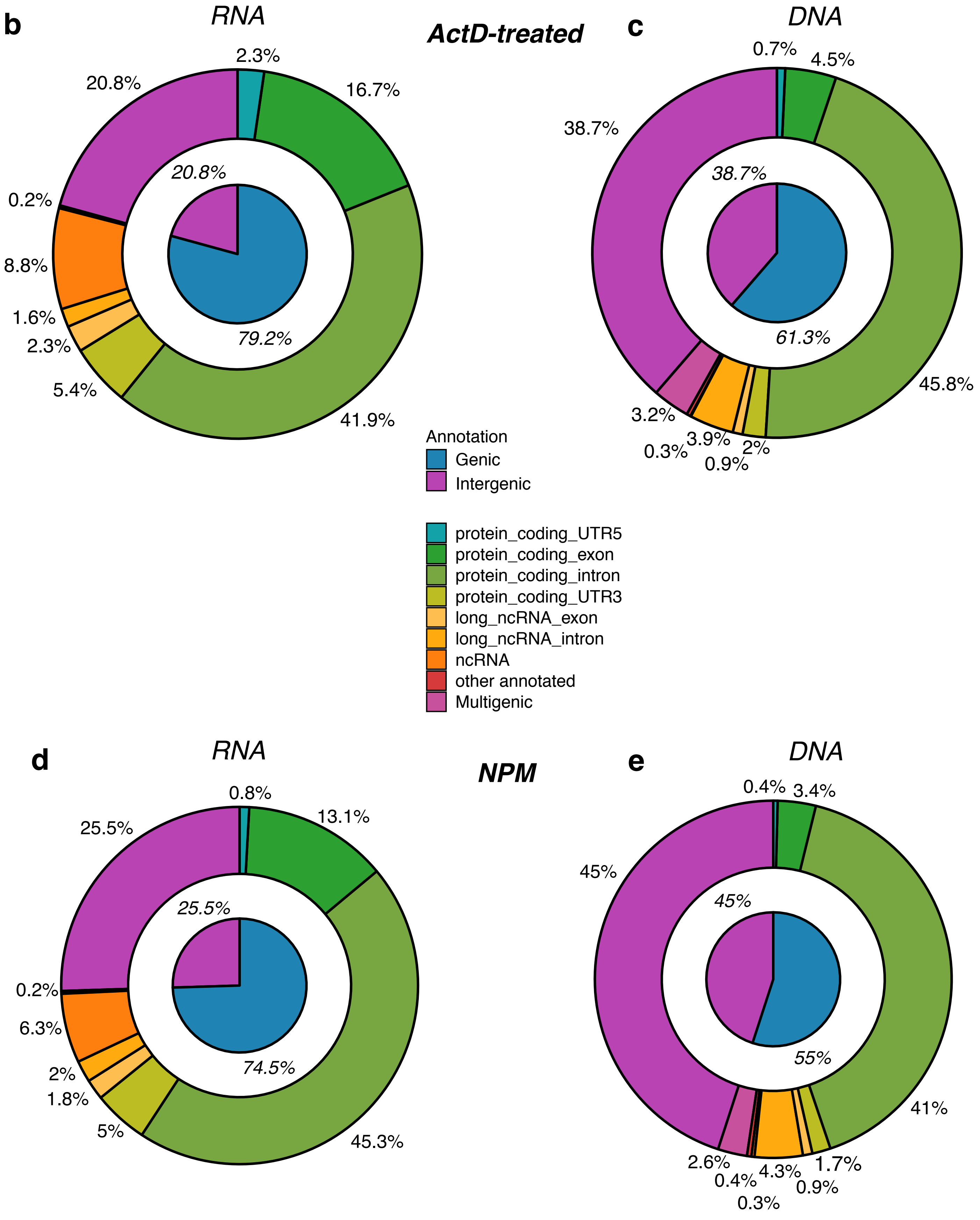


**Suppl. Fig. 3. Features of RADICL-seq libraries from ActD and NPM datasets.** a) Summary statistics of the ActD treated and NPM samples and replicates sequencing results. b,d) RNA and c,e) DNA tags origin in b-c) ActD treated and d-e) control datasets. The inner pie charts represent a broader classification into intergenic and genic (annotated genes), while the outer circles show a finer classification of the genic portion.

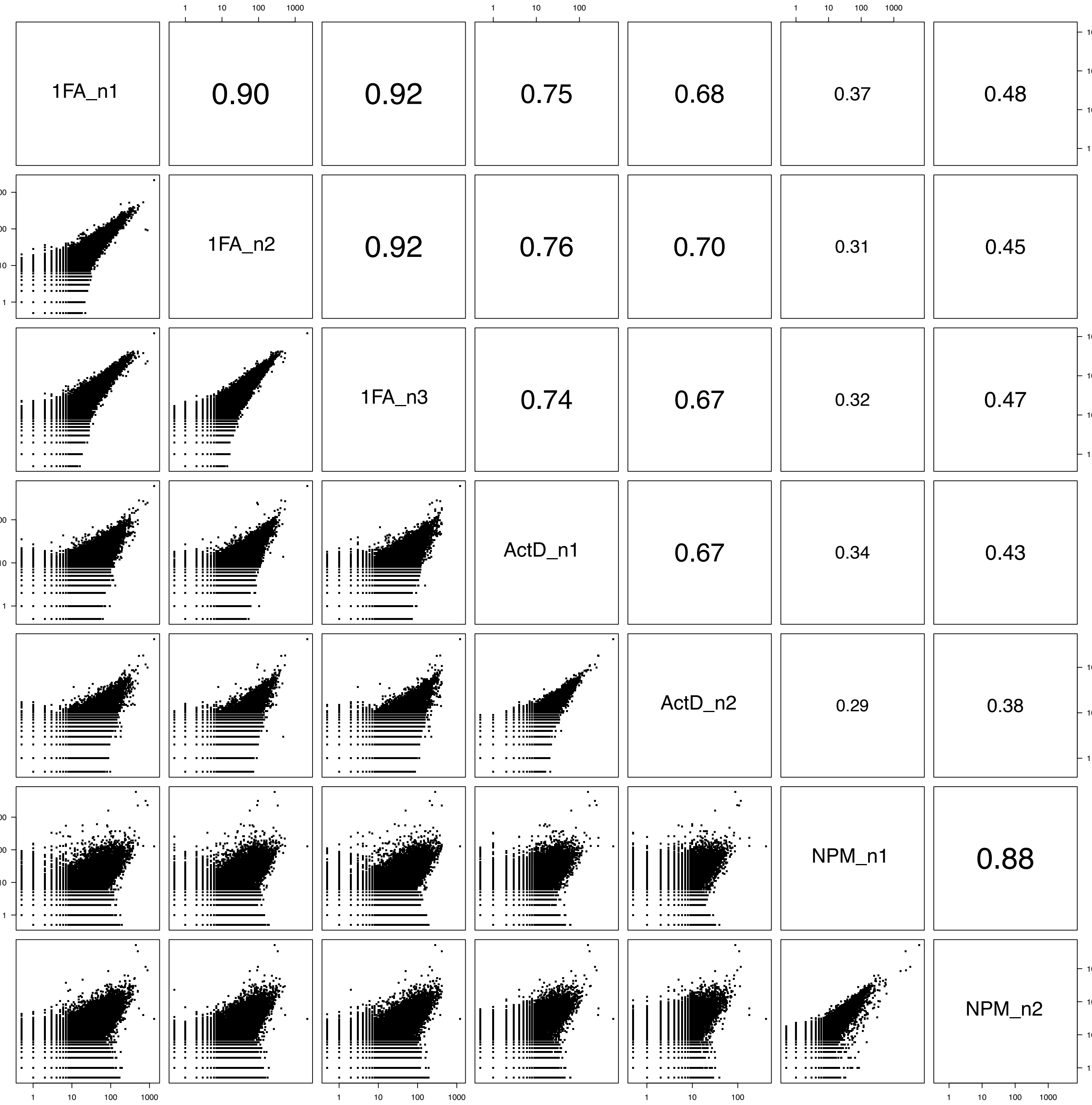


**Suppl. Fig. 4. Reproducibility of RADICL-seq libraries among total, ActD and NPM datasets.** Reproducibility of the RNA-DNA interaction frequencies across 1% formaldehyde, ActD and NPM replicates, assessed by counting the occurrences of transcribed genes and 25 kb genomic bins pairs.

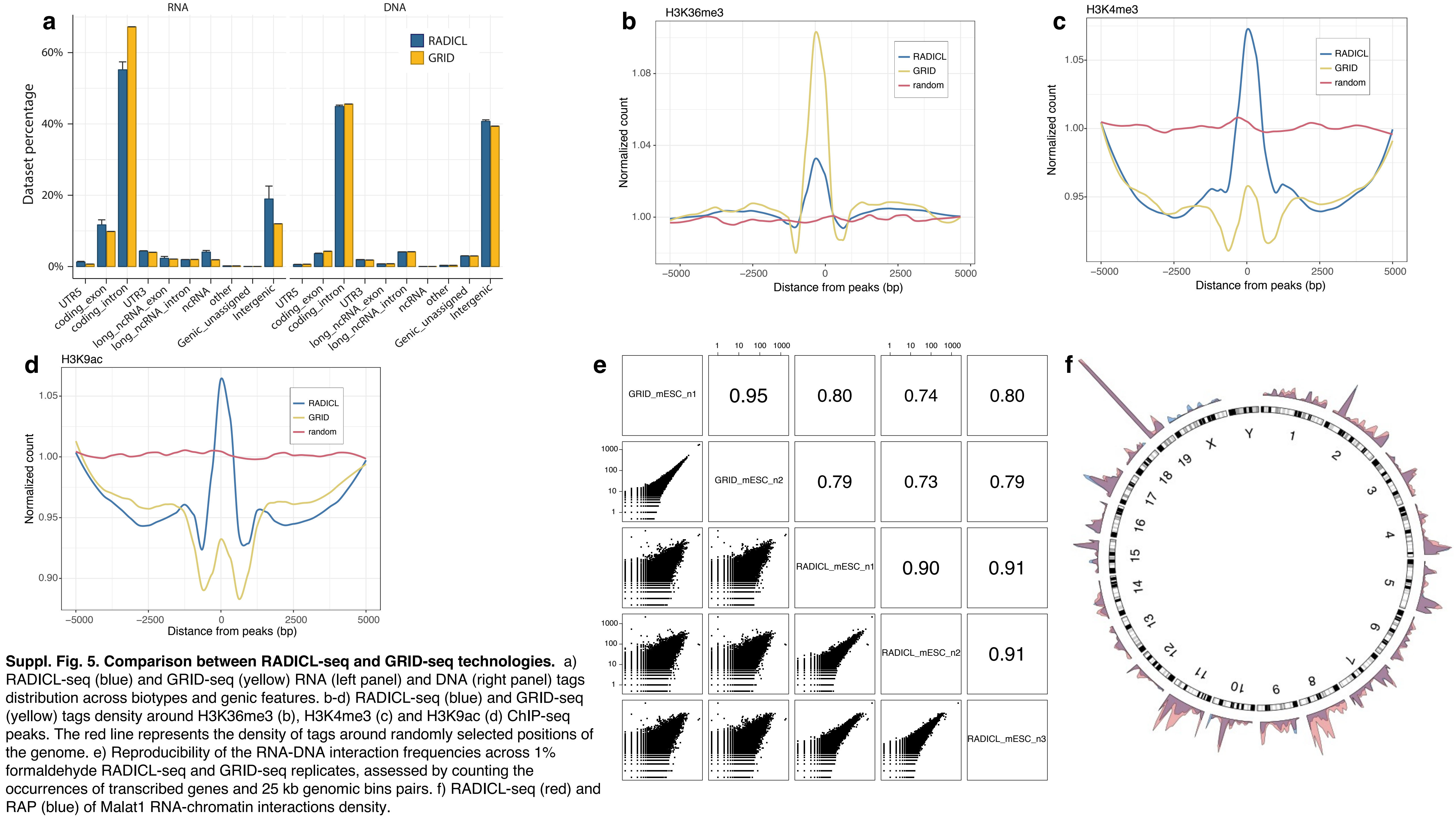


**Suppl. Fig. 5. Comparison between RADICL-seq and GRID-seq technologies.** a) RADICL-seq (blue) and GRID-seq (yellow) RNA (left panel) and DNA (right panel) tags distribution across biotypes and genic features. b-d) RADICL-seq (blue) and GRID-seq (yellow) tags density around H3K36me3 (b), H3K4me3 (c) and H3K9ac (d) ChIP-seq peaks. The red line represents the density of tags around randomly selected positions of the genome. e) Reproducibility of the RNA-DNA interaction frequencies across 1% formaldehyde RADICL-seq and GRID-seq replicates, assessed by counting the occurrences of transcribed genes and 25 kb genomic bins pairs. f) RADICL-seq (red) and RAP (blue) of Malat1 RNA-chromatin interactions density.

**a**

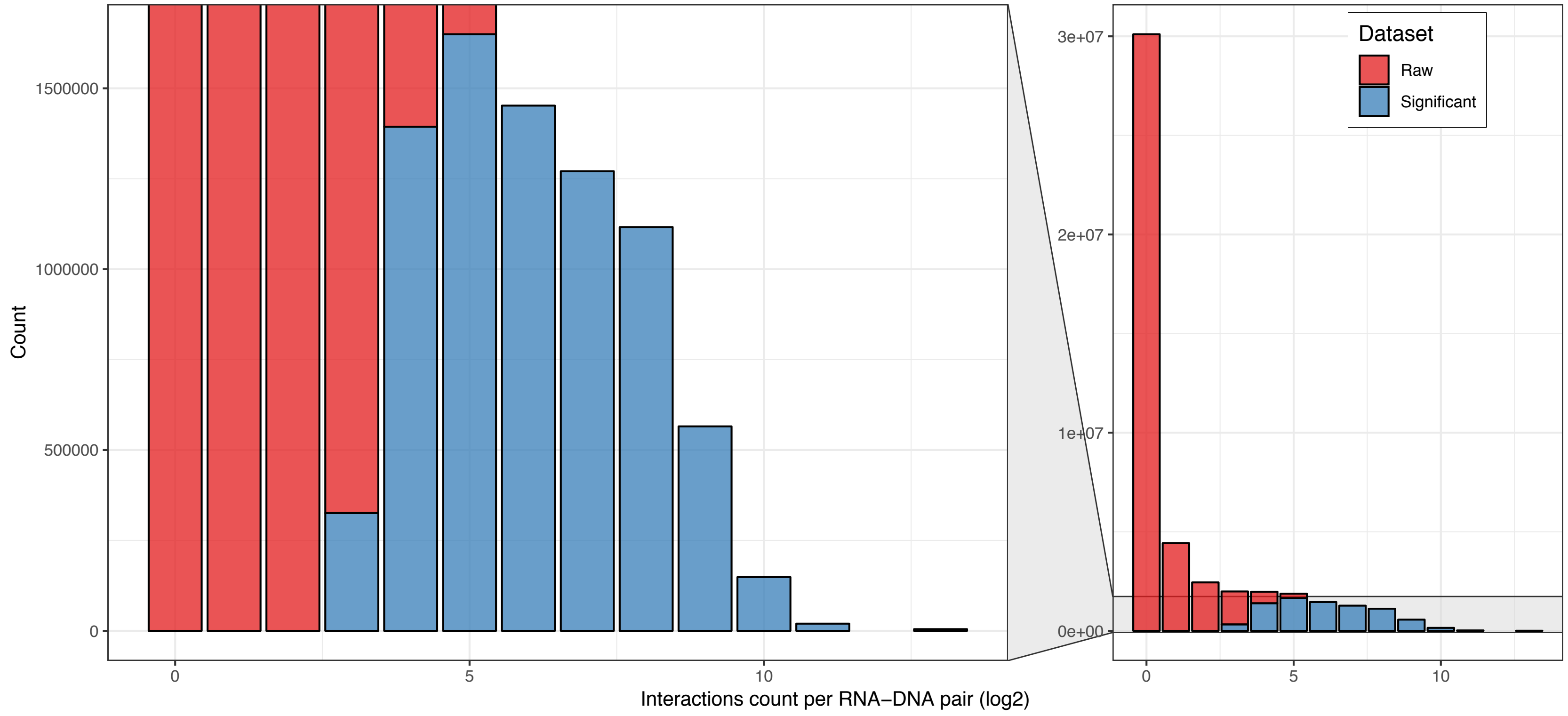


**b**

| Experimental Condition | Raw dataset: Total unique interactions | Raw dataset: Cis *(% total)* | Raw dataset: Trans *(% total)* | Significant dataset: Total unique interactions | Significant dataset: Cis *(% total)* | Significant dataset: Trans *(% total)* |
|---|---|---|---|---|---|---|
| 1FA | 49,914,160 | 17,523,875 *(35.1%)* | 32,390,285 *(64.9%)* | 8420123 | 7954089 *(94.5%)* | 466,034 *(5.5%)* |
| 2FA | 38,763,078 | 14,630,187 *(37.7%)* | 24,132,891 *(62.3%)* | 7,213,577 | 6,867,871 *(95.2%)* | 345,706 *(4.8%)* |
| ActD | 12,002,693 | 2,936,878 *(24.5%)* | 9,065,815 *(75.5%)* | 724,021 | 664,853 *(91.8%)* | 59,168 *(8.2%)* |
| NPM | 21,489,520 | 2,331,587 *(10.8%)* | 19,157,933 *(89.2%)* | 1,136,020 | 1,129,456 *(99.4%)* | 6,564 *(0.6%)* |

**c**

| Condition | Biotype | Raw dataset | Significant dataset | *Percentage* |
|---|---|---|---|---|
| 1FA | Protein-coding | 20,684 | 12,441 | *60.2%* |
| 1FA | Long_ncRNAs | 10,480 | 1,430 | *13.7%* |
| 1FA | ncRNAs | 2,427 | 81 | *3.3%* |
| 1FA | Other | 5,831 | 49 | *0.8%* |
| 2FA | Protein-coding | 20,664 | 12,521 | *60.6%* |
| 2FA | Long_ncRNAs | 10,485 | 1,536 | *14.7%* |
| 2FA | ncRNAs | 2,375 | 90 | *3.8%* |
| 2FA | Other | 5,772 | 48 | *0.8%* |
| ActD | Protein-coding | 20,034 | 8,053 | *40.2%* |
| ActD | Long_ncRNAs | 9,523 | 279 | *2.9%* |
| ActD | ncRNAs | 1,454 | 30 | *2.1%* |
| ActD | Other | 3,646 | 8 | *0.2%* |
| NPM | Protein-coding | 20,479 | 10,702 | *52.3%* |
| NPM | Long_ncRNAs | 10,248 | 762 | *7.4%* |
| NPM | ncRNAs | 1,950 | 130 | *6.7%* |
| NPM | Other | 4,969 | 33 | *0.7%* |

**Suppl. Fig. 6. Comparison between raw and significant datasets.** a) Interaction counts distribution before (raw; red) and after the significance calculation and filtering (significant; blue). b) Summary of the number and types of interactions for the raw and significant datasets. c) Summary of the number and types of RNA biotypes for the raw and significant datasets.

**a**

| Experimental condition | Biological replicate | Raw reads | rRNA-containing read pairs | Multi-mapping read pairs | Uniquely mapped read pairs |
|---|---|---|---|---|---|
| 1% FA | n1 | 109.6 M | 12.4 M | 15.9 M | 15.4 M |
| 1% FA | n2 | 177.1 M | 23.4 M | 20.2 M | 17.6 M |
| 1% FA | n3 | 116.7 M | 21.1 M | 19.5 M | 16.4 M |

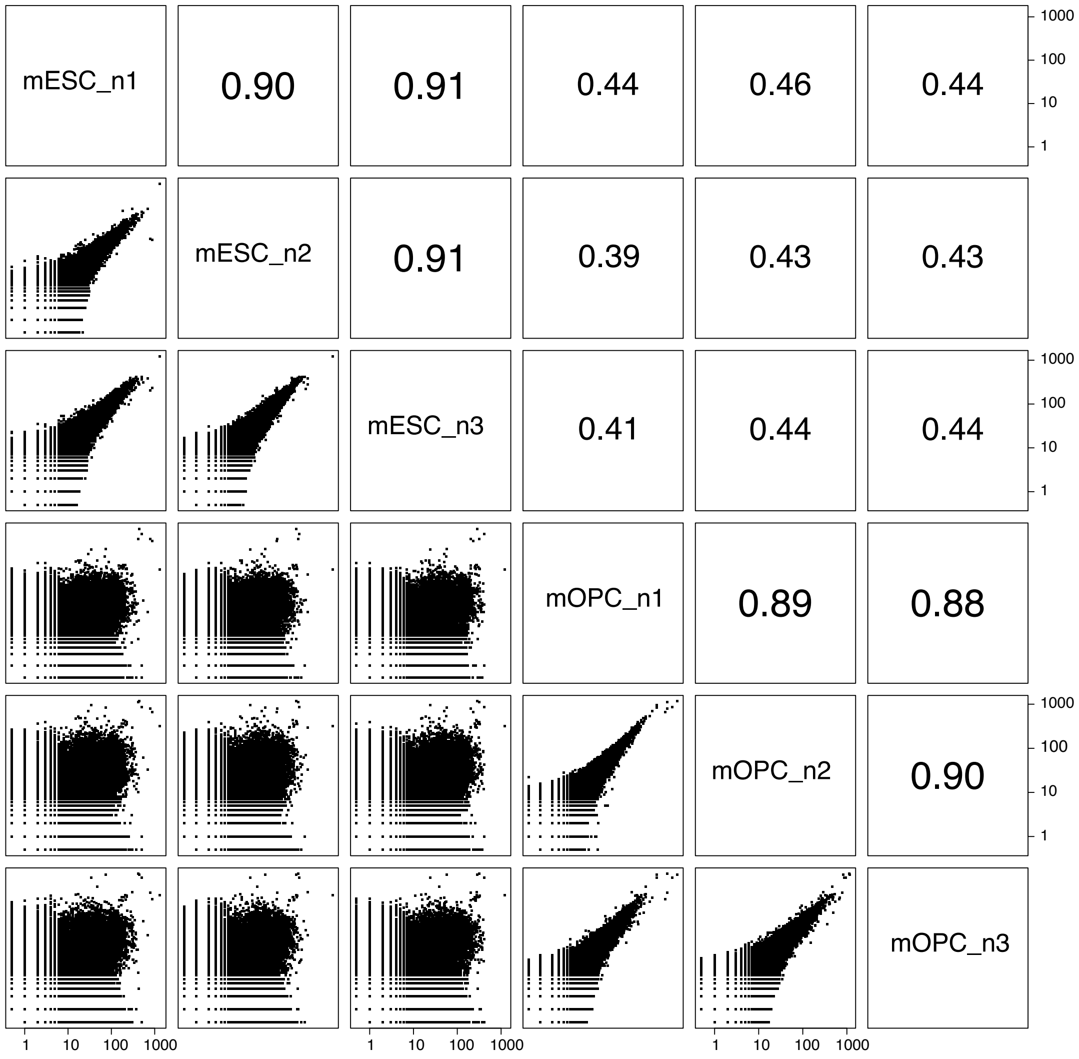


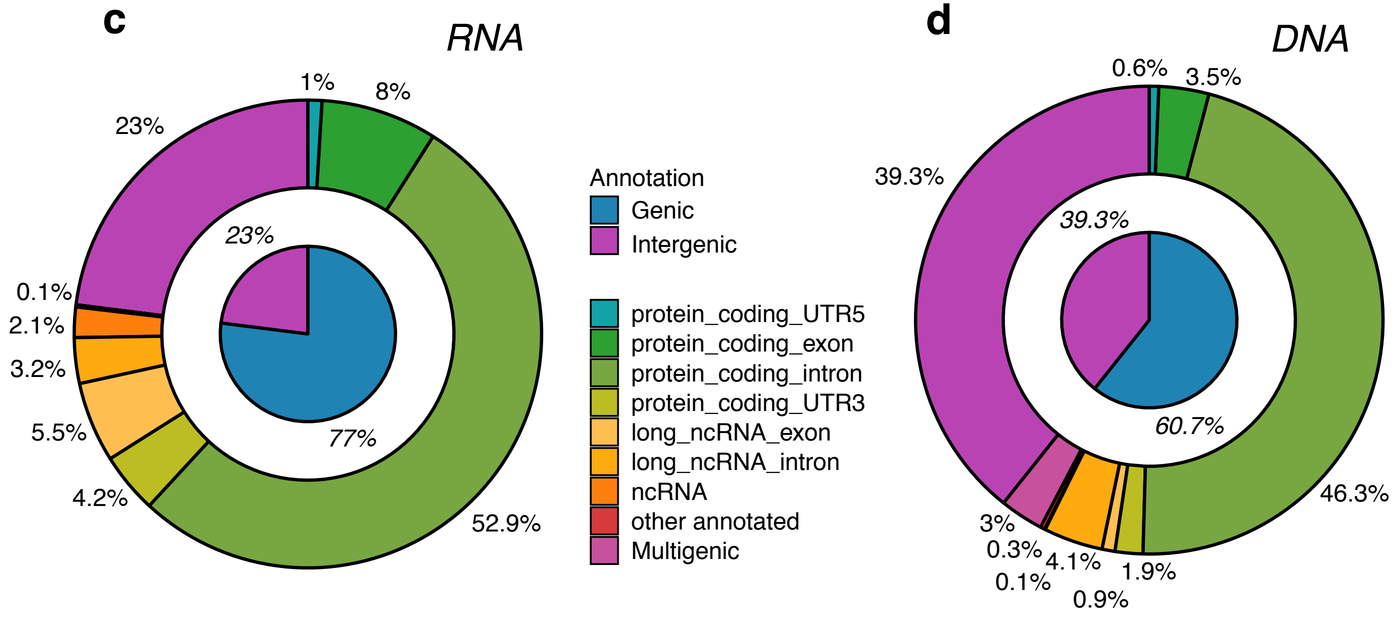


**Suppl. Fig. 7. Features of RADICL-seq libraries from mOPCs.** a) Summary statistics of the 1% formaldehyde mOPC RADICL-seq replicates sequencing results. b) Reproducibility of the RNA-DNA interaction frequencies across 1% formaldehyde replicates in mESCs and mOPCs, assessed by counting the occurrences of transcribed genes and 25 kb genomic bins pairs. c) RNA and d) DNA tags origin in mOPC. The inner pie charts represent a broader classification into intergenic and genic (annotated genes), while the outer circles show a finer classification of the genic portion.

**a**

| Experimental condition | Biological replicate | Raw reads | rRNA-containing read pairs | Multi-mapping read pairs | Uniquely mapped read pairs |
|---|---|---|---|---|---|
| Non protein-mediated | n1 | 183.4 M | 54.2 M | 35.2 M | 24.3 M |
| Non protein-mediated | n2 | 83.2 M | 3.5 M | 29.3 M | 25.5 M |

**b**

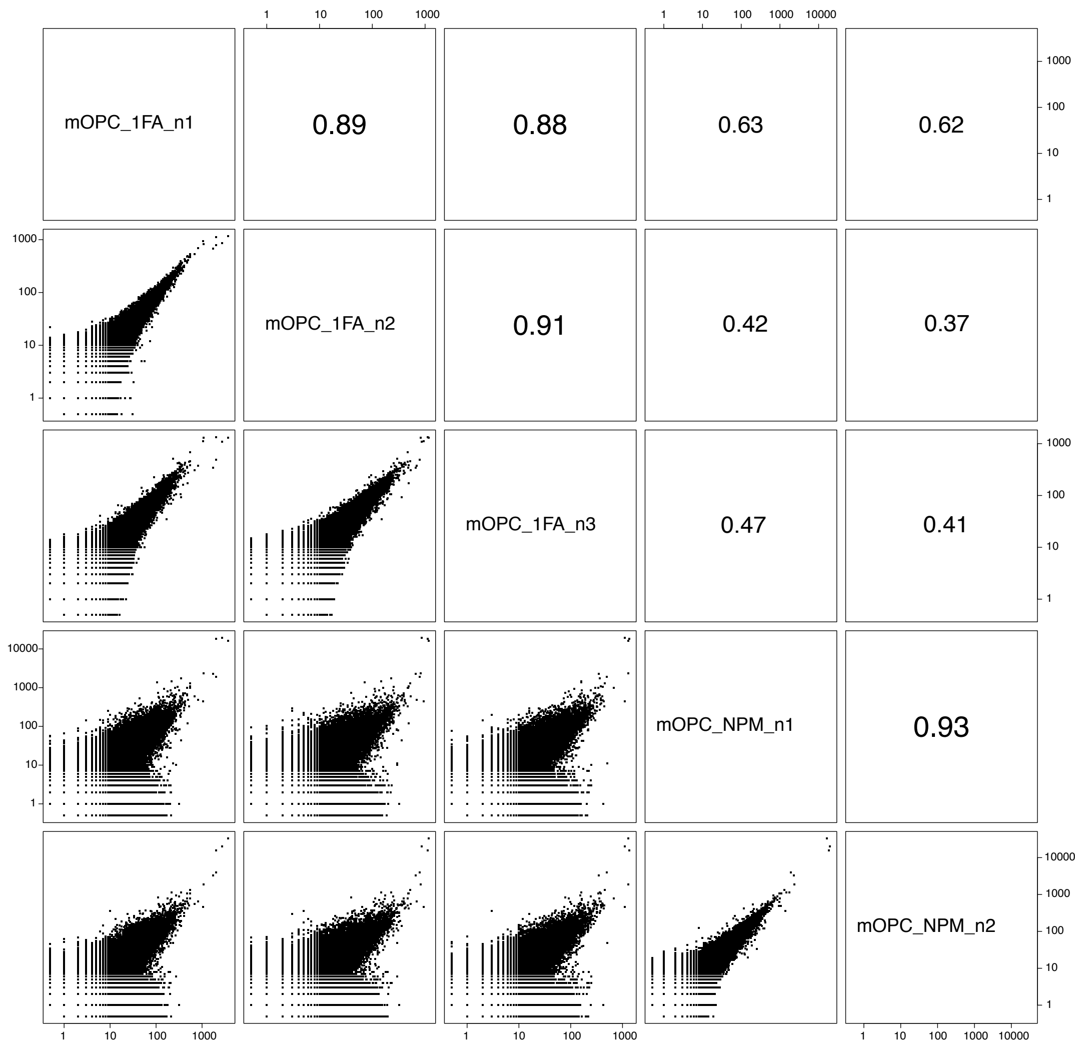


**c**

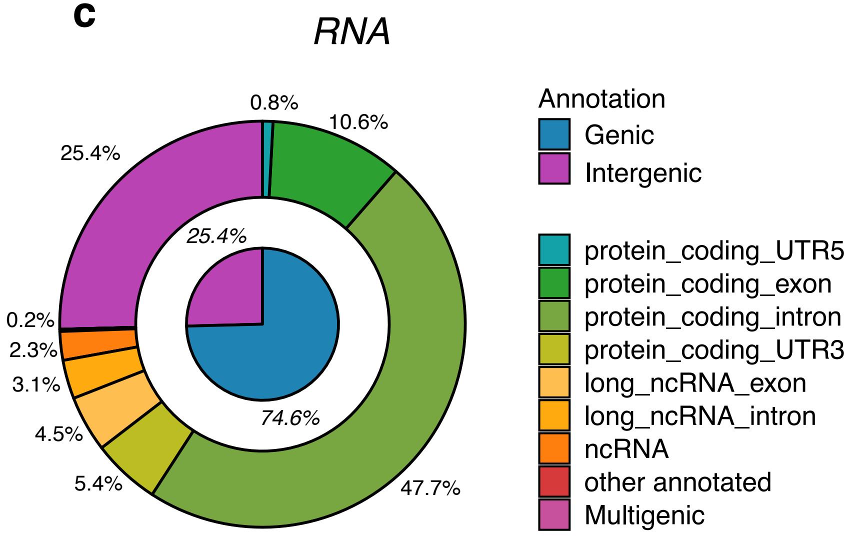


**d**

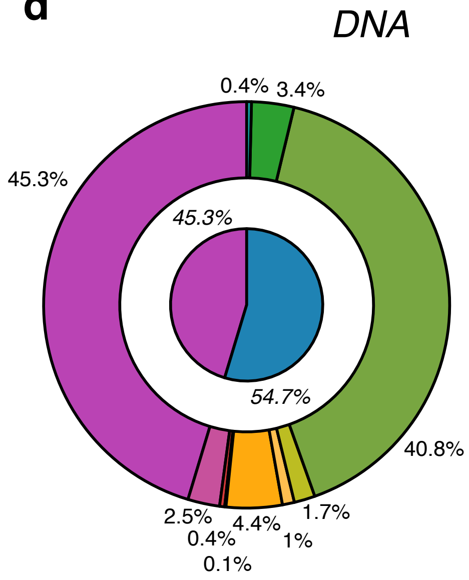


**Suppl. Fig. 8. Features of RADICL-seq libraries from NPM dataset in mOPCs.** a) Summary statistics of the mOPC RADICL-seq NPM replicates sequencing results. b) Reproducibility of the RNA-DNA interaction frequencies across 1% formaldehyde and NPM replicates in mOPCs, assessed by counting the occurrences of transcribed genes and 25 kb genomic bins pairs. c) RNA and d) DNA tags origin in mOPC NPM samples. The inner pie charts represent a broader classification into intergenic and genic (annotated genes), while the outer circles show a finer classification of the genic portion.

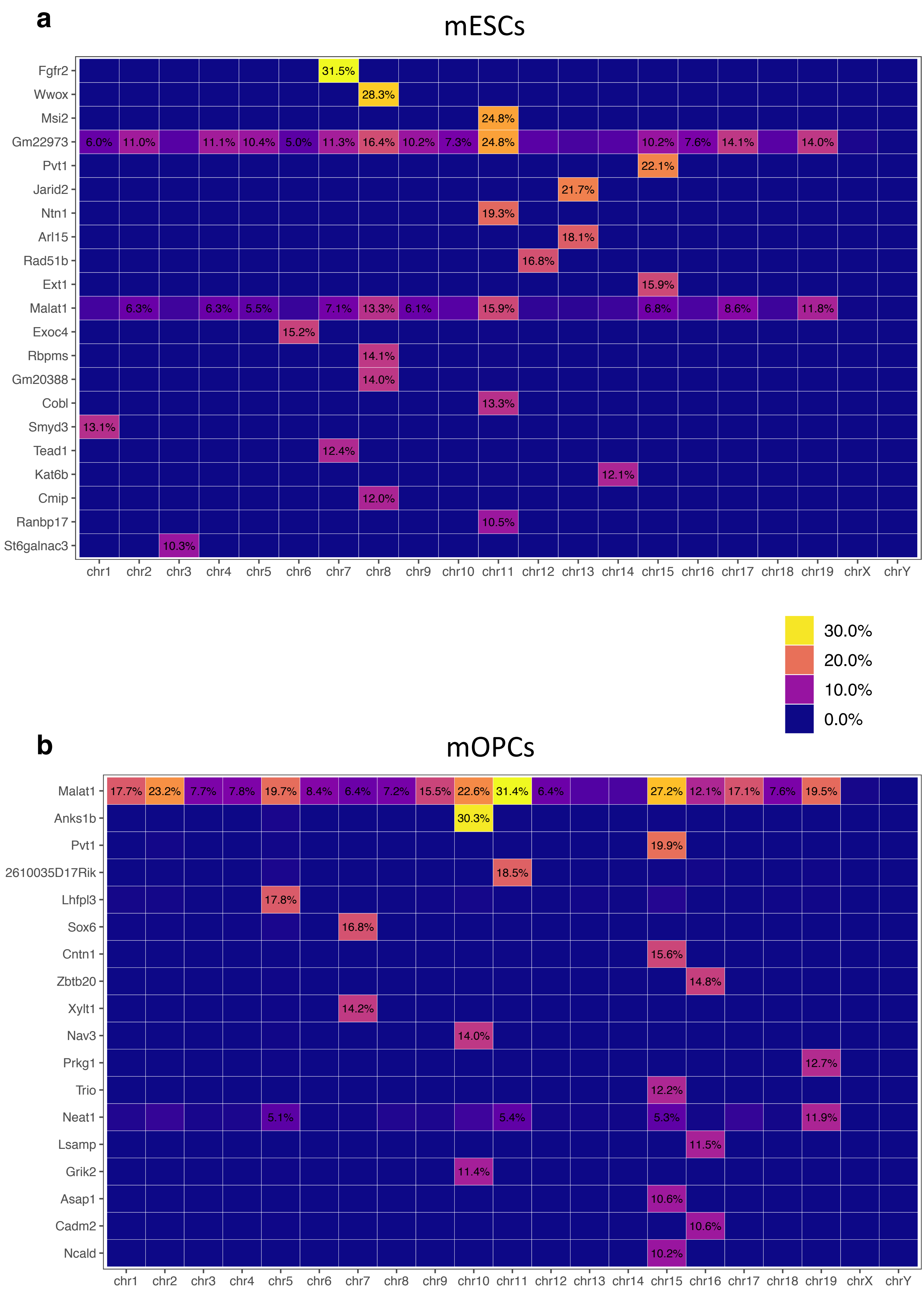


**Suppl. Fig. 9. Chromosome-wide binding of selected transcripts.** a) mESC and b) mOPC RADICL-seq heatmaps showing the percentage of loci covered on each chromosome by the most interacting (>=10% in at least one chromosome) RNAs.

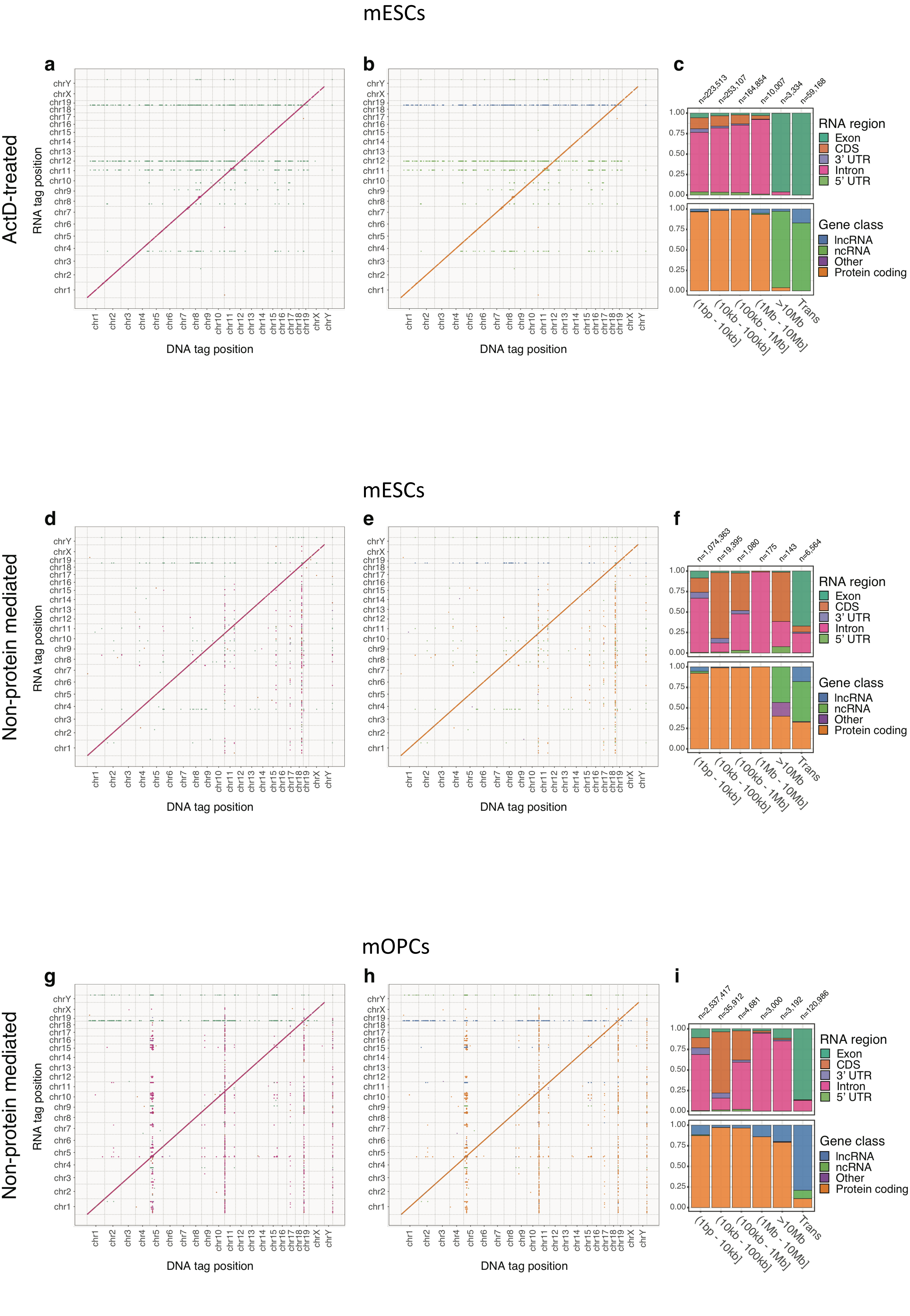


**Suppl. Fig. 10. Genome-wide RNA-chromatin features of RADICL-seq libraries from ActD and NPM datasets in mESCs and mOPCs.** a-b) RNA-DNA interactions shown as a single point per 25 kb bins and coloured by the most represented RNA class or region in that bin for ActD-treated mESCs. c) RNA-DNA interactions quantified for genomic distance between RNA and DNA tags for ActD-treated mESCs. d-e) RNA-DNA interaction matrix for NPM mESCs similar to a-b. f) RNA-DNA interactions quantified for genomic distance between RNA and DNA tags for NPM mESCs. g-h) RNA-DNA interaction matrix for NPM mOPCs similar to a-b. i) RNA-DNA interactions quantified for genomic distance between RNA and DNA tags for NPM mOPCs.

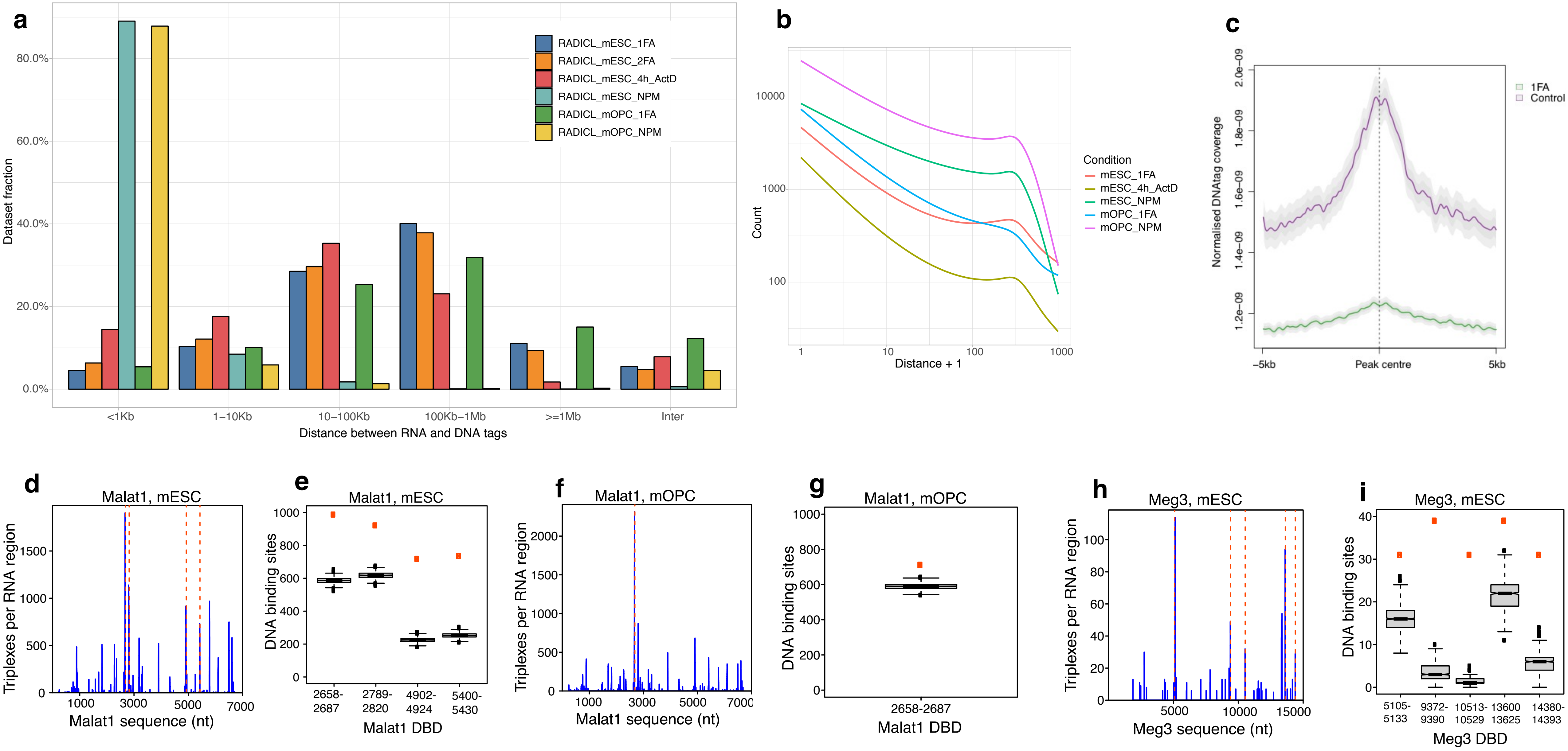


**Suppl. Fig. 11. Characteristics of RADICL-seq libraries from NPM datasets.** a) Distribution of the linear genomic distance between RNA and DNA tags derived from the same read for all datasets. b) Density of the linear genomic distance between RNA and DNA for tags located at less than 1 kb from each other. c) Metadata profiles showing the average coverage of RADICL-seq DNA tags at the peaks defined by DRIP-seq signal in mESCs for total and NPM conditions. d, f, h) Number of the triplexes formed by various regions of Malat1 and Meg3 transcripts in NPM set trans-contacts. Orange dash lines correspond to significant DNA binding domains (DBD). e, g, i) Number of the DNA binding sites involved in triplex formation in DNA contacts (orange dot) and in the random background DNA set (box-plot, 1000 permutations). Only DBD with a significant overrepresentation of triplexes over the background are shown.

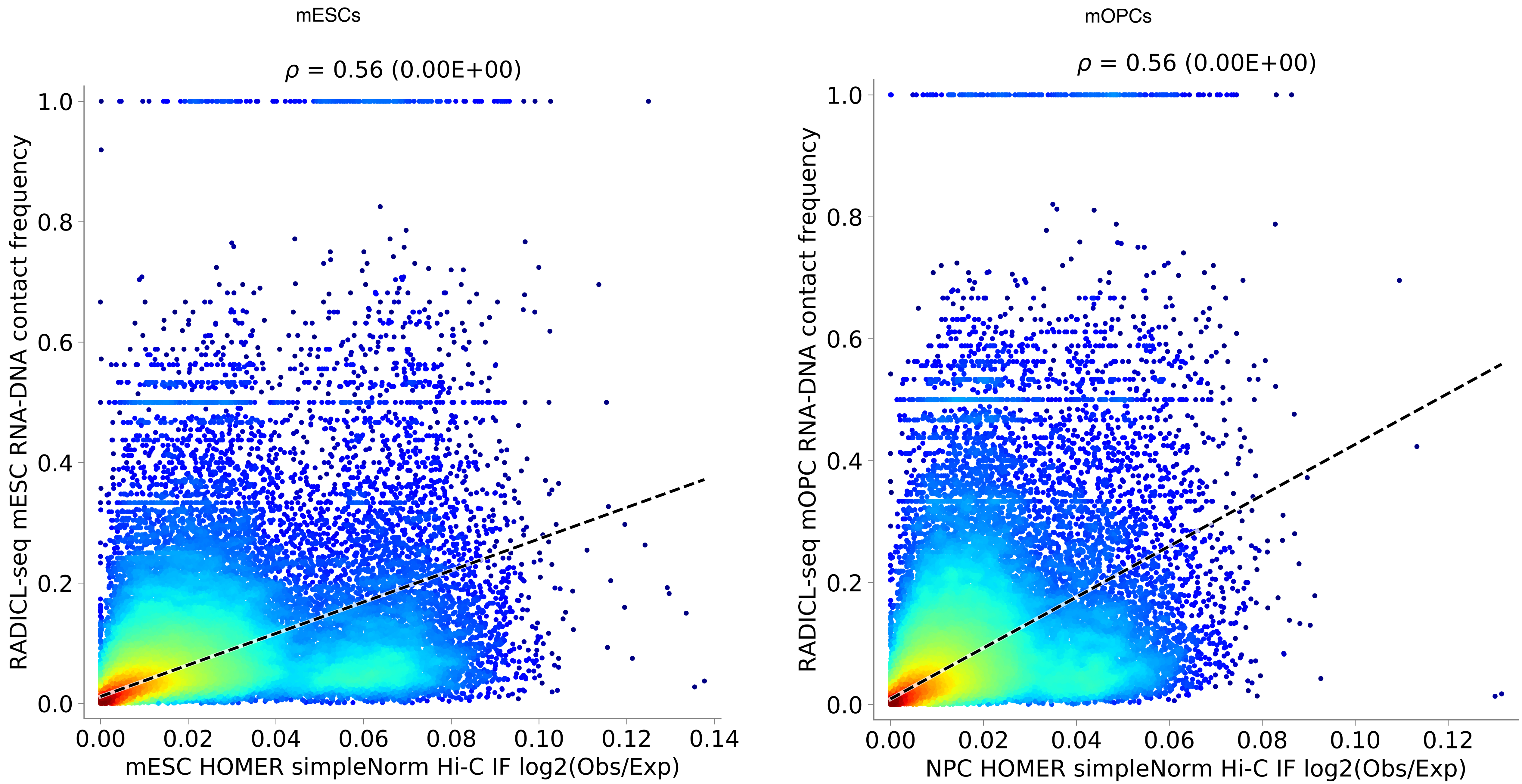


**Suppl. Fig. 12. Comparison between RADICL-seq and Hi-C in mESCs and mOPCs.** Spearman correlation (r) of HOMER 'simpleNorm' normalized Hi-C vs. significant RADICL-seq 1% FA dataset contact frequencies for mESCs (left) and mOPCs (right) at 25 kb resolution.

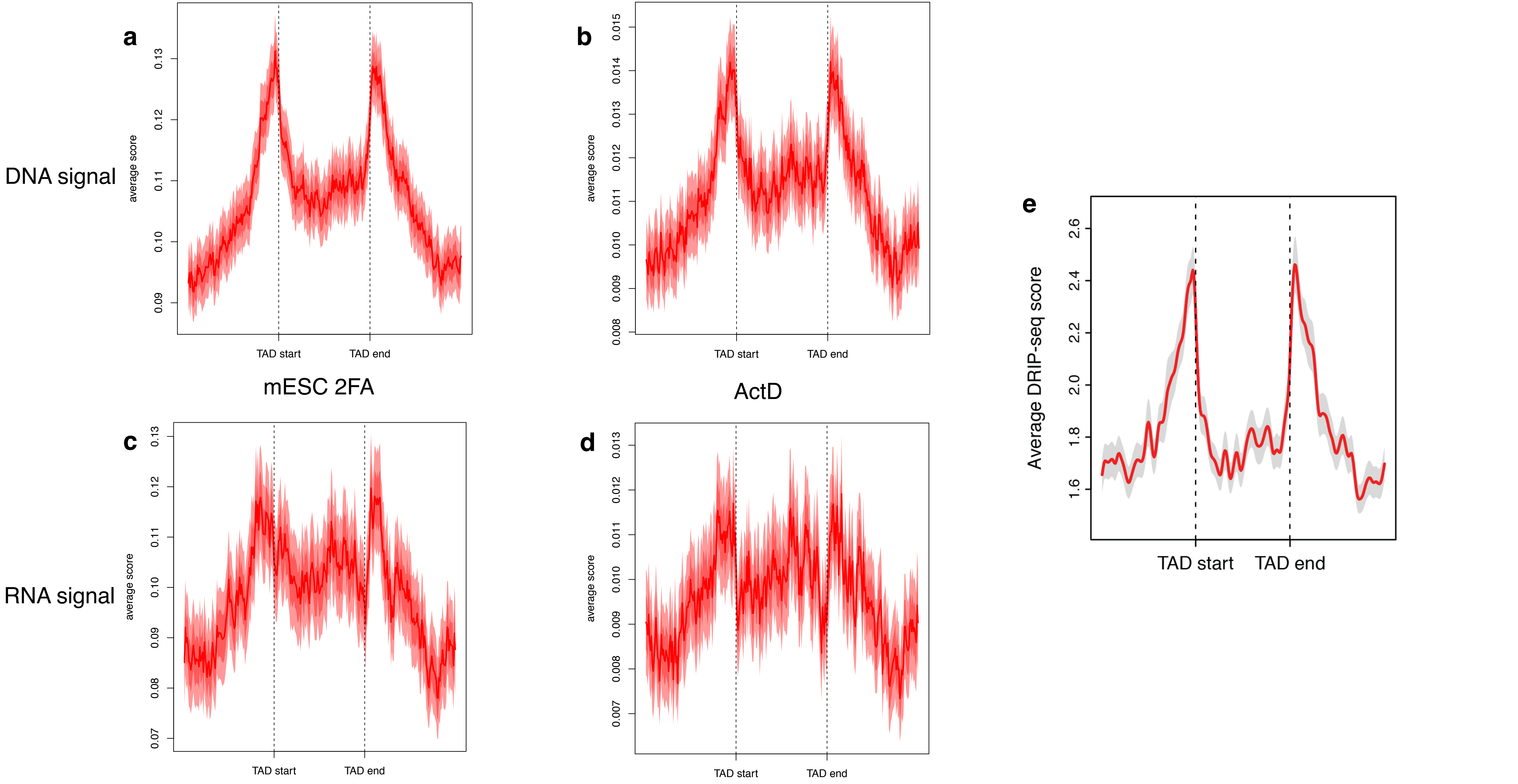


**Suppl. Fig. 13. Frequency of RNA-DNA interactions at TADs in different dataset in mESCs.** a-b) Metadata profiles showing the average DNA tag coverage at TAD boundaries in mESCs for the 2FA and ActD conditions. c-d) Metadata profiles showing the average RNA tag coverage at TAD boundaries in mESCs for the 2FA and ActD conditions. e) Average DRIP-seq signal at TAD boundaries in mESCs.

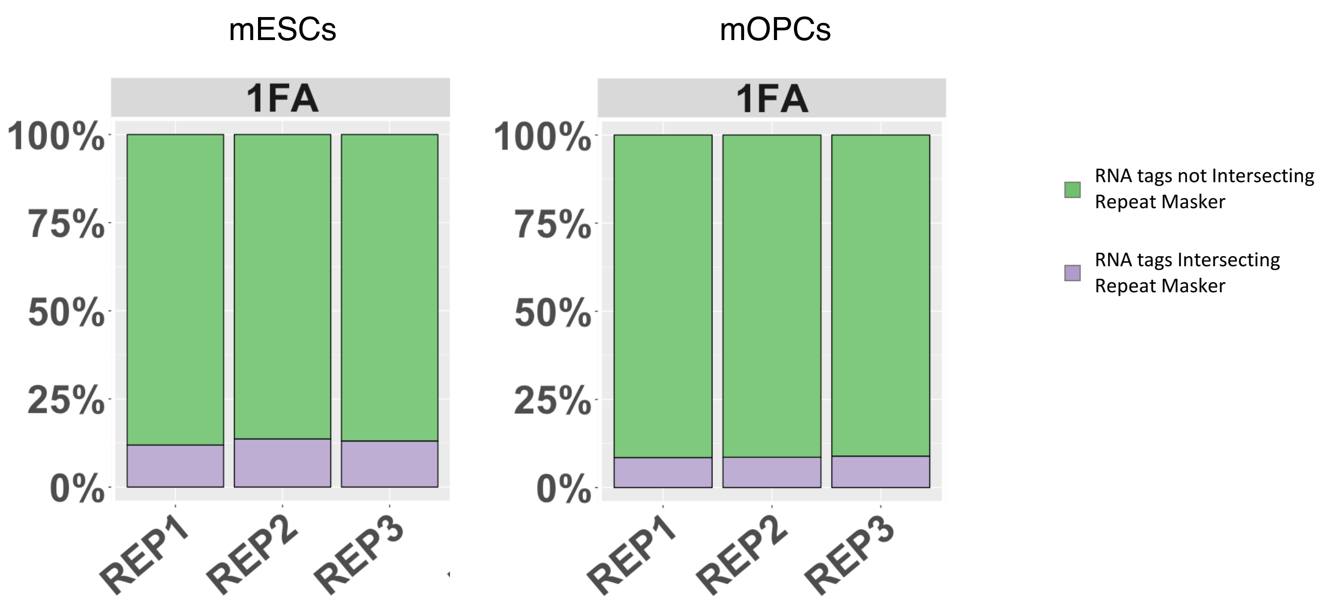


| Condition | Significant Interactions (p<0.05) | RNA tags Not Intersecting Repeat Masker | RNA tags Intersecting Repeat Masker |
|---|---|---|---|
| mESC_1FA-REP1 | 2841875 | 2502970 | 338905 |
| mESC_1FA-REP2 | 2608899 | 2252602 | 356297 |
| mESC_1FA-REP3 | 3014022 | 2619960 | 394062 |
| mOPC_1FA-REP1 | 2378935 | 2176159 | 202776 |
| mOPC_1FA-REP2 | 2261016 | 2066741 | 194275 |
| mOPC_1FA-REP3 | 2141631 | 1950400 | 191231 |

**Suppl. Fig. 14. Frequency of REs in RADICL-seq libraries from mESCs and mOPCs.** Proportion of significant RNA-DNA interactions in RADICL-seq mESCs and mOPCs libraries with RNA tags intersecting or non-intersecting RepeatMasker elements. The table displays the number of interactions shown in the plot.

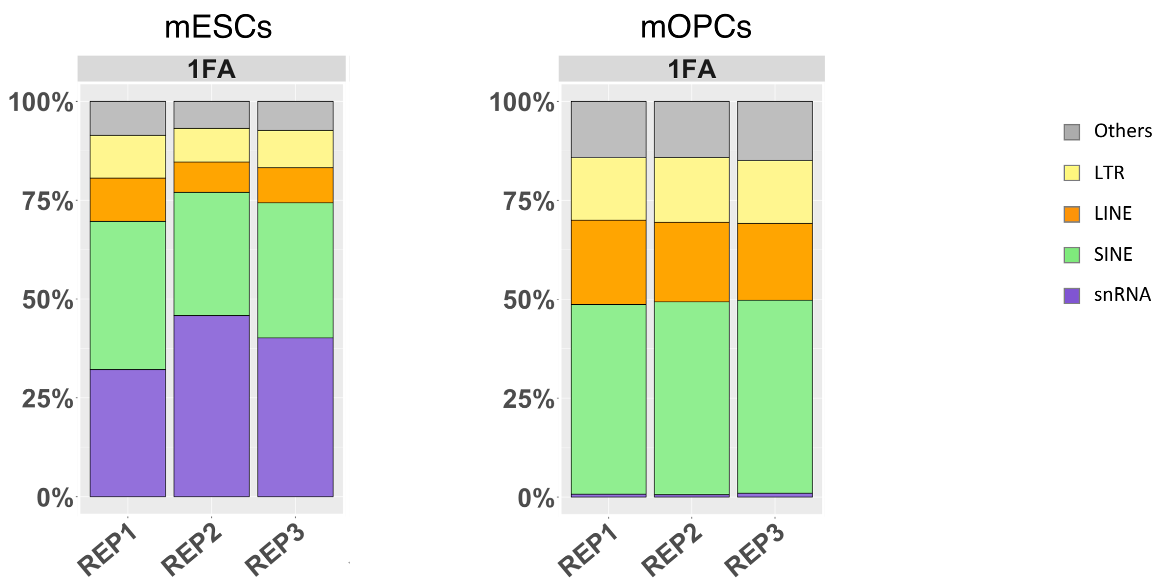


| Breakdown of Repeat Families for RNA-DNA pairs with RNA intersecting Repeat Masker elements annotated within GENCODE vM14 | | | | | | |
|---|---|---|---|---|---|---|
| **Condition** | **Total** | **snRNA** | **SINE** | **LINE** | **LTR** | **Others** |
| mESC_1FA-REP1 | 264627 | 85071 | 99254 | 28945 | 28542 | 22815 |
| mESC_1FA-REP2 | 285197 | 130571 | 88986 | 21821 | 24298 | 19521 |
| mESC_1FA-REP3 | 309959 | 124454 | 105955 | 27486 | 29215 | 22849 |
| mOPC_1FA-REP1 | 264627 | 922 | 99254 | 28945 | 28542 | 22815 |
| mOPC_1FA-REP2 | 285197 | 811 | 88986 | 21821 | 24298 | 19521 |
| mOPC_1FA-REP3 | 309959 | 1202 | 105955 | 27486 | 29215 | 22849 |

**Suppl. Fig. 15. Distribution of RE families in RADICL-seq libraries from mESCs and mOPCs.** Representation of main repeats families for RNA-DNA pairs where the RNA intersect with RepeatMasker elements contained within GENCODE vM14 annotations for RADICL-seq mESCs and mOPCs libraries. snRNA for GRID-seq replicas were <1% and therefore values were not plotted. Only non-self interactions were used for this plot. The table displays the count of the interactions used for the plot.

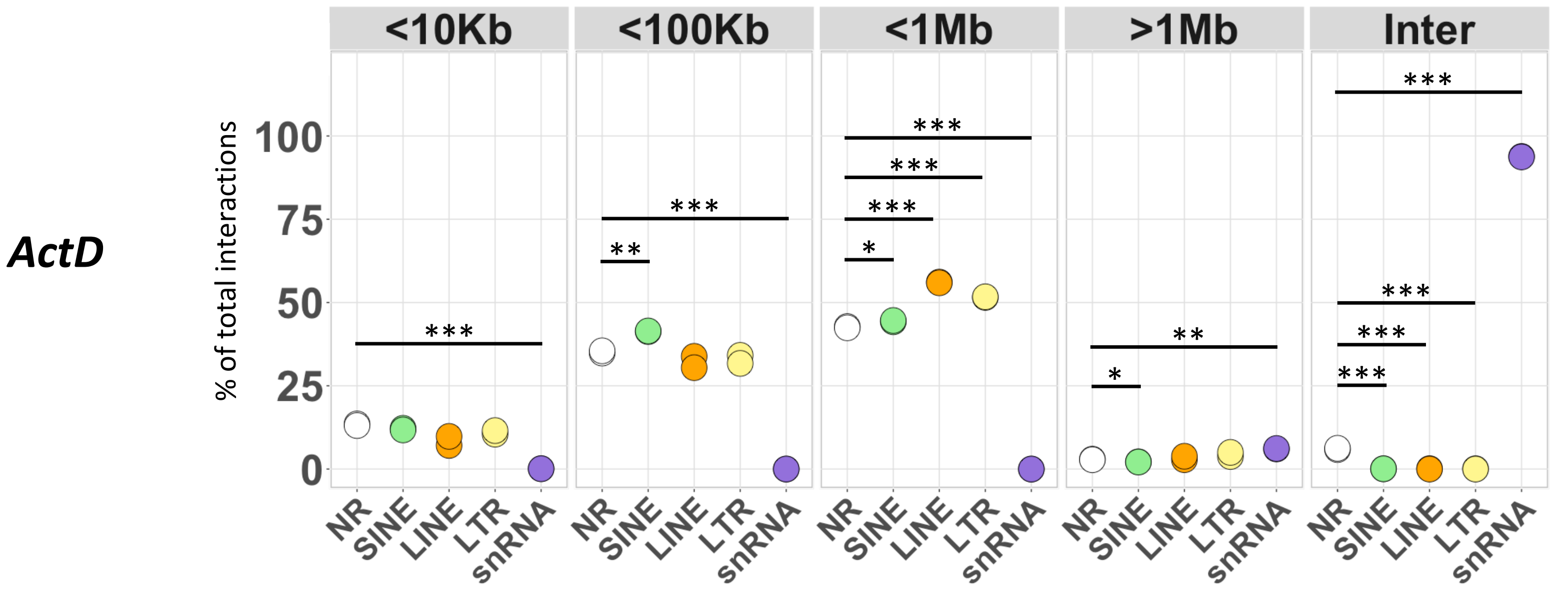


**Suppl. Fig. 16. Patterns of RNA-DNA distance interval for RE-mediated interactions in ActD condition.** Distribution of RNA-DNA pairs distance intervals for RNA tags intersecting RepeatMasker annotations paired with DNA tags intersecting GENCODE annotations in RADICL-seq mESCs libraries upon ActD treatment. Only RNA tags intersecting RepeatMasker elements contained within GENCODE vM14 features were used for this analysis. Statistical significance was calculated with two-tailed t-test. * = p≤0.05; ** = p≤0.01; *** = p≤0.001

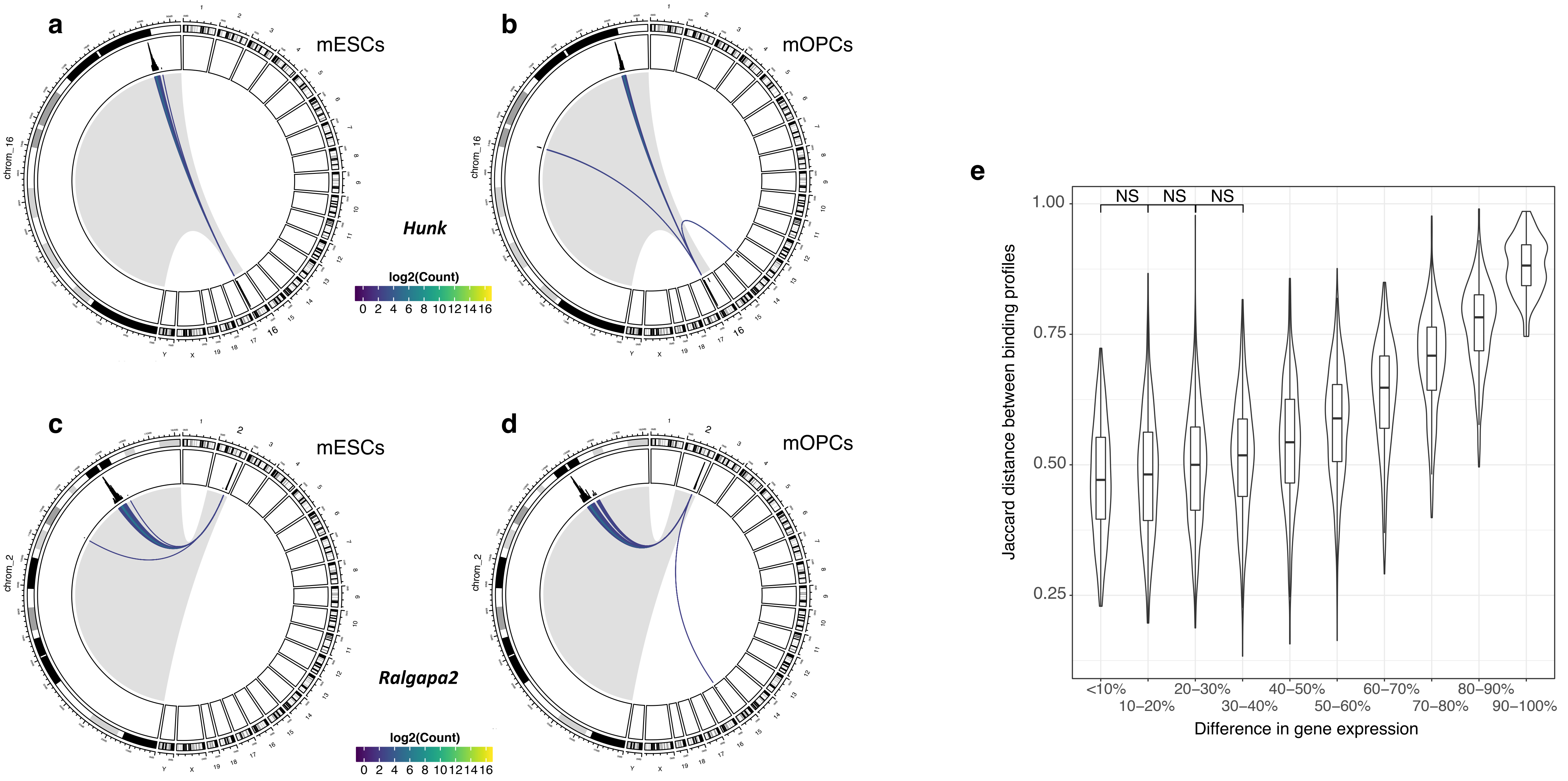


**Suppl. Fig. 17. Cell type-specific patterns of interactions.** a-b) Circos plots depicting Hunk genomic interactions in mESC and mOPC. c-d) Circos plots depicting Ralgapa2 genomic interactions in mESC and mOPC. Each line represents the interaction between the RNA gene and the contacted DNA locus, while its color highlight the occurrence of the association. The chromosome of origin of the RNA under investigation is shown enlarged on the left portion of each circos plot. e) The distribution of Jaccard distances for genome-wide RNA-DNA binding profiles between cell types for deciles of difference in gene expression between cell types. For genes with a difference in gene expression between cell types < 40%, there is no significant difference in the distribution of Jaccard distances for genome-wide RNA-DNA binding profiles.

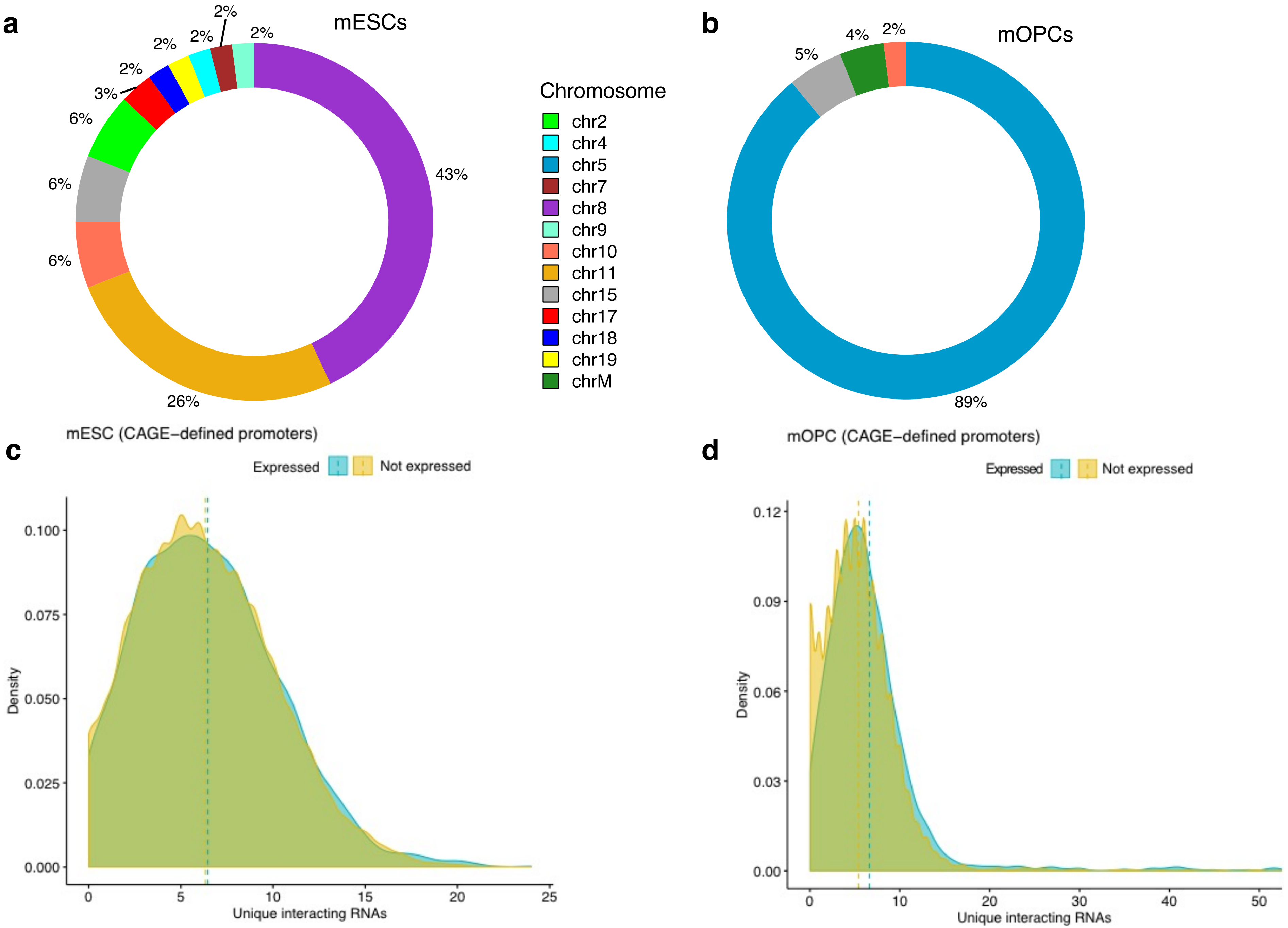


**Suppl. Fig. 18. RNA-DNA interactions at promoter regions.** a-b) Distribution of CAGE-derived promoters (top 100 ranked by number of unique interacting RNAs) per chromosome in a) mESCs and b) mOPCs. c) Distribution of unique interacting RNAs on expressed (>=3TPM; yellow) and not expressed (<3TPM; blue) CAGE-derived promoters (+/- 2 kb) in mESCs and d) mOPCs.

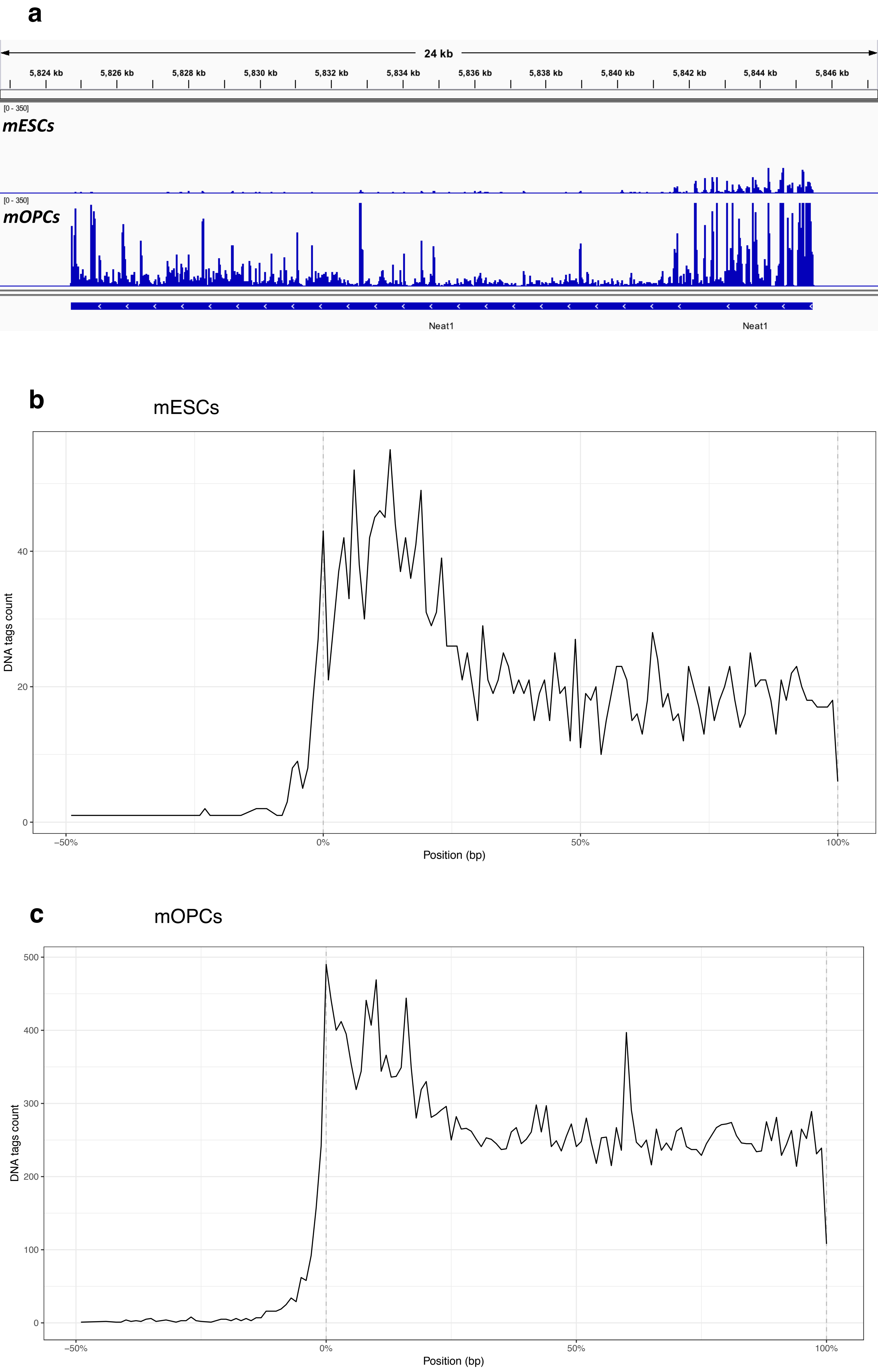


**Suppl. Fig. 19. Characterization of Neat1-mediated RNA-chromatin interactions.** a) Genomic view of the density of Neat1 RNA tags across its gene body in 1% FA mOPCs (top) and mESCs (bottom). b-c) Density of Neat1 mESCs and mOPCs target genes DNA tags across their gene bodies. Position is reported as percentage of the gene body length.

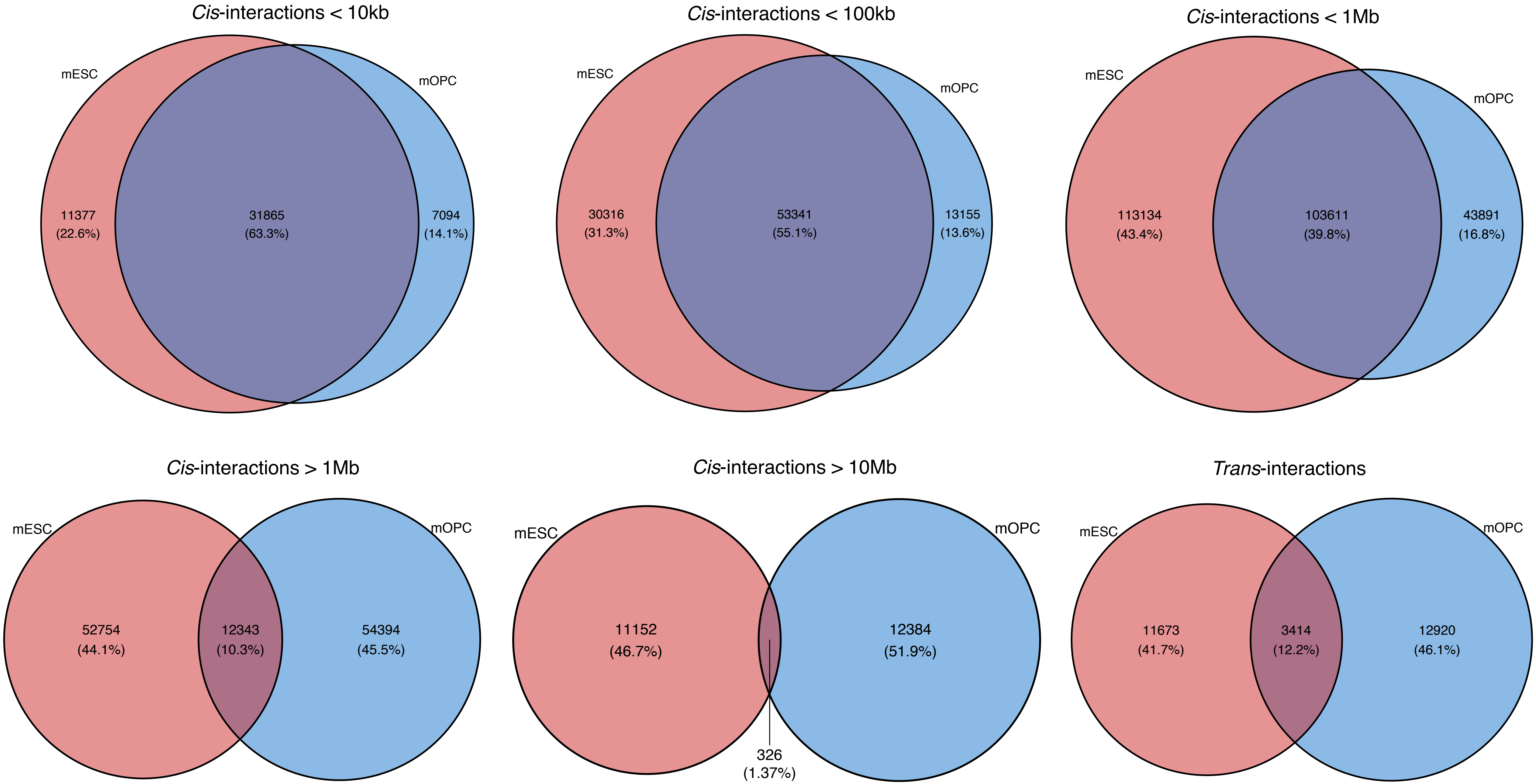


**Suppl. Fig. 20. Comparison of mESCs and mOPCs *cis*- and *trans*-interactions.** Interactions overlaps between mESC (red) and mOPC (blue) at less than 10kb, 100kb, 1Mb and more than 1Mb, 10Mb in linear distance between RNA and DNA tags or interacting in *trans*.

| Condition | Distance interval | Unique lncRNAs | Interactions | Percentage |
|---|---|---|---|---|
| mESCs 1FA | 1bp – 10 kb | 1,422 | 40,952 | 12.8% |
| mESCs 1FA | 10 kb – 100 kb | 1,177 | 52,280 | 16.4% |
| mESCs 1FA | 100 kb – 1 Mb | 285 | 60,331 | 18.9% |
| mESCs 1FA | 1 Mb – 10 Mb | 40 | 23,090 | 7.2% |
| mESCs 1FA | >10 Mb | 5 | 9,267 | 2.9% |
| mESCs 1FA | Trans | 3 | 133,705 | 41.8% |
| mOPCs 1FA | 1bp – 10 kb | 1074 | 58,781 | 5.4% |
| mOPCs 1FA | 10 kb – 100 kb | 958 | 68,932 | 6.4% |
| mOPCs 1FA | 100 kb – 1 Mb | 250 | 91,052 | 8.5% |
| mOPCs 1FA | 1 Mb – 10 Mb | 62 | 66,539 | 6.1% |
| mOPCs 1FA | >10 Mb | 17 | 38,277 | 3.5% |
| mOPCs 1FA | Trans | 10 | 759,844 | 70.1% |

**Suppl. Table 1. Analysis of lncRNA distance interaction patterns.** For each pair of RNA and DNA distance interval the number of unique lncRNAs and the number and fraction of all lncRNA-mediated interactions are reported for both mESCs and mOPCs datasets.

**Supplementary Table 2. Top ten interacting transcripts for cis- and trans-contacts.**
Only lncRNAs and protein-coding transcripts are listed for each condition.

| Experiment type | ENSG | Gene Name | #Contacts |
|---|---|---|---|
| RADICL_mESC_1FA | ENSMUSG00000092341.2 | Malat1 | 134436 |
| RADICL_mESC_1FA | ENSMUSG00000106943.1 | Dancr | 104 |
| RADICL_mESC_1FA | ENSMUSG00000092274.2 | Neat1 | 57 |
| RADICL_mESC_1FA | ENSMUSG00000097451.10 | Rian | 15 |
| RADICL_mESC_1FA | ENSMUSG00000021268.17 | Meg3 | 12 |
| RADICL_mESC_1FA | ENSMUSG00000101609.1 | Kcnq1ot1 | 12 |
| RADICL_mESC_1FA | ENSMUSG00000108414.1 | Snhg1 | 9 |
| RADICL_mESC_1FA | ENSMUSG00000078952.9 | Lncenc1 | 7 |
| RADICL_mESC_1FA | ENSMUSG00000085396.7 | Firre | 6 |
| RADICL_mESC_1FA | ENSMUSG00000097391.8 | Mirg | 3 |
| RADICL_mESC_4h_ActD | ENSMUSG00000092341.2 | Malat1 | 9277 |
| RADICL_mESC_4h_ActD | ENSMUSG00000106943.1 | Dancr | 9 |
| RADICL_mESC_4h_ActD | ENSMUSG00000097451.10 | Rian | 5 |
| RADICL_mESC_NPM | ENSMUSG00000092341.2 | Malat1 | 2206 |
| RADICL_mESC_NPM | ENSMUSG00000092274.2 | Neat1 | 58 |
| RADICL_mESC_NPM | ENSMUSG00000021268.17 | Meg3 | 41 |
| RADICL_mESC_NPM | ENSMUSG00000101609.1 | Kcnq1ot1 | 38 |
| RADICL_mESC_NPM | ENSMUSG00000097451.10 | Rian | 13 |
| RADICL_mESC_NPM | ENSMUSG00000078952.9 | Lncenc1 | 13 |
| RADICL_mESC_NPM | ENSMUSG00000085438.1 | 1700020I14Rik | 8 |
| RADICL_mESC_NPM | ENSMUSG00000097391.8 | Mirg | 6 |
| RADICL_mESC_NPM | ENSMUSG00000085715.2 | Tsix | 5 |
| RADICL_mESC_NPM | ENSMUSG00000085396.7 | Firre | 5 |
| RADICL_mOPC_1FA | ENSMUSG00000092341.2 | Malat1 | 735180 |
| RADICL_mOPC_1FA | ENSMUSG00000092274.2 | Neat1 | 24374 |
| RADICL_mOPC_1FA | ENSMUSG00000106943.1 | Dancr | 173 |
| RADICL_mOPC_1FA | ENSMUSG00000073147.3 | 5031425E22Rik | 16 |
| RADICL_mOPC_1FA | ENSMUSG00000087259.7 | 2610035D17Rik | 13 |
| RADICL_mOPC_1FA | ENSMUSG00000101609.1 | Kcnq1ot1 | 12 |
| RADICL_mOPC_1FA | ENSMUSG00000097039.8 | Pvt1 | 12 |
| RADICL_mOPC_1FA | ENSMUSG00000097536.2 | 2610037D02Rik | 8 |
| RADICL_mOPC_1FA | ENSMUSG00000097814.5 | Panct2 | 2 |
| RADICL_mOPC_1FA | ENSMUSG00000075555.11 | Gm10863 | 1 |
| RADICL_mOPC_NPM | ENSMUSG00000092341.2 | Malat1 | 76184 |
| RADICL_mOPC_NPM | ENSMUSG00000092274.2 | Neat1 | 12781 |
| RADICL_mOPC_NPM | ENSMUSG00000101609.1 | Kcnq1ot1 | 47 |
| RADICL_mOPC_NPM | ENSMUSG00000097039.8 | Pvt1 | 36 |
| RADICL_mOPC_NPM | ENSMUSG00000073147.3 | 5031425E22Rik | 24 |
| RADICL_mOPC_NPM | ENSMUSG00000087259.7 | 2610035D17Rik | 22 |
| RADICL_mOPC_NPM | ENSMUSG00000098243.3 | Gm4258 | 4 |
| RADICL_mOPC_NPM | ENSMUSG00000097207.7 | 6030443J06Rik | 3 |
| RADICL_mOPC_NPM | ENSMUSG00000105265.4 | Sox2ot | 3 |
| RADICL_mOPC_NPM | ENSMUSG00000054556.6 | Gm4876 | 2 |

pcRNA_interchrom

| Experiment type | ENSG | Gene Name | #Contacts |
|---|---|---|---|
| RADICL_mESC_1FA | ENSMUSG00000062647.16 | Rpl7a | 18 |
| RADICL_mESC_1FA | ENSMUSG00000027404.15 | Snrpb | 14 |
| RADICL_mESC_1FA | ENSMUSG00000036427.5 | Gpi1 | 9 |
| RADICL_mESC_1FA | ENSMUSG00000004980.16 | Hnrnpa2b1 | 5 |
| RADICL_mESC_1FA | ENSMUSG00000031939.16 | Taf1d | 5 |
| RADICL_mESC_1FA | ENSMUSG00000078578.9 | Ube2d3 | 5 |
| RADICL_mESC_1FA | ENSMUSG00000000131.15 | Xpo6 | 4 |
| RADICL_mESC_1FA | ENSMUSG00000021767.16 | Kat6b | 4 |
| RADICL_mESC_1FA | ENSMUSG00000022884.14 | Eif4a2 | 4 |
| RADICL_mESC_1FA | ENSMUSG00000030275.6 | Etnk1 | 4 |
| RADICL_mESC_NPM | ENSMUSG00000010608.15 | Rbm25 | 14 |
| RADICL_mESC_NPM | ENSMUSG00000021196.14 | Pfkp | 14 |
| RADICL_mESC_NPM | ENSMUSG00000025964.15 | Adam23 | 12 |
| RADICL_mESC_NPM | ENSMUSG00000021767.16 | Kat6b | 11 |
| RADICL_mESC_NPM | ENSMUSG00000022884.14 | Eif4a2 | 10 |
| RADICL_mESC_NPM | ENSMUSG00000030275.6 | Etnk1 | 9 |
| RADICL_mESC_NPM | ENSMUSG00000035569.17 | Ankrd11 | 9 |
| RADICL_mESC_NPM | ENSMUSG00000004980.16 | Hnrnpa2b1 | 8 |
| RADICL_mESC_NPM | ENSMUSG00000029817.11 | Tra2a | 8 |
| RADICL_mESC_NPM | ENSMUSG00000032582.14 | Rbm6 | 8 |
| RADICL_mOPC_1FA | ENSMUSG00000026872.17 | Zeb2 | 22 |
| RADICL_mOPC_1FA | ENSMUSG00000023951.17 | Vegfa | 13 |
| RADICL_mOPC_1FA | ENSMUSG00000027404.15 | Snrpb | 11 |
| RADICL_mOPC_1FA | ENSMUSG00000031939.16 | Taf1d | 11 |
| RADICL_mOPC_1FA | ENSMUSG00000051910.13 | Sox6 | 11 |
| RADICL_mOPC_1FA | ENSMUSG00000062647.16 | Rpl7a | 10 |
| RADICL_mOPC_1FA | ENSMUSG00000039630.10 | Hnrnpu | 6 |
| RADICL_mOPC_1FA | ENSMUSG00000068748.7 | Ptprz1 | 6 |
| RADICL_mOPC_1FA | ENSMUSG00000005103.12 | Wdr1 | 5 |
| RADICL_mOPC_1FA | ENSMUSG00000029673.17 | Auts2 | 4 |
| RADICL_mOPC_NPM | ENSMUSG00000051910.13 | Sox6 | 15 |
| RADICL_mOPC_NPM | ENSMUSG00000026872.17 | Zeb2 | 9 |
| RADICL_mOPC_NPM | ENSMUSG00000005103.12 | Wdr1 | 8 |
| RADICL_mOPC_NPM | ENSMUSG00000056073.16 | Grik2 | 7 |
| RADICL_mOPC_NPM | ENSMUSG00000053477.16 | Tcf4 | 7 |
| RADICL_mOPC_NPM | ENSMUSG00000004642.13 | Slbp | 6 |
| RADICL_mOPC_NPM | ENSMUSG00000026615.14 | Eprs | 5 |
| RADICL_mOPC_NPM | ENSMUSG00000059921.15 | Unc5c | 5 |
| RADICL_mOPC_NPM | ENSMUSG00000068748.7 | Ptprz1 | 4 |
| RADICL_mOPC_NPM | ENSMUSG00000004980.16 | Hnrnpa2b1 | 3 |

| Experiment type | ENSG | Gene Name | #Contacts |
|---|---|---|---|
| RADICL_mESC_1FA | ENSMUSG00000101609.1 | Kcnq1ot1 | 9410 |
| RADICL_mESC_1FA | ENSMUSG00000092341.2 | Malat1 | 9363 |
| RADICL_mESC_1FA | ENSMUSG00000092274.2 | Neat1 | 2186 |
| RADICL_mESC_1FA | ENSMUSG00000021268.17 | Meg3 | 1628 |
| RADICL_mESC_1FA | ENSMUSG00000078952.9 | Lncenc1 | 812 |
| RADICL_mESC_1FA | ENSMUSG00000085385.7 | Snhg17 | 536 |
| RADICL_mESC_1FA | ENSMUSG00000110282.1 | B930086L07Rik | 536 |
| RADICL_mESC_1FA | ENSMUSG00000097451.10 | Rian | 515 |
| RADICL_mESC_1FA | ENSMUSG00000097039.8 | Pvt1 | 483 |
| RADICL_mESC_1FA | ENSMUSG00000085715.2 | Tsix | 474 |
| RADICL_mESC_4h_ActD | ENSMUSG00000092341.2 | Malat1 | 489 |
| RADICL_mESC_4h_ActD | ENSMUSG00000101609.1 | Kcnq1ot1 | 171 |
| RADICL_mESC_4h_ActD | ENSMUSG00000096751.3 | Gm28373 | 86 |
| RADICL_mESC_4h_ActD | ENSMUSG00000021268.17 | Meg3 | 84 |
| RADICL_mESC_4h_ActD | ENSMUSG00000110558.1 | Gm45779 | 52 |
| RADICL_mESC_4h_ActD | ENSMUSG00000078952.9 | Lncenc1 | 39 |
| RADICL_mESC_4h_ActD | ENSMUSG00000097000.2 | Gm17435 | 38 |
| RADICL_mESC_4h_ActD | ENSMUSG00000092274.2 | Neat1 | 31 |
| RADICL_mESC_4h_ActD | ENSMUSG00000100691.1 | 2010320M18Rik | 22 |
| RADICL_mESC_4h_ActD | ENSMUSG00000097039.8 | Pvt1 | 20 |
| RADICL_mESC_Control | ENSMUSG00000092203.7 | 1110038B12Rik | 20 |
| RADICL_mESC_NPM | ENSMUSG00000105260.1 | Gm42434 | 16 |
| RADICL_mESC_NPM | ENSMUSG00000092341.2 | Malat1 | 9 |
| RADICL_mESC_NPM | ENSMUSG00000110282.1 | B930086L07Rik | 8 |
| RADICL_mESC_NPM | ENSMUSG00000111394.1 | AC160637.1 | 6 |
| RADICL_mESC_NPM | ENSMUSG00000079489.2 | C030013D06Rik | 5 |
| RADICL_mESC_NPM | ENSMUSG00000102657.1 | Gm37899 | 5 |
| RADICL_mESC_NPM | ENSMUSG00000110993.1 | Gm47963 | 5 |
| RADICL_mESC_NPM | ENSMUSG00000070461.3 | 9230112E08Rik | 4 |
| RADICL_mESC_NPM | ENSMUSG00000102411.1 | Gm36936 | 4 |
| RADICL_mOPC_1FA | ENSMUSG00000092341.2 | Malat1 | 26738 |
| RADICL_mOPC_1FA | ENSMUSG00000092274.2 | Neat1 | 9033 |
| RADICL_mOPC_1FA | ENSMUSG00000101609.1 | Kcnq1ot1 | 8420 |
| RADICL_mOPC_1FA | ENSMUSG00000073147.3 | 5031425E22Rik | 2155 |
| RADICL_mOPC_1FA | ENSMUSG00000097039.8 | Pvt1 | 936 |
| RADICL_mOPC_1FA | ENSMUSG00000107197.1 | Gm43312 | 738 |
| RADICL_mOPC_1FA | ENSMUSG00000107083.1 | Gm43313 | 664 |
| RADICL_mOPC_1FA | ENSMUSG00000070461.3 | 9230112E08Rik | 532 |
| RADICL_mOPC_1FA | ENSMUSG00000097536.2 | 2610037D02Rik | 441 |
| RADICL_mOPC_1FA | ENSMUSG00000104118.1 | Gm37298 | 310 |
| RADICL_mOPC_NPM | ENSMUSG00000092341.2 | Malat1 | 552 |
| RADICL_mOPC_NPM | ENSMUSG00000092274.2 | Neat1 | 174 |
| RADICL_mOPC_NPM | ENSMUSG00000073147.3 | 5031425E22Rik | 45 |
| RADICL_mOPC_NPM | ENSMUSG00000094832.1 | Gm21846 | 26 |
| RADICL_mOPC_NPM | ENSMUSG00000103038.1 | Gm37124 | 22 |
| RADICL_mOPC_NPM | ENSMUSG00000103427.1 | Gm37534 | 20 |
| RADICL_mOPC_NPM | ENSMUSG00000070461.3 | BX649560.1 | 16 |
| RADICL_mOPC_NPM | ENSMUSG00000107197.1 | Gm43312 | 16 |
| RADICL_mOPC_NPM | ENSMUSG00000085334.7 | Gm12940 | 14 |
| RADICL_mOPC_NPM | ENSMUSG00000111212.1 | CAAA01194877.1 | 13 |

pcRNA_intrachrom

| Experiment type | ENSG | Gene Name | #Contacts |
|---|---|---|---|
| RADICL_mESC_1FA | ENSMUSG00000021767.16 | Kat6b | 863 |
| RADICL_mESC_1FA | ENSMUSG00000035569.17 | Ankrd11 | 815 |
| RADICL_mESC_1FA | ENSMUSG00000078578.9 | Ube2d3 | 814 |
| RADICL_mESC_1FA | ENSMUSG00000025964.15 | Adam23 | 799 |
| RADICL_mESC_1FA | ENSMUSG00000022995.16 | Enah | 724 |
| RADICL_mESC_1FA | ENSMUSG00000031575.18 | Ash2l | 645 |
| RADICL_mESC_1FA | ENSMUSG00000038518.15 | Jarid2 | 616 |
| RADICL_mESC_1FA | ENSMUSG00000055320.17 | Tead1 | 594 |
| RADICL_mESC_1FA | ENSMUSG00000029178.14 | Klf3 | 570 |
| RADICL_mESC_1FA | ENSMUSG00000040952.16 | Rps19 | 520 |
| RADICL_mESC_4h_ActD | ENSMUSG00000033732.10 | Sf3b3 | 92 |
| RADICL_mESC_4h_ActD | ENSMUSG00000029817.11 | Tra2a | 64 |
| RADICL_mESC_4h_ActD | ENSMUSG00000035569.17 | Ankrd11 | 55 |
| RADICL_mESC_4h_ActD | ENSMUSG00000029439.14 | Sfswap | 50 |
| RADICL_mESC_4h_ActD | ENSMUSG00000037957.14 | Wdr20 | 49 |
| RADICL_mESC_4h_ActD | ENSMUSG00000038095.15 | Sbno1 | 49 |
| RADICL_mESC_4h_ActD | ENSMUSG00000034269.12 | Setd5 | 48 |
| RADICL_mESC_4h_ActD | ENSMUSG00000024576.14 | Csnk1a1 | 47 |
| RADICL_mESC_4h_ActD | ENSMUSG00000059796.16 | Eif4a1 | 44 |
| RADICL_mESC_4h_ActD | ENSMUSG00000040731.13 | Eif4h | 42 |
| RADICL_mESC_Control | ENSMUSG00000071172.12 | Srsf3 | 5 |
| RADICL_mESC_Control | ENSMUSG00000023944.14 | Hsp90ab1 | 4 |
| RADICL_mESC_Control | ENSMUSG00000024073.14 | Birc6 | 4 |
| RADICL_mESC_Control | ENSMUSG00000024163.17 | Mapk8ip3 | 3 |
| RADICL_mESC_Control | ENSMUSG00000028461.12 | Ccdc107 | 3 |
| RADICL_mESC_Control | ENSMUSG00000036026.15 | Tmem63b | 3 |
| RADICL_mESC_Control | ENSMUSG00000036036.15 | Zfp57 | 3 |
| RADICL_mESC_Control | ENSMUSG00000039218.16 | Srrm2 | 3 |
| RADICL_mESC_Control | ENSMUSG00000023991.16 | Foxp4 | 2 |
| RADICL_mESC_Control | ENSMUSG00000024459.18 | H2-M5 | 2 |
| RADICL_mOPC_1FA | ENSMUSG00000026872.17 | Zeb2 | 2262 |
| RADICL_mOPC_1FA | ENSMUSG00000005103.12 | Wdr1 | 1592 |
| RADICL_mOPC_1FA | ENSMUSG00000014956.15 | Ppp1cb | 1160 |
| RADICL_mOPC_1FA | ENSMUSG00000032525.15 | Nktr | 722 |
| RADICL_mOPC_1FA | ENSMUSG00000037235.13 | Mxd4 | 711 |
| RADICL_mOPC_1FA | ENSMUSG00000032911.6 | Cspg4 | 624 |
| RADICL_mOPC_1FA | ENSMUSG00000053007.9 | Creb5 | 491 |
| RADICL_mOPC_1FA | ENSMUSG00000059921.15 | Unc5c | 481 |
| RADICL_mOPC_1FA | ENSMUSG00000053477.16 | Tcf4 | 469 |
| RADICL_mOPC_1FA | ENSMUSG00000053007.9 | Creb5 | 453 |
| RADICL_mOPC_Control | ENSMUSG00000029673.17 | Auts2 | 138 |
| RADICL_mOPC_Control | ENSMUSG00000029138.4 | 4930548H24Rik | 138 |
| RADICL_mOPC_Control | ENSMUSG00000079555.2 | Haus3 | 144 |
| RADICL_mOPC_Control | ENSMUSG00000037605.16 | Adgrl3 | 141 |
| RADICL_mOPC_Control | ENSMUSG00000052139.18 | Bre | 176 |
| RADICL_mOPC_Control | ENSMUSG00000015452.14 | Ager | 407 |
| RADICL_mOPC_Control | ENSMUSG00000020882.17 | Cacnb1 | 96 |
| RADICL_mOPC_Control | ENSMUSG00000025138.14 | Sirt7 | 180 |
| RADICL_mOPC_Control | ENSMUSG00000028035.13 | Dnajb4 | 180 |
| RADICL_mOPC_Control | ENSMUSG00000033102.15 | Cdc14b | 264 |

mESCs markers, list modified from Zhao, W. et al. *Molecules* **17**, 6196-6246 (2012).

| Gene name | Gene ID |
|---|---|
| Cd324 | ENSMUSG00000000303.12 |
| Nacc1 | ENSMUSG00000001910.4 |
| Klf4 | ENSMUSG00000003032.8 |
| Stat3 | ENSMUSG00000004040.16 |
| Cd117 | ENSMUSG00000005672.12 |
| Nanog | ENSMUSG00000012396.12 |
| Tbx3 | ENSMUSG00000018604.18 |
| Essrb | ENSMUSG00000021255.17 |
| Pou5f1 | ENSMUSG00000024406.16 |
| Tfcp2l1 | ENSMUSG00000026380.10 |
| Sall4 | ENSMUSG00000027547.17 |
| Lef1 | ENSMUSG00000027985.14 |
| Cd90 | ENSMUSG00000032011.5 |
| Cripto | ENSMUSG00000032494.12 |
| Fbxo15 | ENSMUSG00000034391.10 |
| Gbx2 | ENSMUSG00000034486.8 |
| Cd326 | ENSMUSG00000045394.8 |
| Dppa3 | ENSMUSG00000046323.8 |
| Utf1 | ENSMUSG00000047751.9 |
| Ssea-1 | ENSMUSG00000049307.6 |
| Rex1 | ENSMUSG00000051176.6 |
| Klf2 | ENSMUSG00000055148.7 |
| Hmga2 | ENSMUSG00000056758.14 |
| Dppa4 | ENSMUSG00000058550.14 |
| Dppa5 | ENSMUSG00000060461.5 |
| Gcnf | ENSMUSG00000063972.13 |
| Foxd3 | ENSMUSG00000067261.4 |
| Dppa2 | ENSMUSG00000072419.4 |
| Sox2 | ENSMUSG00000074637.7 |
| Zfx | ENSMUSG00000079509.10 |

mOPCs markers, list modified from Marques S. et al. Science 352, 1326-1329 (2016).

| Gene name | Gene id |
|---|---|
| Gria3 | ENSMUSG00000001986.16 |
| Bcan | ENSMUSG00000004892.13 |
| Slc1a2 | ENSMUSG00000005089.15 |
| Atp1a2 | ENSMUSG00000007097.14 |
| Tnr | ENSMUSG00000015829.13 |
| Matn4 | ENSMUSG00000016995.17 |
| Fabp7 | ENSMUSG00000019874.11 |
| Ascl1 | ENSMUSG00000020052.9 |
| Id2 | ENSMUSG00000020644.9 |
| Zfp36l1 | ENSMUSG00000021127.7 |
| Vcan | ENSMUSG00000021614.16 |
| Ednrb | ENSMUSG00000022122.14 |
| Serpine2 | ENSMUSG00000026249.10 |
| Gpr37l1 | ENSMUSG00000026424.8 |
| Pdgfra | ENSMUSG00000029231.15 |
| Tmem176b | ENSMUSG00000029810.15 |
| Tpm1 | ENSMUSG00000032366.15 |
| Cspg5 | ENSMUSG00000032482.9 |
| Cspg4 | ENSMUSG00000032911.6 |
| Sox10 | ENSMUSG00000033006.9 |
| Neu4 | ENSMUSG00000034000.15 |

| | |
|---|---|
| Rlbp1 | ENSMUSG00000039194.15 |
| Olig2 | ENSMUSG00000039830.8 |
| Tril | ENSMUSG00000043496.7 |
| S100a1 | ENSMUSG00000044080.9 |
| C1ql1 | ENSMUSG00000045532.5 |
| Olig1 | ENSMUSG00000046160.6 |
| Ncald | ENSMUSG00000051359.14 |
| Sox6 | ENSMUSG00000051910.13 |
| Gpr17 | ENSMUSG00000052229.5 |
| Pcdh15 | ENSMUSG00000052613.16 |
| Sox11 | ENSMUSG00000063632.6 |
| Nnat | ENSMUSG00000067786.16 |
| Ptprz1 | ENSMUSG00000068748.7 |
| Tmem100 | ENSMUSG00000069763.3 |
| Ccnd1 | ENSMUSG00000070348.5 |
| Kcnip3 | ENSMUSG00000079056.12 |
| Lhfpl3 | ENSMUSG00000106379.1 |

**Suppl. Table 3. List of cell type-specific markers.**

| Transcript ID | Name | mESC (1FA) | mOPC (1FA) |
|---|---|---|---|
| ENSMUSG00000092341.2 | Malat1 | 86,770 | 209,954 |
| ENSMUSG00000064899.1 | Snord118 | 294 | 103 |
| ENSMUSG00000088088.1 | Rmrp | 112 | 106 |
| ENSMUSG00000092274.2 | Neat1 | 15 | 48 |
| ENSMUSG00000064337.1 | Mt-Rnr1 | 15 | 36 |
| ENSMUSG00000023795.16 | Pisd-ps2 | 14 | 23 |
| ENSMUSG00000097039.8 | Pvt1 | 14 | 18 |
| ENSMUSG00000064339.1 | Mt-Rnr2 | 13 | 13 |
| ENSMUSG00000065037.1 | Rn7sk | 12 | 20 |
| ENSMUSG00000026131.18 | Dst | 9 | 11 |
| ENSMUSG00000065878.1 | Snord34 | 8 | 21 |
| ENSMUSG00000077734.1 | Snord83b | 8 | 4 |
| ENSMUSG00000048264.16 | Dip2c | 7 | 7 |
| ENSMUSG00000104627.1 | Mir3535 | 4 | 6 |

**Suppl. Table 4. Shared *trans*-interacting long noncoding RNAs captured in both mESCs and mOPCs.** For each transcript, the numbers of shared interactions in each cell type are shown.
